# Understanding the complexity of Epimorphic Regeneration in zebrafish: A Transcriptomic and Proteomic approach

**DOI:** 10.1101/2021.05.20.445069

**Authors:** Sarena Banu, Namami Gaur, Sowmya Nair, Tanuja Ravikrishnan, Shahida Khan, Sandhya Mani, Swarna Bharathi, Komal Mandal, Naga Anusha Kuram, Sowmya Vuppaladadium, Ch. Lakshmi N. Murthy, Mir Quoseena, Nukala Sarath Babu, Mohammed M Idris

## Abstract

Genomic and Proteomic changes play a crucial role in perpetuating regeneration of complex tissues through differentiation and growth. The complex Epimorphic regeneration of zebrafish caudal fin tissue is hasty and absolute. This study was executed to understand the role of various genes/proteins involved in the regeneration of zebrafish caudal fin tissue through differential expression analysis. High throughput transcriptomics analysis involving Next Generation Sequencing approach and iTRAQ based quantitative proteomics analyses were performed on the regenerating tissue samples for various regenerating time points. Based on our study 1408 genes and 661 proteins were found differentially regulated in the regenerating caudal fin tissue for having at least 1-log fold change in their expression at 12hpa, 1, 2, 3 and 7dpa stages against control non-regenerating tissue. Interleukin, SLC, PRMT, HOX, neurotransmitter and several novel genes were found to be associated with regeneration for its differential regulation during the mechanism. Based on the network and pathway analysis the differentially regulated genes and proteins were found allied with activation of cell proliferation, cell viability, cell survival & cell movement and inactivation of organismal death, morbidity, necrosis, death of embryo & cell death. Network pathways such as Cancer & development disorder, Cell signaling molecular transport, organismal injury & abnormalities and Cellular development, growth & proliferation were found most significantly associated with the zebrafish caudal fin regeneration mechanism. This study has mapped a detailed insight of the genes/proteins expression associated with the epimorphic regeneration more profoundly.

## Introduction

Regeneration is a remarkable biological process which restores cellular architecture and function of an injured tissue by virtue of a complex mechanism. Blastema-mediated epimorphic regeneration is an exquisitely synchronized proliferation and patterning of cells observed in many organisms to restore damaged and amputated organs. The process of regeneration in higher vertebrates like mammals is still enigmatic, whereas it is a prevalent feature among the lower vertebrates like teleost fish and amphibians like salamander, newts and *Xenopus* ^(1).^Understanding the regeneration mechanism is necessary to revamp and benefit the regeneration deficient organisms.

Syntenic correlation of human with zebrafish proves it to be a unique vertebrate model for human disease ^(2)^. Zebrafish possess the ability to regenerate heart, appendages, optic nerve and spinal cord throughout life by epimorphic regeneration ^(3,4)^. Quick regenerative ability, easy access, and simpler anatomy furnish the caudal fin of zebrafish as an exemplary system to comprehend vertebrate organ regeneration. Zebrafish caudal fin regeneration comprises of four stages: “epithelialization or wound healing” (0–1day post-amputation (dpa)), “blastema formation” (1–2 dpa), “regenerative outgrowth” (2–7 dpa), and “termination” ^(5)^. At 1 dpa, a drift of the proximal epidermis is seen that covers the amputated region, followed by inflammation. Histolysis, cell dedifferentiation and blastema formation starts at 2dpa. Subsequently, between 3-7 dpa, regenerative outgrowth is seen with robust proliferation of dedifferentiated cells, progressive redifferentiation and morphogenesis takes place. After 8-10 dpa, the fin regenerates completely and regenerative homeostasis is achieved ^(3,5)^.

Previous studies have deciphered the key roles of various genes and associated pathways concerning regeneration in lower vertebrates. Retinoic acid (RA) pathway is one such key pathway that balances fibroblast growth factor (FGF), Wnt /β-catenin and fgf signaling during blastema formation in caudal fin of zebrafish ^(6)^. The bone morphogenetic protein (BMP) and FGFwhich primarily are involved in organ growth and proliferation, are also involved in regeneration ^(7)^. Another significant signaling pathway in tissue regeneration is Wnt signaling pathway which majorly regulates blastema proliferation and redifferentiation ^(8)^. Juxtacrine-Notch signaling pathways have also been linked to fin and heart regeneration of zebrafish ^(9)^. The biomolecular characterization of Msx homeobox gene family has demonstrated that these genes are highly expressed during blastema formation post fin amputation ^(10)^. Cytokeratins are well known intermediate filament proteins involved in cellular differentiation and wound healing. Keratin 5 and keratin 8 have shown to be linked to regeneration of caudal fin and its human orthologues are considered as regeneration epidermal markers ^(11, 12)^. Several studies validate the substantial role of ECM (Extra Cellular Matrix) during tissue development, regeneration, morphogenesis and wound healing. Biomolecular cues have shown that Hyaluronic acid, Fibronectin and Tenascin C are highly upregulated during fin regeneration ^(13)^. Moreover, blastemal proliferation and rate of growth post injury is tightly controlled by FGF signaling pathway which helps in maintaining homeostatic balance ^(14)^. In sum, regeneration is a pool of different cellular and molecular mechanisms to harmonize the signals that induce cell proliferation and those that halt cell division and maintains homeostasis ^(15)^.In this work, we analyzed the global transcriptomics and proteomics changes during the zebrafish caudal fin tissue regeneration for different regenerating time points post amputation.

## Results

### Regeneration of Caudal fin tissue

Amputation of zebrafish caudal fin tissue leads to the regeneration of the fin tissue following wound healing and physical growth (Figure 1). The regeneration of the fin is first marked by the formation of a wound epidermis and later on followed by the blastema (Figure 1b – 1d). The blastema is very fragile and much care was taken while handling the animals without disrupting it. The regeneration growth was monitored until 7dpa. The formation of bony rays or lepidotrichia was visible from 2 to 3 dpa (Figure 1e – 1f) and the actinotrichia were seen visible on the 7dpa stage at the proximal-distal axis (Figure 1g). 70% of its lost tissue was found regenerated by the end of 7dpa (Figure 1i). The rate of regeneration was found rapid in the lobe part of the fin tissue with a mean growth of 0.91 cms tissue at the end of 7dpa, whereas in the cleft region of the fin tissue the growth rate was 0.49 cm at the end of 7dpa (Figure 1i).

**Figure 1:**
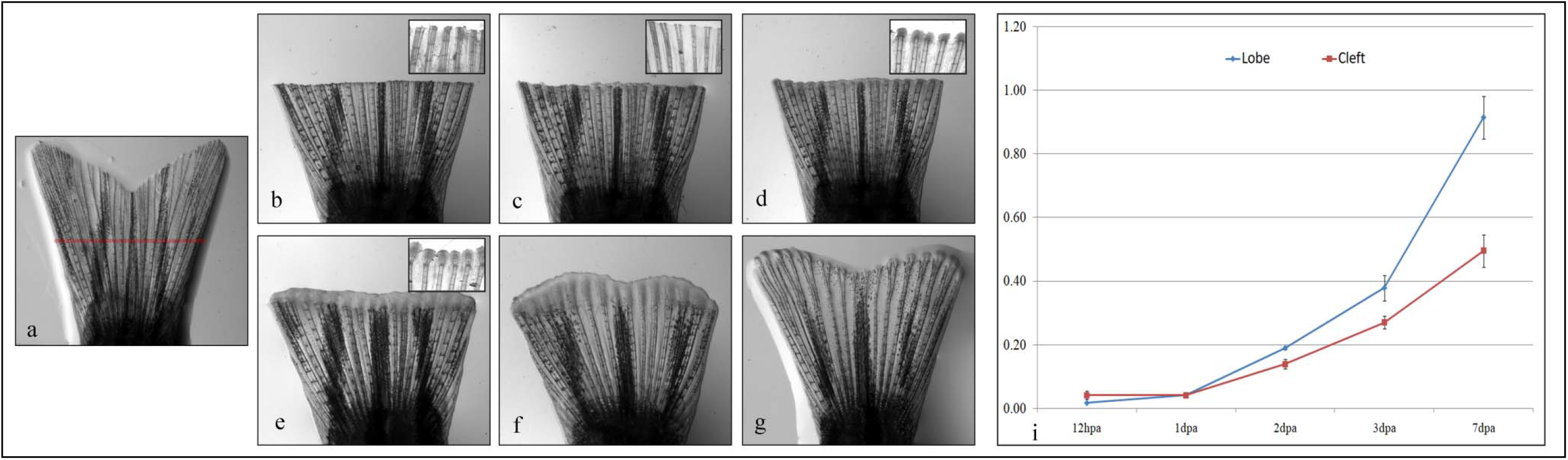
Regeneration of zebrafish caudal fin tissue post amputation a. Uncut zebrafish caudal fin; b. Amputated zebrafish fin at 0hpa; c. Regenerated caudal fin at 12hpa; d. Regenerated caudal fin at 1dpa; e. Regenerated caudal fin at 2dpa; f. Regenerated caudal fin at 3dpa; g. Regenerated caudal fin at 7dpa; i. Regenerative growth index of zebrafish caudal fin in the lobe and cleft region for different regenerating time points.

### Differential Transcriptomics analysis

The transcriptome sequence obtained from NGS data for 0hpa, 12hpa, 1dpa, 2pa, 3pa and 7dpa were submitted to NCBI database and obtained PRJNA248169 BioProject accession number. Based on differential next generation sequencing analysis a total of 1408 genes were found to be differentially expressed during the regeneration of caudal fin tissue in zebrafish (Figure 2a). The spectrum of changes in the expression of different genes was analyzed among controls (0hpa) and 12hpa, 1dpa, 2dpa, 3dpa and 7dpa. The genes were differentially expressed by at least one log fold change in one of the selected regenerating time points. Theprominent number of genes and their familieswere found to be associated with regeneration through differential regulation which includestranscription factors (TFs), protein arginine methyltransferases (PRMTs), solute career genes (SLC genes), neurotransmitter genes, Interleukin family genes, homeobox genes and several novel genes.NRSN1, RCOR2, RASA4 and ANAPC4 were a few of the most up regulated genes and TMEM9, RHOBTB2, EML1 and ednrb1a were a few of the most down regulated genes associated with the regeneration of zebrafish caudal fin tissue based on differential NGS analysis (Supplementary Table 1). Based on heat map analysis of the gene expression it was found that 12hpa and 1dpa showed similar expression pattern rooting with 2dpa, which was out rooted with 3 and 7dpa (Figure 2a).

**Figure 2:**
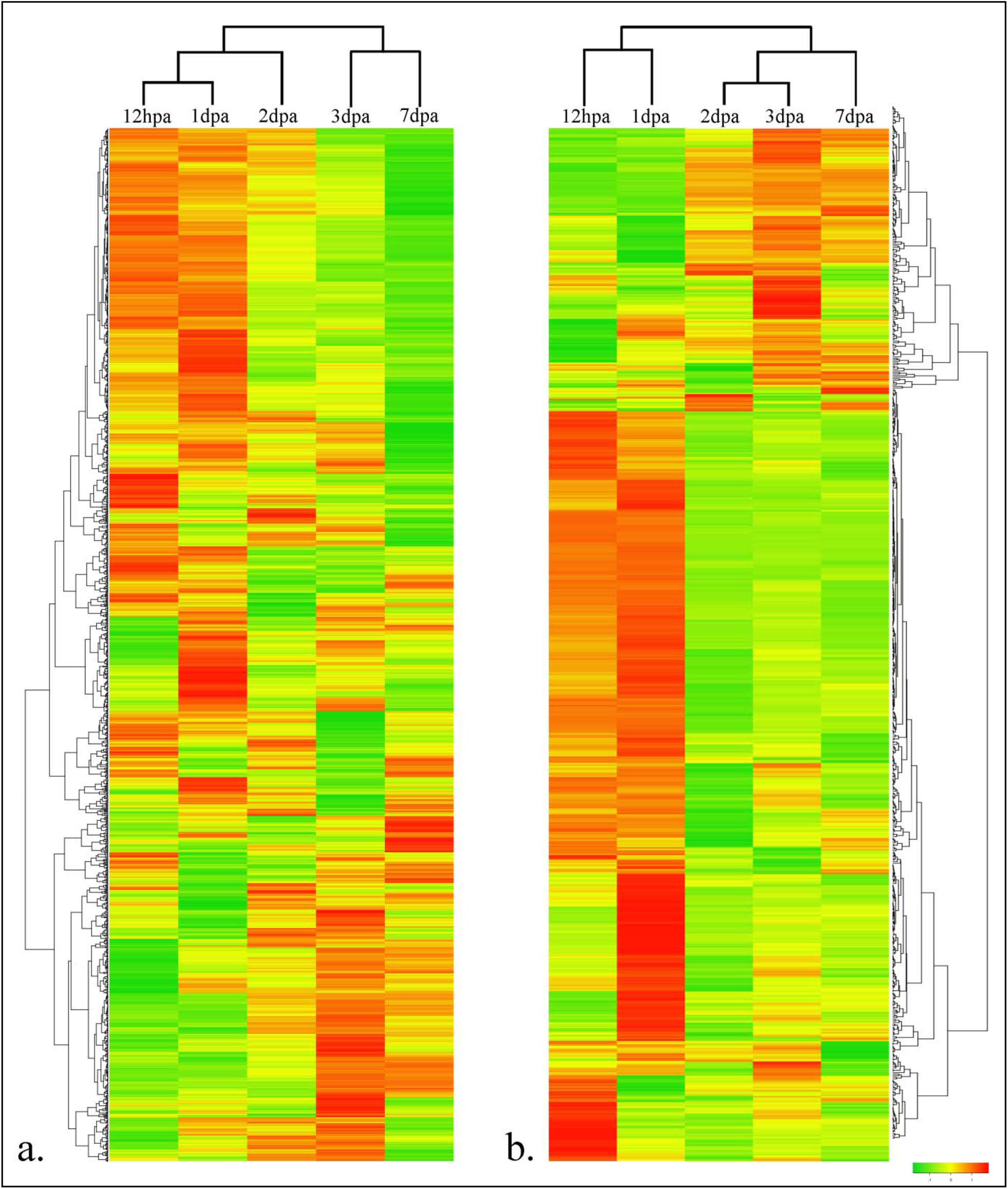
Heat map expression associated with zebrafish caudal fin regeneration for different regenerating time points, a. Transcriptome Heat map and b. Proteome Heat map expression at 12hpa, 1,2,3 and 7dpa stages.

### Differential Proteomic analysis

A Total of 661 proteins were found to be associated with the caudal fin tissue regeneration based on comparative proteomics analysis for having at least one log fold changes in one of the selected time points (Figure 2b, Supplementary Table 2). HECW1, GIMAP8, and RTF1 were the most up regulated and BRPF3, PKP4 and MCF2 were the most down regulated proteins associated with regeneration of the caudal fin tissue at the protein level. Collagen, dedicator of cytokinesis, DNA polymerase and replication, Dynein, E3 ubiquitin, interferon, keratin, kinesin, myosin, neuronal and several hypothetic family proteins were found to be associated with the regeneration mechanism (Supplementary Table 2). At protein level, 12hpa and 1dpa showed similar protein expression pattern, which out rooted with 2, 3 and 7dpa expressions found based on heat map analysis (Figure 2b)

### Validation of Gene expression

A total of 227 genes were selected for the validation of gene expression involving qRT-PCR analysis. The gene list includes 140 genes from the transcriptomics analysis, 28 interleukin family genes, 16 SLC family genes, 9 PRMT genes, 13 HOX family genes and 21 neurotransmitter genes (Supplementary Table 3, Figure 3). The genes were differentially expressed in the regenerating tissue either immediately after amputation (12hpa) or later during the later time points of regeneration. Adam8a, fn1b, krt18, lepb, mmp13a, orc3, orc6, psmb7, txn and zgc 153629 were few of the genes identified from NGS study and were found to be most significantly up regulated for all the regenerating time points (Figure 3a). Similarly, the most significantly down regulated genes from NGS studies for all the regenerating time points includes papss2a, eif4ebp3, pax7b, kcnj1a, ptgdsb, ppp1r3cb, syt4 and zgc163079 (Figure 3a).

**Figure 3:**
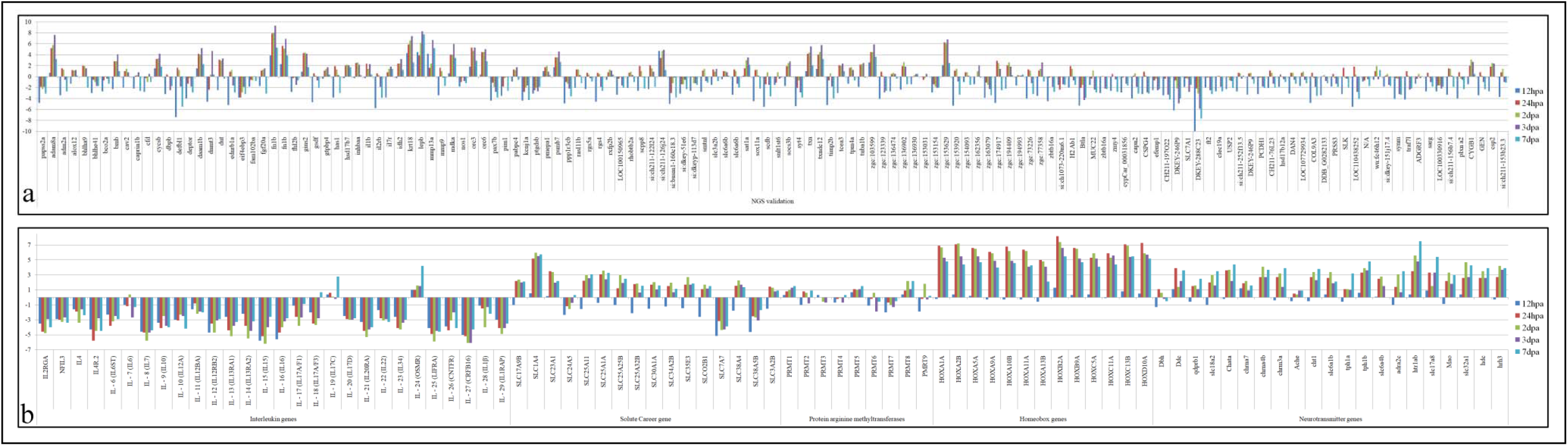
RTPCR expression analysis of genes associated with zebrafish caudal fin regeneration for different regenerating time poitns. a. Genes selected from the NGS analysis, b. List of genes associated as families.

Gene expression analysis of 28 Interleukin family genes showed a spectrum of down regulation for all the selected interleukin gene members except IL19 and IL24. The down regulation of Interleukin genes spread from 12hpa to 7dpa (Figure 3b). Solute Career family genes showed a down regulation at 12hpa followed by up regulation for the rest of the selected regenerating time points except for SLC24A5, SLC7A7 and SLC38A5B which were found to be down regulated for all the regenerating time points (Figure 3b). Theprotein arginine methyltransferase (PRMT) family gene showed a subtle differential gene expression pattern for the selected regenerating time points (Figure 3b). Homeobox genes showed a colossal up regulation of the gene expression from 1dpa onwards for all the selected HOX genes (Figure 3b). An overall pattern of up regulation was observed from 1dpa for all the selected neurotransmitter genes during regeneration of the zebrafish caudal fin tissue (Figure 3b).

### Cytokine Array

Expression of all the 34 cytokine proteins in the antibody array was found expressed in the caudal fin tissue at 0, 1, 2 and 3dpa (Figure 4a). Differential expression analysis of the cytokine protein based on antibody array showed an absolute up regulation of all the 34 selected cytokines at 1dpa. But the expressions of the cytokines were found down regulated during the later regenerating time points such as 2 and 3dpa (Figure 4b). Fas Ligand, Fractalkine, IL13, IL1R6, IL2 and IL4 were the few selected cytokines found up regulated extensively at 1dpa in comparison to other cytokines. IL13, VEGF and TNF-α were a few of the cytokines found extremely down regulated at 2 and 3dpa (Figure 4b).

**Figure 4:**
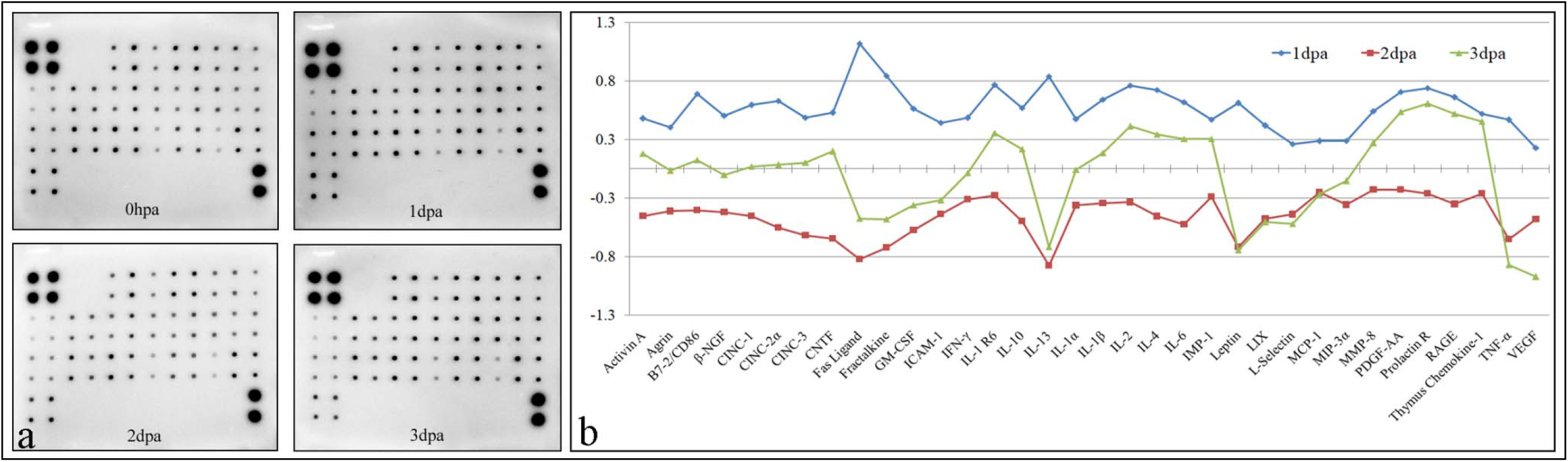
Cytokine protein array expression associated with zebrafish caudal fin regeneration. a. Immuno Dot blot image of cytokine array at 0hpa, 1dpa, 2dpa and 3dpa. b. Differential expression of cytokine protein at 1dpa, 2dpa and 3dpa against control, 0hpa.

### Network and Pathway analysis

A total of 1702 genes/proteins were mapped from the list of 2122 genes/proteins obtained from the transcriptomics, proteomics and gene expression analysis for the network and pathway analysis. The most significantly up regulated canonical pathway from the list of genes/proteins includes Cell cycle, Cell Cycle Control of Chromosomal Replication, NER (Nucleotide Excision Repair, Enhanced Pathway), Oxidative Phosphorylation, Spliceosomal Cycle and Role of BRCA1 in DNA Damage Response pathways. Similarly, the most significantly down regulated canonical pathways for the regeneration mechanism includes ErbB signaling, Huntington’s Disease Signaling, AMPK Signaling, Regulation of The Epithelial-Mesenchymal Transition by Growth Factors Pathway, Growth Factors Pathway and IL9 signaling (Figure 5). The most significantly up regulated disease and functions from the list of differentially expressed genes/proteins includes Cell viability of tumor cell lines, cell survival, cell movement, migration of cell and cell viability. Organismal death, morbidity, necrosis, death of embryo and cell death were found to be down regulated in the regenerating caudal fin tissue (Figure 5).

**Figure 5:**
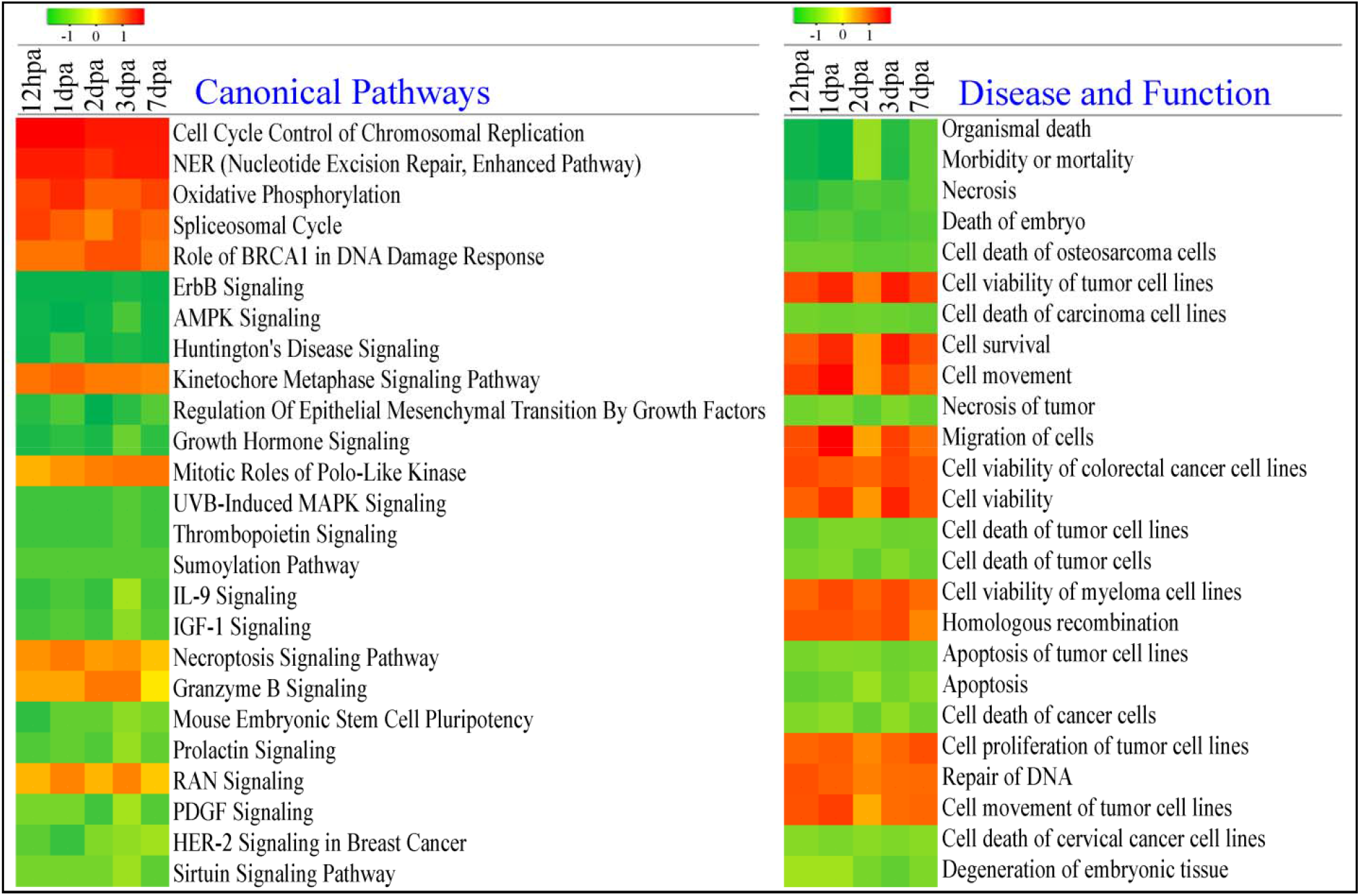
Canonical Pathways and Disease & Functions associated with the differentially expressed genes/proteins based on network and pathway analysis for zebrafish caudal fin regeneration.

Activation of cell proliferation, invasion of cells & cell movement and inhibition of formation of nuclear foci and SMARCB1 were found mapped at 12hpa from the list of genes/proteins (Figure 6a). At 1dpa G1/S phase transition, colony formation, cycling of centrosome and proliferation of fibroblast cell line were found activated (Figure 6b). Neoplasia of cells, cell proliferation of carcinoma cell lines, cell proliferation of breast cancer cell line and cell proliferation of tumor cell lines were activated at 2dpa (Figure 6c). At 3 and 7dpa Invasion of tumor cell lines, cell movement of tumor cell line and colony formation of cells was found activated (Figure 5d and Figure 6e) based on network pathway analysis.

**Figure 6:**
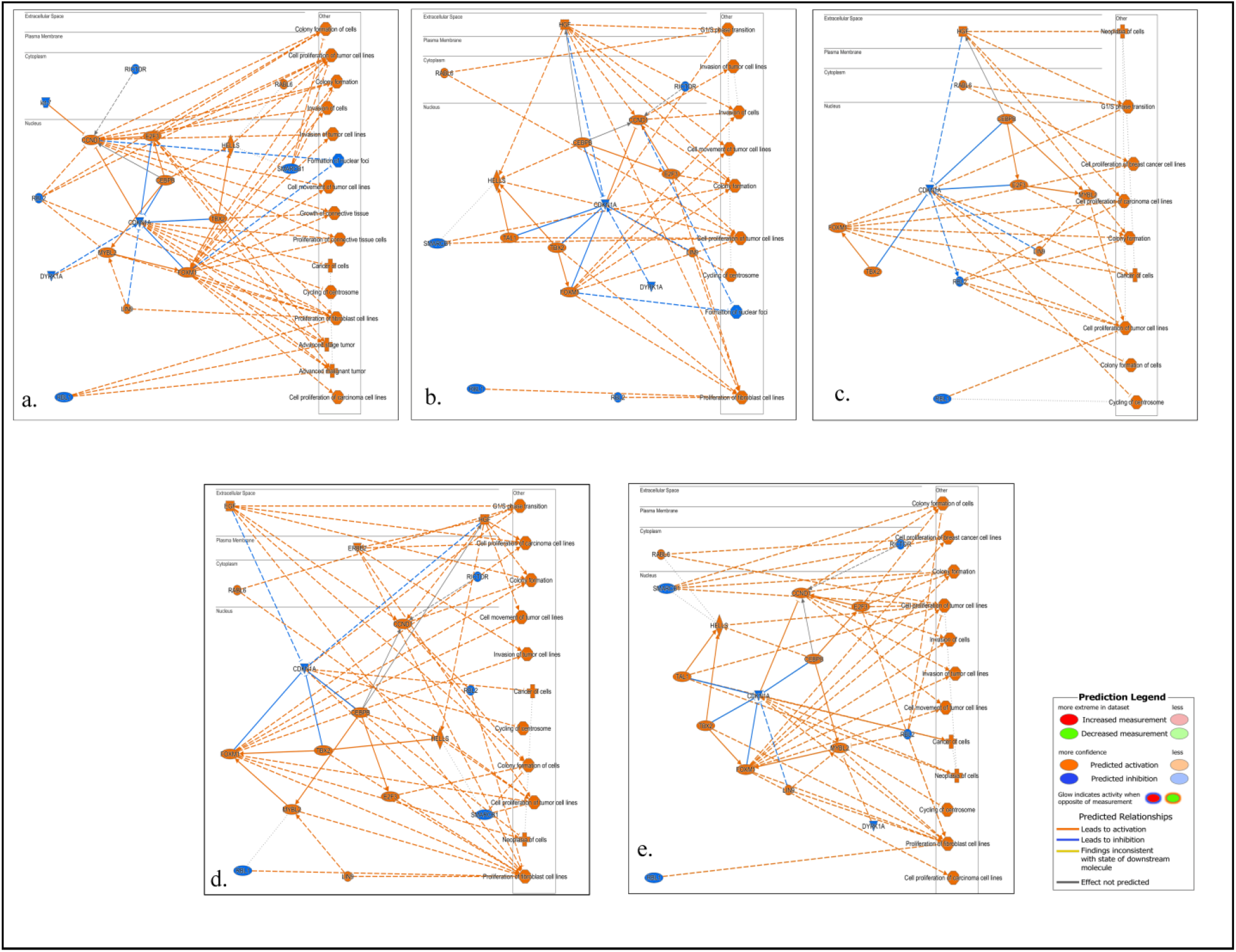
Various networks associated with differentially expressed genes/proteins for zebrafish caudal fin regeneration at a. 12hpa; b. 1dpa; c. 2dpa; d. 3dpa and e. 7dpa.

The most significantly associated network pathways from the list of differentially regulated genes/proteins includes Cancer and development disorder pathway involving 131 genes/proteins, Cell signaling molecular transport pathway involving 123 genes/proteins, Organismal injury and abnormalities involving 122 genes/proteins and Cellular development, growth and proliferation pathway involving 110 genes/proteins (Figure 7). Up regulated genes/proteins such as SDF2, DTL, KCTN1, YWHAQ, NKD2, NDST4, HELLS, RNASEH2A, DNMT1, TANGO2, SMYD5, PRMT5, STIM2, SHMT1, PRMT1, LPIN1, DENND5B, SREBF2, SCAPER, ZNF271 and other genes/proteins were found associated with Cellular development, growth and proliferation pathway. Similarly, down regulated genes/proteins for this pathway includes TMEM9, ZBTB16, BRPF3, KCNJ8, SLC7A1, SULT1C1, CD74, SLCO2B1, MPZ, EPC1, and other genes/proteins (Figure 7).

**Figure 7:**
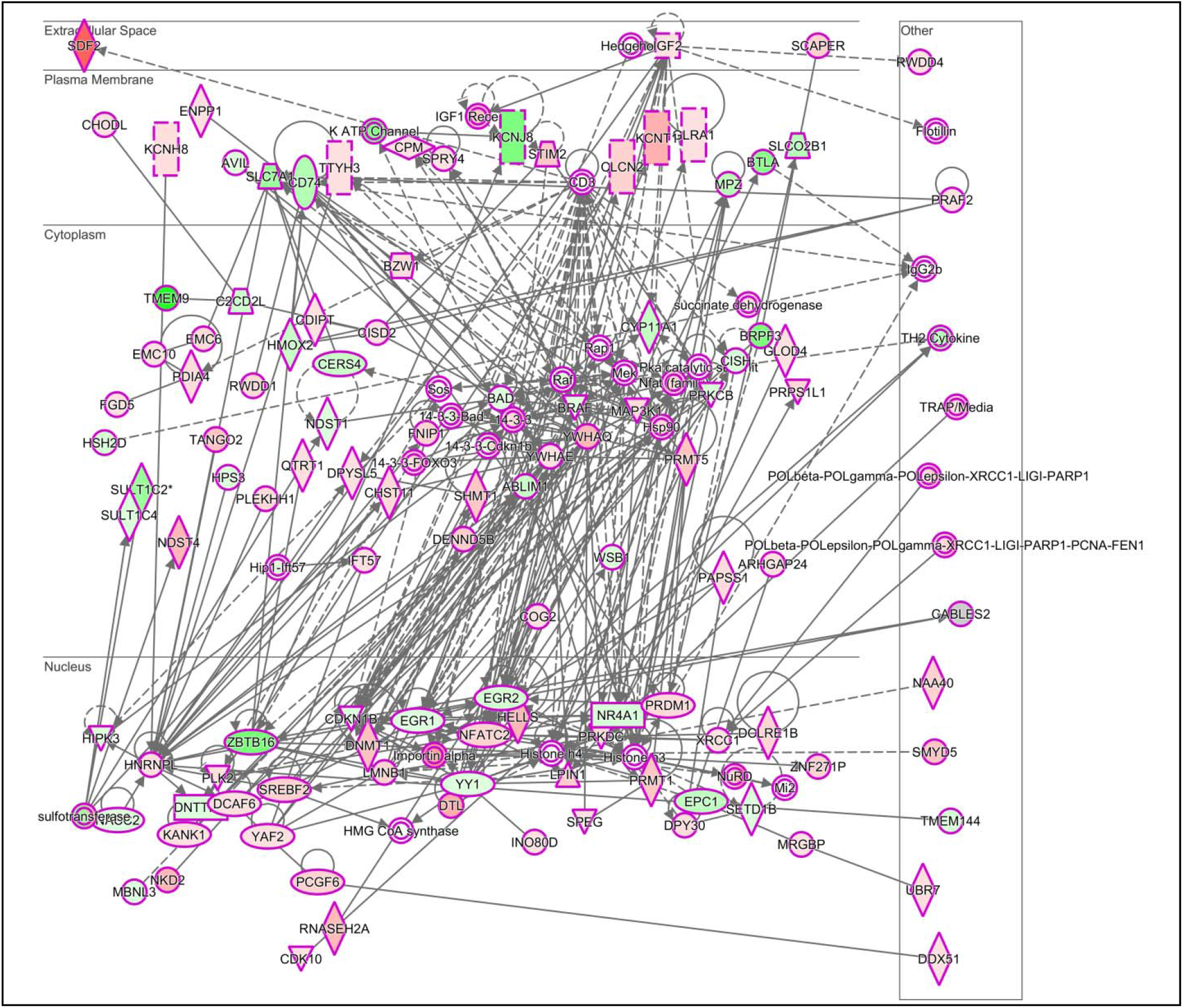
Cellular development, growth and proliferation pathway associated with zebrafish caudal fin regeneration based on network pathway analysis of differentially expressed genes/proteins.

## Discussion

In an attempt to unravel the complexity of epimorphic regeneration mechanism, this study executed a high throughput analysis of the genome and proteome changes stirring in the zebrafish caudal fin tissue post amputation for six different regenerating time points (Figure 1). Regeneration of zebrafish caudal fin tissue post amputation is rapid and discrete. At the end 7-day post amputation almost 70% of the amputated tissue were found regenerated at both lobe and cleft region of the caudal fin tissue. Understanding the differential expression of genes/proteins for these time points might shed complete information about the molecular mechanism of wound healing, proliferation, differentiation, dedifferentiation and structural regeneration stages. The study was performed for enlisting all the genes/proteins associated with caudal fin regeneration through differential regulation involving NGS and iTRAQ based proteomics analysis and gene expression analysis.

A total of 1408 genes and 661 proteins were found differentially expressed in all the regenerating time points based on NGS and comparative proteomics analysis. An up regulation of 18 log fold change (NRSN1 gene) to down regulation of 16 log fold change (TMEM9 gene) were observed at the gene level during the zebrafish caudal fin tissue regeneration (Supplementary Table 1). Similarly, a 6.6 log fold up regulation and 5.6 log fold down regulation were observed at protein level during the caudal fin tissue regeneration (Supplementary Table 2). Validation of 227 differentially expressed genes representing different gene families including interleukin, SLC, PRMT, HOX and Neurotransmitter showed a continuum of expression pattern during the regeneration mechanism.

Interleukin family genes showed a complete down regulation of gene expression right from wound healing to structural regeneration stages at gene level (Figure 3). But at protein level, the cytokines including few interleukins were found up regulated at 1dpa, i.e., at proliferation stage of regeneration, whereas at later stages, the cytokine proteins were found down regulated (Figure 4). Interleukins are a group of cytokines, classed by TIR domain (Toll-IL-1 resistance) and their biologic role exceeds the anticipated one in communication between the cells of the immune system during tissue damage^16, 17^. Proteins of interleukins (IL) family regulate cellular proliferation and it activates different signaling pathways, which can share intracellular signaling cascades e.g., MAPK, Ras or STAT^18^. During inflammation, infiltrate cytokines are dominant to remove cell debris and leads to tissue re-growth ^19, 20^. The prominent cytokine IL-1 is a pathogenic mediator of degenerative disease^21^. Il1b knockout wild type zebrafish illustrated that Il1b expression in epithelial cells initiates zebrafish fin fold regeneration^20^. Among cytokine family IL4 and IL13 found to be involved in TLR signaling pathway and IL10 assist in inflammatory reduction^22^.

Hox genes are transcriptional regulators that are involved in positional patterning in vertebrates during embryo development^23, 24^. These genes have the ability to re-express during regeneration^25^. HOX A, B, C, D clusters have expressed in fin development along with sonic hedgehog (Shh) signaling^23^. Hoxc13a and b knockdown showed retarded growth in caudal fin^25^. Our study showed a global up regulation of the HOX genes at gene level from 1dpa onwards (Figure 3).

Solute Carrier transporters are a super family of membrane-bound proteins that mediate the translocation of a wide spectrum of substrates and solutes across biological membranes by conferring selective membrane permeability^26, 27^. To date, 395 membrane-spanning SLC transporters have been distinguished all of which are categorized into 52 families respectively and they are prevalently settled at the cell surface and organelle membranes^27^. As membrane proteins, SLCs are supervised by ubiquitination^28^. SLC transporters are either electrochemical gradient or ion gradients^26^. Zebrafish golden mutants suggest that SLC24A5, a putative cation exchanger, has an established role in melanin pigmentation and melanosome biogenesis^29^. SLC1A4 significantly involved in melanocyte differentiation^30^. Our study showed an overall up regulation of the selected SLC gene from 1 to 7-day post amputation regenerating time points (Figure 3).

PRMT family showed disparate gene expression changes for different time points of regeneration stages. PRMT1, PRMT5 and PRMT8 were found up regulated for the regeneration of caudal fin tissue (Figure 3). PRMT family genes play a role in gene regulation through histone markers activation and repression^31–33^. PRMT1 in interferon (IFN) signaling leads to regulating the transcription induction of IFN^33^; PRMT2 promotes apoptosis^33^ and PRMT3 involved in ribosomal biosynthesis through methylating the 40S ribosomal protein S2^34^;CARM1 directly methylate p300^32,33^; PRMT5 methylates histones and act as growth control^32,35^. PRMT6 activate and repress transcription by methylating H4R3/H2AR3, H3R2^35, 36^. PRMT7 plays a role in transcriptional regulation, snRNP biogenesis, and splicing. PRMT7 depletion leads to increase sensitivity to topoisomerase II inhibitors^37, 38^. PRMT8 methylate H2A and H4 via N-terminal myristoylation and PRMT8 knockout fish has shown neural developmental defects^39^. PRMT9 is widely expressed and found in a variety of human tissues and associated with pigment loss in a chronic skin disease^35^.

The association of neurotransmitters during regeneration is evident from the up regulation of all the selected neurotransmitters. Several foldup regulations of neurotransmitter were observed at 1, 2, 3 and 7dpa (Figure 3). Neurotransmitters are involved in neuronal communication; it has a functional role in neuronal outgrowth during development. Up-regulated neurotransmitters during regeneration involved in the regenerative response, whereas down-regulated transmitterswere only responsible for transmission^40^. tph1b geneencoded tryptophan hydroxylase was found to be associated with blastema formation after the amputation of zebrafish caudal fin^41^.

Pathway analysis of the differentially expressed genes/proteins exhibited association of wide range of canonical pathways, disease and functions and network pathways. It was well documented from our earlier studies for association of several diverse network pathways such as cytoskeleton remodeling pathway^12,42^, Immune response based alternative complement pathway^12^,Integrin mediated cell adhesion and migration pathway^42^, VEGF signaling pathway^42^, cellular morphology and embryonic development network pathway^43^, cellular assembly and organization network pathway and organ & organismal development pathway^43^ for the regeneration of zebrafish caudal fin regeneration^12^, brittle star arm regeneration^42^ and lizard tail regeneration^43^.

This study has found association of similar pathways as identified before and also several other new pathways such as Cellular development, growth and proliferation pathway involving a highernumber of genes/proteins identified from the study. The wide range of association of pathways such as Cancer and development disorder pathway, Cell signaling molecular transport pathway, organismal injury and abnormalities pathway and, In canonical pathways such as cell cycle control of chromosomal replication, NER, ErbB signaling AMPK signaling and IL9 signaling with zebrafish caudal fin tissue regeneration is due to the high throughput approach executed in this study and also the identification of wide range of genes/proteins associated with the zebrafish caudal fin regeneration mechanism.

## Conclusion

This study of understanding the rapid and complex epimorphic regeneration of zebrafish caudal fin tissue involving high throughput NGS and proteomic analysis has identified the association of several hundreds of genes/proteins. Interleukin, SLC, PRMT, HOX and neurotransmitter family genes/proteins were identified to be associated with the regeneration mechanism. The identified differentially regulated genes/proteins were found to be associated with epimorphic regeneration of the zebrafish caudal fin tissue through several complex canonical pathways, disease, function and network pathways.

## Material and Methods

### 1. Animal Experiment

Adult zebrafish aged 6 to 12 months were selected for the study. Equal lengths of caudal fin tissues were amputated from the mid-fin to the distal region after brief anesthesia as described earlier ^12, 44^. For the experimental study, the regenerating caudal fin tissues were collected for different time points like 0, 12, 24, 48, 72, and 168 hours of post amputation in batches of 10 animals per group. The zebrafish animal experiments were performed as per the protocol approved by the institutional animal ethics committee of Center for Cellular and Molecular Biology (IAEC/CCMB/Protocol #50/2013).

### 2. Transcriptome analysis

Next generation sequencing analysis using Illumina HiSeq 2000 was performed for the total RNA extracted from each time point of regenerating tissue with biological duplicates^43, 42^. The obtained transcripts of all the tissues were assembled for *de novo* transcriptome analysis and subjected for functional annotation using Blastxalong with *Danio rerio* database. Furthermore, the gene sequences obtained through NGS analysis were submitted in the NCBI repository and their accession numbers were obtained ^42, 43^. Differential expressions of mRNA transcripts were analyzed by FKPM software having 0hpa transcripts ascontrol and those with more than 1 log fold changes at each time point of regeneration. ^42^.

### 3. Proteomic Analysis

Total protein of all time points of regenerating caudal fin tissues were extracted and quantified usingAmido black assay^12, 44, 43^with bovine serum albumin as standard.iTRAQ based quantitative proteomics was carried out for all the time points of regeneration against control (0hpa) in succession to a trypsin based 10% SDS gel digestion ^12, 43^. All the digested peptides were labelled withiTRAQ 4-plex and purified using C-18 spin columns (Thermo Scientific). Labelled and purified peptides were analysed using LC-MS/MS OrbitrapVelos Nano analyser (Thermo Scientific) ^12, 45^. The resulted raw data were analysed using Sequest HT proteome discoverer 1.4 (Thermo Scientific), with 1% FDR percolator and XCorr (Score Vs Charge) from the *Danio rerio*database ^43, 42^. Proteins having more than one log fold change in the selected time points were recruited for the study.

### 4. Validation of Gene expression

Real time PCR (RTPCR) analysis was performed in biological and technical replicates for the most significantly expressed families of genes such as PRMT, SLC, Interleukin, HOX and Neurotransmitter. Zebrafish ODC gene was used as the reference housekeeping gene and all the required primers were designed using Primer3 software. Reverse transcription PCR was performed using 2μg of RNA. Obtained cDNA was used as template for qPCR analysis with SYBR green reagent (TakaraBio). The Ct values for each time points from the qPCR, were checked for fold changes against control (0hpa) and for gene expression validation.

### 5. Antibody array

Cytokine expression analysis was performed using Cytokine Array - RAT Cytokine antibody array (Abcam, USA) kit. 300 μg of total proteins from control, 1, 2 and 3-dpa were immunoblotted in the kit as per manufacturer’s protocol. The obtained spot patterns were densitometrically analyzed using ImageJ software to estimate the expression level of cytokines during caudal fin regeneration.

### 6. Heat and Network Pathway Analysis

The differentially expressed genes/proteins were analyzed for the heat map and network pathway analysis involving heatmapper portal (www.heatmapper.ca) and Ingenuity pathway analysis (IPA) software respectively. The heat maps were generated separately for the differentially expressed genes and proteins involving hierarchical cluster analysis. Network pathways were generated using IPA software for the canonical pathway, disease and functions, graphical summary and gene network pathway.

## Acknowledgements

This work was supported by CSIR-YSA project. The Authors are thankful to Ms. Noorul Fowzia for critically reviewing the manuscript.

## Authors Contribution

SB, MQ, NG, KM, NSB, TR, SN, SB2, SK, NAK and LNM performed the experiment and analyzed the data; MMI –Designed the experiment and analyzed the data, MMI, SB and NG wrote the manuscript.

## Notes

### Competing Interest Statement

The authors have declared no competing interest.

