## Supplementary Table No. 1 for "Understanding the complexity of Epimorphic Regeneration in zebrafish: A Transcriptomic and Proteomic approach"

Supplementary Table 1: List of Genes differentially expressed based on NGS analysis.

| **S. No** | **Gene Name** | **Gene Symbol** | **12hpa** | **1dpa** | **2dpa** | **3dpa** | **7dpa** | **RT Validation** |
| --- | --- | --- | --- | --- | --- | --- | --- | --- |
| 1 | 3'-phosphoadenosine 5'-phosphosulfate synthase 2a | papss2a | -2.3 | -1.5 | -2.0 | -2.2 | -1.8 | Yes |
| 2 | a disintegrin and metalloproteinase domain 8a | adam8a | 5.1 | 5.3 | 6.4 | 6.5 | 4.8 | Yes |
| 3 | adrenomedullin 2a | adm2a | 3.8 | 3.5 | 3.6 | 3.8 | 3.3 | Yes |
| 4 | arachidonate 12-lipoxygenase, 12S type | alox12 | -1.9 | -2.0 | -2.8 | -2.1 | -1.9 | Yes |
| 5 | basic helix-loop-helix family member a9 | bhlha9 | 4.2 | 4.2 | 2.6 | 2.6 | 1.9 | Yes |
| 6 | basic helix-loop-helix family member e41 | bhlhe41 | -1.8 | -1.6 | -2.2 | -2.5 | -1.4 | Yes |
| 7 | beta-carotene oxygenase 2a | bco2a | -2.2 | -2.4 | -1.9 | -2.5 | -2.6 | Yes |
| 8 | brambleberry | bmb | 2.6 | 2.9 | 3.9 | 3.8 | 3.3 | Yes |
| 9 | caveolin 2 | cav2 | -1.9 | -1.5 | -2.0 | -1.8 | -1.7 | Yes |
| 10 | cell cycle associated protein 1b | caprin1b | -1.7 | -1.9 | -1.7 | -1.8 | -1.6 | Yes |
| 11 | complement factor D (adipsin) | cfd | -1.8 | -1.4 | -1.6 | -2.5 | -2.0 | Yes |
| 12 | cytochrome c, somatic b | cycsb | 2.6 | 2.1 | 1.7 | 1.5 | 1.4 | Yes |
| 13 | D site of albumin promoter (albumin D-box) binding protein b | dbpb | -4.0 | -2.0 | -2.5 | -2.4 | -2.2 | Yes |
| 14 | defensin, beta-like 1 | defbl1 | -2.7 | -2.9 | -3.0 | -2.9 | -4.5 | Yes |
| 15 | DEP domain containing MTOR-interacting protein | deptor | -3.1 | -2.0 | -2.1 | -2.2 | -2.0 | Yes |
| 16 | dishevelled associated activator of morphogenesis 1b | daam1b | 3.2 | 3.2 | 3.1 | 2.7 | 2.2 | Yes |
| 17 | DNA (cytosine-5-)-methyltransferase 3 | dnmt3 | 2.5 | 3.9 | 3.9 | 3.9 | 2.7 | Yes |
| 18 | dUTP pyrophosphatase | dut | 2.4 | 4.4 | 2.3 | 3.0 | 2.6 | Yes |
| 19 | endothelin receptor B1a | ednrb1a | -4.5 | -4.1 | -5.2 | -4.3 | -2.9 | Yes |
| 20 | eukaryotic translation initiation factor 4E binding protein 3 | eif4ebp3 | -1.5 | -2.4 | -1.9 | -1.8 | -2.4 | Yes |
| 21 | family with sequence similarity 102, member B, a | fam102ba | -2.2 | -1.6 | -2.0 | -1.6 | -1.3 | Yes |
| 22 | fibroblast growth factor 20a | fgf20a | 3.2 | 5.0 | 6.1 | 5.3 | 3.7 | Yes |
| 23 | fibronectin 1b | fn1b | 2.9 | 4.0 | 5.7 | 4.9 | 4.0 | Yes |
| 24 | GINS complex subunit 2 | gins2 | 3.0 | 4.2 | 2.9 | 3.1 | 2.7 | Yes |
| 25 | gonadal somatic cell derived factor | gsdf | -2.3 | -2.4 | -2.8 | -2.7 | -2.4 | Yes |
| 26 | GTP binding protein 4 | gtpbp4 | -2.6 | -3.4 | -2.5 | -2.6 | -2.5 | Yes |
| 27 | hyaluronan synthase 1 | has1 | 3.7 | 3.5 | 2.6 | 3.1 | 4.1 | Yes |
| 28 | hydroxysteroid (17-beta) dehydrogenase 7 | hsd17b7 | -1.6 | -1.4 | -1.4 | -1.8 | -2.1 | Yes |
| 29 | inhibin, beta Aa | inhbaa | 3.8 | 2.6 | 2.3 | 3.4 | 3.7 | Yes |
| 30 | interleukin 1, beta | il1b | 3.4 | 2.6 | 3.3 | 2.6 | 2.9 | Yes |
| 31 | interleukin 2 receptor, beta | il2rb | -1.6 | -2.1 | -2.7 | -1.9 | -2.2 | Yes |
| 32 | interleukin 7 receptor | il7r | -1.6 | -1.7 | -2.4 | -1.8 | -1.4 | Yes |
| 33 | isocitrate dehydrogenase 2 (NADP+), mitochondrial | idh2 | 1.3 | 1.7 | 1.5 | 1.9 | 1.7 | Yes |
| 34 | keratin 18 | krt18 | 6.1 | 6.1 | 5.7 | 4.6 | 3.8 | Yes |
| 35 | leptin b | lepb | 5.8 | 6.3 | 9.4 | 9.6 | 9.3 | Yes |
| 36 | matrix metalloproteinase 13a | mmp13a | 4.3 | 2.9 | 2.4 | 4.9 | 5.0 | Yes |
| 37 | matrix metalloproteinase 9 | mmp9 | 3.8 | 2.5 | 3.7 | 4.4 | 4.0 | Yes |
| 38 | midkine-related growth factor | mdka | 2.4 | 2.7 | 4.2 | 4.9 | 4.2 | Yes |
| 39 | serine protease 3 | PRSS3 | -2.2 | -2.8 | -4.3 | -3.4 | -3.3 | Yes |
| 40 | synemin | synm | -1.8 | -2.6 | -2.6 | -1.6 | -2.1 | Yes |
| 41 | NA | traf7l | -1.8 | -1.9 | -2.0 | -1.8 | -2.1 | Yes |
| 42 | nitric oxide synthase 1 (neuronal) | nos1 | -2.0 | -2.3 | -3.4 | -3.1 | -2.9 | Yes |
| 43 | origin recognition complex, subunit 3 | orc3 | 2.1 | 2.8 | 1.8 | 2.2 | 1.7 | Yes |
| 44 | origin recognition complex, subunit 6 | orc6 | 2.1 | 3.8 | 3.5 | 3.2 | 2.0 | Yes |
| 45 | paired box gene 7b | pax7b | -1.7 | -2.6 | -2.8 | -3.7 | -2.7 | Yes |
| 46 | Pim-1 proto-oncogene, serine/threonine kinase | pim1 | -2.1 | -1.5 | -2.3 | -2.6 | -2.2 | Yes |
| 47 | poly(A) binding protein, cytoplasmic 4 (inducible form) | pabpc4 | 3.0 | 3.2 | 3.2 | 3.2 | 3.0 | Yes |
| 48 | prostaglandin D2 synthase b | ptgdsb | -2.3 | -2.6 | -2.8 | -3.6 | -1.9 | Yes |
| 49 | prostate transmembrane protein, androgen induced 1 | pmepa1 | 2.7 | 1.6 | 1.7 | 1.7 | 1.9 | Yes |
| 50 | proteasome (prosome, macropain) subunit, beta type, 7 | psmb7 | 3.1 | 2.5 | 2.8 | 2.9 | 2.5 | Yes |
| 51 | protein phosphatase 1, regulatory (inhibitor) subunit 3Cb | ppp1r3cb | -1.9 | -1.6 | -2.3 | -2.2 | -2.2 | Yes |
| 52 | RAS-like, family 11, member B | rasl11b | 3.0 | 2.0 | 2.5 | 2.8 | 2.4 | Yes |
| 53 | regulator of G-protein signaling 5a | rgs5a | -1.4 | -2.1 | -2.0 | -2.1 | -2.1 | Yes |
| 54 | regulator of G-protein signalling 4 | rgs4 | -2.2 | -2.6 | -1.9 | -2.6 | -2.0 | Yes |
| 55 | relaxin/insulin-like family peptide receptor 2b | rxfp2b | -3.0 | -2.2 | -2.4 | -3.5 | -2.4 | Yes |
| 56 | retinol dehydrogenase 12-like | LOC100150965 | -1.9 | -2.6 | -3.0 | -4.4 | -3.2 | Yes |
| 57 | Rho-related BTB domain containing 2a | rhobtb2a | -2.6 | -2.2 | -2.2 | -2.0 | -2.4 | Yes |
| 58 | secretory calcium-binding phosphoprotein 8 | scpp8 | 6.4 | 5.1 | 2.6 | 3.7 | 3.7 | Yes |
| 59 | si:ch211-122l24.4 | si:ch211-122l24 | 3.1 | 2.5 | 2.7 | 3.2 | 2.0 | Yes |
| 60 | si:dkey-33b17.3 ; MHC class II | si:busm1-160c18.3 | -4.1 | -3.8 | -4.1 | -2.9 | -4.1 | Yes |
| 61 | si:dkey-51e6.1 | si:dkey-51e6 | -1.8 | -1.9 | -2.2 | -2.0 | -1.9 | Yes |
| 62 | si:dkeyp-113d7.4 | si:dkeyp-113d7 | -3.9 | -4.3 | -2.6 | -3.3 | -1.8 | Yes |
| 63 | smoothelin, like | smtnl | 3.7 | 3.9 | 3.3 | 2.8 | 2.6 | Yes |
| 64 | solute carrier family 3, member 2b | slc3a2b | 2.3 | 1.4 | 1.8 | 1.7 | 1.8 | Yes |
| 65 | solute carrier family 6 (neurotransmitter transporter, taurine), member 6b | slc6a6b | -3.8 | -3.5 | -3.4 | -3.4 | -2.4 | Yes |
| 66 | spermidine/spermine N1-acetyltransferase 1a, duplicate 1 ; diamine N-acetyltransferase 1 | sat1a | -2.1 | -2.2 | -2.0 | -2.1 | -1.8 | Yes |
| 67 | SRY-box containing gene 11a | sox11a | 2.4 | 2.2 | 2.8 | 2.9 | 1.9 | Yes |
| 68 | stearoyl-CoA desaturase b | scdb | -1.8 | -2.6 | -2.4 | -2.3 | -2.8 | Yes |
| 69 | sulfotransferase family 1, cytosolic sulfotransferase 6 | sult1st6 | -3.9 | -2.5 | -3.2 | -2.9 | -2.2 | Yes |
| 70 | suppressor of cytokine signaling 3b | socs3b | 2.4 | 2.3 | 2.6 | 2.3 | 1.8 | Yes |
| 71 | synaptotagmin IV | syt4 | 5.3 | 3.5 | 3.3 | 3.2 | 5.4 | Yes |
| 72 | thioredoxin | txn | 3.3 | 2.7 | 3.1 | 2.6 | 2.4 | Yes |
| 73 | thioredoxin domain containing 12 (endoplasmic reticulum) | txndc12 | 2.2 | 1.9 | 2.4 | 3.1 | 2.1 | Yes |
| 74 | tissue inhibitor of metalloproteinase 2b | timp2b | 4.1 | 4.1 | 2.8 | 4.3 | 3.4 | Yes |
| 75 | transcription elongation factor A (SII), 3 | tcea3 | -1.9 | -1.9 | -1.6 | -1.8 | -2.0 | Yes |
| 76 | tropomyosin 4a | tpm4a | 5.0 | 4.0 | 2.1 | 2.3 | 2.7 | Yes |
| 77 | tubulin, alpha 1b | tuba1b | 3.3 | 3.1 | 4.0 | 4.6 | 3.9 | Yes |
| 78 | zgc:103599 | zgc:103599 | -2.4 | -2.2 | -1.6 | -1.4 | -1.9 | Yes |
| 79 | zgc:123339 | zgc:123339 | -2.1 | -2.0 | -1.8 | -1.7 | -1.6 | Yes |
| 80 | zgc:136474 | zgc:136474 | -2.2 | -2.4 | -2.1 | -1.8 | -1.8 | Yes |
| 81 | zgc:136902 | zgc:136902 | 5.0 | 5.5 | 5.5 | 4.6 | 4.5 | Yes |
| 82 | zgc:136930 | zgc:136930 | -4.1 | -4.6 | -3.5 | -3.7 | -2.5 | Yes |
| 83 | zgc:153031 | zgc:153031 | -2.2 | -3.7 | -2.5 | -3.3 | -2.9 | Yes |
| 84 | zgc:153154 | zgc:153154 | -2.2 | -2.7 | -2.9 | -3.3 | -2.3 | Yes |
| 85 | zgc:153629 | zgc:153629 | 2.2 | 3.9 | 3.4 | 3.0 | 2.3 | Yes |
| 86 | zgc:153920 | zgc:153920 | -3.0 | -2.6 | -1.8 | -1.6 | -2.9 | Yes |
| 87 | zgc:154093 | zgc:154093 | 2.3 | 2.1 | 2.2 | 1.5 | 2.0 | Yes |
| 88 | zgc:162356 | zgc:162356 | -1.9 | -1.3 | -1.8 | -1.7 | -1.7 | Yes |
| 89 | zgc:163079 | zgc:163079 | -2.4 | -2.7 | -2.4 | -2.8 | -3.1 | Yes |
| 90 | zgc:174917 | zgc:174917 | 4.8 | 3.1 | 6.4 | 3.8 | 4.1 | Yes |
| 91 | zgc:194409 | zgc:194409 | -2.3 | -2.4 | -1.5 | -1.4 | -2.4 | Yes |
| 92 | zgc:194993 | zgc:194993 | -4.6 | -3.4 | -4.3 | -3.8 | -2.1 | Yes |
| 93 | zgc:77358 | zgc:77358 | 1.7 | 2.6 | 4.6 | 4.1 | 3.2 | Yes |
| 94 | zinc finger and BTB domain containing 16a | zbtb16a | -4.0 | -3.4 | -3.9 | -3.3 | -3.8 | Yes |
| 95 | adhesion G protein-coupled receptor F3 | ADGRF3 | -1.7 | -1.7 | -1.7 | -2.0 | -1.6 | Yes |
| 96 | B and T lymphocyte associated | Btla | -4.3 | -2.9 | -4.4 | -5.8 | -3.5 | Yes |
| 97 | calpain 2 | capn2 | -3.8 | -3.6 | -2.2 | -2.7 | -2.6 | Yes |
| 98 | None | CH211-76L23 | -2.5 | -3.1 | -2.2 | -3.8 | -3.0 | Yes |
| 99 | C-type lectin domain containing 19A | [clec19a](https://www.ncbi.nlm.nih.gov/gene/100491258) | -2.9 | -2.5 | -2.7 | -2.7 | -1.7 | Yes |
| 100 | collagen type IX alpha 3 chain | COL9A3 | -2.3 | -2.8 | -2.2 | -2.3 | -2.2 | Yes |
| 101 | Cold shock protein 2 | csp2 | 2.7 | 3.9 | 4.5 | 6.4 | 4.7 | Yes |
| 102 | chondroitin sulfate proteoglycan 4 | CSPG4 | -3.8 | -4.4 | -2.7 | -3.4 | -3.0 | Yes |
| 103 | cytoglobin 1 | CYGB1 | 2.5 | 4.9 | 6.3 | 7.5 | 5.8 | Yes |
| 104 | Cytochrome c oxidase subunit 1 | cypCar_00031856 | -3.9 | -3.3 | -2.7 | -2.8 | -3.9 | Yes |
| 105 | cell wall protein DAN4-like | DAN4 | -2.5 | -3.4 | -3.9 | -3.3 | -3.3 | Yes |
| 106 | None | DDB_G0282133 | -2.3 | -3.4 | -2.5 | -3.3 | -3.3 | Yes |
| 107 | None | DKEY-246P9 | -3.3 | -3.4 | -3.0 | -2.2 | -2.4 | Yes |
| 108 | None | DKEY-246P9 | -2.6 | -2.5 | -1.8 | -1.8 | -1.8 | Yes |
| 109 | None | DKEY-288C23 | -3.1 | -2.6 | -3.7 | -4.2 | -3.3 | Yes |
| 110 | EGF containing fibulin extracellular matrix protein 1 | [efemp1](https://www.google.co.in/url?sa=t&rct=j&q=&esrc=s&source=web&cd=1&cad=rja&uact=8&ved=0ahUKEwjm1KTL5t7YAhUMT7wKHQGyDq0QFggmMAA&url=http%3A%2F%2Fwww.uniprot.org%2Funiprot%2FQ5PR37&usg=AOvVaw14PUO3ZkXJgVNbWO0GNf2v) | -3.6 | -2.8 | -2.5 | -3.5 | -2.8 | Yes |
| 111 | alpha(1,3)fucosyltransferase gene 2 | [ft2](https://www.ncbi.nlm.nih.gov/gene/111265018) | -3.0 | -3.1 | -4.8 | -2.8 | -3.8 | Yes |
| 112 | XPG-like endonuclease | GEN | 2.6 | 3.5 | 2.1 | 2.5 | 2.1 | Yes |
| 113 | hydroxysteroid (17-beta) dehydrogenase 12a | hsd17b12a | -2.5 | -2.9 | -2.4 | -2.2 | -4.0 | Yes |
| 114 | potassium inwardly-rectifying channel, subfamily J, member 1a, tandem duplicate 1 | kcnj1a | -2.7 | -3.7 | -2.8 | -4.6 | -2.7 | Yes |
| 115 | None | LOC100330916 | -1.4 | -2.0 | -2.2 | -2.4 | -1.5 | Yes |
| 116 | None | LOC107729934 | -2.4 | -2.9 | -3.0 | -2.6 | -4.1 | Yes |
| 117 | None | LOC110438252 | -2.0 | -2.6 | -3.8 | -2.7 | -2.7 | Yes |
| 118 | None | N/A | -2.0 | -2.0 | -2.7 | -2.8 | -1.6 | Yes |
| 119 | protocadherin 1 | [PCDH1](https://www.ncbi.nlm.nih.gov/gene/416194) | -2.6 | -2.4 | -3.3 | -2.9 | -2.1 | Yes |
| 120 | plexin A2 | plxn a2 | 2.1 | 2.7 | 2.6 | 3.4 | 2.8 | Yes |
| 121 | chromosome 1 open reading frame 116 | sarg | -1.5 | -1.7 | -1.7 | -1.9 | -1.5 | Yes |
| 122 | chromosome 1 open reading frame 116 | si:ch211-126j24 | 5.4 | 3.6 | 4.9 | 4.2 | 2.7 | Yes |
| 123 | None | si:ch211-153b23.3 | 3.2 | 3.3 | 6.7 | 3.0 | 2.5 | Yes |
| 124 | None | si:ch211-156b7.4 | 1.8 | 3.2 | 2.0 | 2.9 | 1.9 | Yes |
| 125 | None | si:ch211-252f13.5 | -2.7 | -2.8 | -1.8 | 1.7 | 2.3 | Yes |
| 126 | None | si:dkey-151j17.4 | -2.0 | -2.0 | -2.4 | -1.7 | -1.8 | Yes |
| 127 | solute carrier family 7 member 1 | SLC7A1 | -3.2 | -2.5 | -3.0 | -2.5 | -4.0 | Yes |
| 128 | solute carrier family 7 member 1 | SLK | -2.1 | -2.1 | -2.6 | -1.9 | -2.5 | Yes |
| 129 | STE20 like kinase | USP2 | -2.7 | -2.6 | -2.8 | -2.0 | -1.7 | Yes |
| 130 | ubiquitin specific peptidase 2 | wu:fc46h12 | -2.0 | -2.9 | -3.2 | -3.8 | -1.7 | Yes |
| 131 | Septin 5 | Sept5 | 0.5 | 0.8 | 1.4 | 1.8 | 1.5 | No |
| 132 | septin 5 | 0610037L13RIK | 0.8 | 1.2 | 0.8 | 1.0 | 0.4 | No |
| 133 | None | 1110004F10RIK | 1.1 | 1.2 | 0.6 | 0.8 | 0.4 | No |
| 134 | None | 1110059E24RIK | 0.5 | 0.3 | 0.4 | 0.6 | 0.5 | No |
| 135 | None | 2210016L21RIK | 1.1 | 1.3 | 0.6 | 0.6 | 0.3 | No |
| 136 | None | 4921524J17RIK | 0.8 | 1.3 | 1.3 | 1.6 | 1.3 | No |
| 137 | None | 4932438A13RIK | -0.9 | -1.1 | -0.6 | -2.1 | -2.4 | No |
| 138 | None | 9330182L06RIK | -1.2 | -2.0 | -7.3 | -0.3 | -15.1 | No |
| 139 | None | A630033H20RIK | -0.7 | -1.8 | -0.8 | -1.1 | -2.1 | No |
| 140 | acetoacetyl-CoA synthetase | AACS | 1.7 | 1.3 | 1.0 | 0.1 | 1.1 | No |
| 141 | alpha and gamma adaptin binding protein | AAGAB | 0.9 | 1.1 | 0.7 | 0.6 | 0.5 | No |
| 142 | aralkylamine N-acetyltransferase | AANAT | 2.9 | 3.2 | 3.0 | 2.5 | 2.3 | No |
| 143 | apoptosis antagonizing transcription factor | AATF | 2.1 | 1.6 | 0.8 | 0.7 | -0.7 | No |
| 144 | ATP binding cassette subfamily B member 8 | ABCB8 | 0.7 | 1.0 | 0.6 | 0.9 | 0.6 | No |
| 145 | ATP binding cassette subfamily C member 4 | ABCC4 | 1.2 | 0.6 | 1.1 | 0.6 | 0.1 | No |
| 146 | ATP binding cassette subfamily D member 4 | ABCD4 | -0.8 | -0.7 | -1.2 | -1.4 | -0.9 | No |
| 147 | ATP binding cassette subfamily E member 1 | ABCE1 | 1.5 | 1.4 | 0.8 | 0.3 | 0.4 | No |
| 148 | ATP binding cassette subfamily F member 1 | ABCF1 | 1.7 | 1.6 | 1.1 | 0.9 | 0.5 | No |
| 149 | ATP binding cassette subfamily F member 2 | ABCF2 | 2.4 | 2.0 | 1.7 | 1.4 | 1.3 | No |
| 150 | abl interactor 1 | ABI1 | 1.3 | 2.0 | 1.9 | 2.7 | 1.8 | No |
| 151 | actin binding LIM protein 1 | ABLIM1 | -2.5 | -1.4 | -2.4 | -2.0 | -0.7 | No |
| 152 | None | AC124712.1 | -0.6 | 0.0 | -1.8 | -1.0 | -1.4 | No |
| 153 | None | AC163637.2 | 1.2 | 1.6 | 0.8 | 0.3 | 0.3 | No |
| 154 | acyl-CoA dehydrogenase family member 11 | ACAD11 | -0.3 | 0.2 | -0.2 | 0.0 | -1.3 | No |
| 155 | acyl-CoA dehydrogenase family member 9 | ACAD9 | 1.1 | 0.6 | 1.0 | 1.4 | 0.6 | No |
| 156 | ArfGAP with coiled-coil, ankyrin repeat and PH domains 1 | ACAP1 | -1.0 | -0.8 | -1.0 | -1.2 | -1.1 | No |
| 157 | Acat3 acetyl-Coenzyme A acetyltransferase 3 | ACAT3 | 1.6 | 2.7 | 1.3 | 1.8 | 1.4 | No |
| 158 | acetylcholinesterase (Cartwright blood group) | ACHE | -2.5 | -1.6 | -1.4 | -13.7 | -1.9 | No |
| 159 | acyl-CoA thioesterase 7 | ACOT7 | 1.7 | 0.8 | 1.6 | 1.2 | 0.1 | No |
| 160 | acyl-CoA thioesterase 8 | ACOT8 | 0.8 | 1.0 | 0.8 | 1.4 | 0.5 | No |
| 161 | acid phosphatase 1 | ACP1 | 1.3 | 1.8 | 0.8 | 1.1 | 0.9 | No |
| 162 | acid phosphatase 2, lysosomal | ACP2 | -1.3 | -3.9 | -0.9 | -0.8 | -1.6 | No |
| 163 | acyl-CoA synthetase long chain family member 3 | ACSL3 | 0.4 | 0.9 | 0.4 | 0.3 | 0.2 | No |
| 164 | acyl-CoA synthetase long chain family member 4 | ACSL4 | 1.0 | 0.1 | 0.4 | 0.4 | 0.4 | No |
| 165 | actin like 6A | ACTL6A | 1.8 | 1.9 | 1.6 | 1.6 | 1.1 | No |
| 166 | ARP1 actin related protein 1 homolog B | ACTR1B | 0.7 | 1.0 | 0.7 | 0.7 | 0.5 | No |
| 167 | ARP2 actin related protein 2 homolog | ACTR2 | -0.7 | -1.2 | -0.5 | -0.8 | -0.9 | No |
| 168 | ARP6 actin related protein 6 homolog | ACTR6 | 0.9 | 1.0 | 0.6 | 0.6 | -0.1 | No |
| 169 | adenosine deaminase | ADA | -0.6 | -0.7 | -0.8 | -0.8 | -1.3 | No |
| 170 | ADAM metallopeptidase domain 10 | ADAM10 | 0.6 | 0.0 | 1.4 | 1.4 | 1.2 | No |
| 171 | ADAM metallopeptidase domain 8 | ADAM8 | 6.2 | 6.5 | 6.7 | 6.5 | 4.8 | No |
| 172 | ADAM metallopeptidase domain 9 | ADAM9 | 0.0 | 0.2 | 0.6 | 1.1 | 0.7 | No |
| 173 | adenosine deaminase, tRNA specific 2 | ADAT2 | 2.3 | 1.9 | 1.3 | 2.1 | 1.7 | No |
| 174 | adenylate cyclase 8 | ADCY8 | 3.3 | 4.1 | 1.7 | 1.4 | 1.4 | No |
| 175 | adenylate cyclase activating polypeptide 1 | ADCYAP1 | -0.9 | -0.4 | -2.0 | -1.4 | -1.2 | No |
| 176 | adenosine kinase | ADK | 0.5 | 1.1 | 0.8 | 0.8 | 0.6 | No |
| 177 | adrenomedullin 2 | ADM2 | 4.2 | 2.5 | 3.3 | 3.8 | 3.3 | No |
| 178 | ADP-ribosylhydrolase like 2 | ADPRHL2 | 0.5 | 1.1 | 0.4 | 0.5 | 0.4 | No |
| 179 | adrenoceptor alpha 2A | ADRA2A | -0.4 | -1.2 | -1.5 | -3.2 | -1.7 | No |
| 180 | acylglycerol kinase | AGK | 1.1 | 1.1 | 0.5 | 0.8 | 0.4 | No |
| 181 | None | AI837181 | -1.5 | -0.9 | -0.8 | -0.5 | -0.7 | No |
| 182 | aminoacyl tRNA synthetase complex interacting multifunctional protein 1 | AIMP1 | 1.8 | 1.6 | 0.9 | 1.0 | 0.7 | No |
| 183 | autoimmune regulator | AIRE | -0.8 | -0.5 | -0.9 | -1.1 | 0.4 | No |
| 184 | adenylate kinase 1 | AK1 | -1.3 | -3.2 | -0.9 | -3.3 | -1.9 | No |
| 185 | activated leukocyte cell adhesion molecule | ALCAM | 0.8 | 0.6 | 1.3 | 1.4 | 0.1 | No |
| 186 | aldehyde dehydrogenase 1 family member A2 | ALDH1A2 | 0.4 | -0.3 | 2.2 | 3.1 | 1.3 | No |
| 187 | aldehyde dehydrogenase 3 family member B1 | ALDH3B1 | 0.9 | 1.0 | 0.5 | 0.4 | 0.3 | No |
| 188 | aldehyde dehydrogenase 6 family member A1 | ALDH6A1 | -0.8 | -0.7 | -0.5 | -0.5 | -0.4 | No |
| 189 | aldolase, fructose-bisphosphate B | ALDOB | -2.1 | -2.1 | -1.3 | -1.6 | -1.6 | No |
| 190 | ALG2, alpha-1,3/1,6-mannosyltransferase | ALG2 | 1.6 | 1.5 | 0.8 | 0.9 | 0.8 | No |
| 191 | ALG3, alpha-1,3- mannosyltransferase | ALG3 | 0.8 | 0.7 | 0.8 | 1.2 | 0.7 | No |
| 192 | ALG9, alpha-1,2-mannosyltransferase | ALG9 | 1.5 | 1.9 | 1.2 | 1.8 | 1.4 | No |
| 193 | alkB homolog 7 | ALKBH7 | 0.6 | 0.7 | 0.8 | 1.1 | 0.8 | No |
| 194 | alkaline phosphatase, germ cell | ALPPL2 | 0.3 | 1.4 | 2.9 | 3.8 | 3.0 | No |
| 195 | anti-Mullerian hormone | AMH | -0.8 | -1.4 | -1.5 | -1.2 | -2.0 | No |
| 196 | Alport syndrome, mental retardation, midface hypoplasia and elliptocytosis chromosomal region gene 1 | AMMECR1 | 0.4 | 0.4 | 0.8 | 0.7 | 0.4 | No |
| 197 | amphiphysin | AMPH | -1.4 | -2.2 | -1.2 | -5.7 | -2.8 | No |
| 198 | anaphase promoting complex subunit 4 | ANAPC4 | 8.4 | 8.4 | 8.2 | 16.1 | 15.8 | No |
| 199 | anaphase promoting complex subunit 5 | ANAPC5 | 0.7 | 0.9 | 0.8 | 1.7 | 1.3 | No |
| 200 | angiopoietin like 2 | ANGPTL2 | 3.0 | 3.8 | 3.8 | 4.4 | 4.4 | No |
| 201 | Ankyrin | ANK | 1.2 | 0.3 | 0.3 | 0.7 | 0.5 | No |
| 202 | ankyrin repeat domain 10 | ANKRD10 | 0.5 | 1.3 | 0.7 | 1.4 | 1.0 | No |
| 203 | ankyrin repeat domain 6 | ANKRD6 | -0.6 | -1.8 | -1.5 | -1.0 | -0.9 | No |
| 204 | ankyrin repeat domain 65 | ANKRD65 | -2.7 | -2.3 | -1.5 | -1.4 | -1.4 | No |
| 205 | ankyrin repeat and sterile alpha motif domain containing 1B | ANKS1B | -0.2 | -0.9 | 0.5 | 0.4 | 0.1 | No |
| 206 | anoctamin 5 | ANO5 | 2.1 | 3.9 | 5.8 | 13.5 | 0.9 | No |
| 207 | acidic nuclear phosphoprotein 32 family member B | ANP32B | 1.2 | 1.6 | 1.0 | 1.1 | 0.8 | No |
| 208 | annexin A2 | ANXA2 | 2.2 | 2.3 | 1.3 | 0.8 | 1.0 | No |
| 209 | annexin A3 | ANXA3 | -0.4 | -0.2 | -0.6 | -0.3 | -2.1 | No |
| 210 | adaptor related protein complex 2 subunit mu 1 | AP2M1 | 1.0 | 0.4 | 0.5 | 1.2 | 0.1 | No |
| 211 | adaptor related protein complex 3 subunit mu 1 | AP3M1 | 0.1 | 0.5 | 0.6 | 1.0 | 0.5 | No |
| 212 | adaptor related protein complex 3 subunit mu 2 | AP3M2 | 0.7 | 0.6 | 0.7 | 0.7 | 0.5 | No |
| 213 | adaptor related protein complex 3 subunit sigma 2 | AP3S2 | 0.4 | 0.2 | 0.4 | 1.0 | 0.7 | No |
| 214 | aph-1 homolog B, gamma-secretase subunit | APH1B | 0.7 | 0.8 | 0.3 | 0.2 | 0.1 | No |
| 215 | apoptosis inhibitor 5 | API5 | 1.3 | 0.3 | 1.4 | 0.4 | -0.8 | No |
| 216 | apelin receptor | APLNR | 1.8 | 2.2 | 1.1 | 1.3 | 1.5 | No |
| 217 | adipocyte plasma membrane associated protein | APMAP | 1.9 | 1.5 | 0.6 | 0.9 | 0.5 | No |
| 218 | apolipoprotein L domain containing 1 | APOLD1 | -0.8 | -0.5 | -1.1 | -1.5 | -1.3 | No |
| 219 | adaptor protein, phosphotyrosine interacting with PH domain and leucine zipper 1 | APPL1 | -0.7 | -1.3 | -0.5 | 0.0 | -0.5 | No |
| 220 | adenine phosphoribosyltransferase | APRT | 1.6 | 1.2 | 0.9 | 0.5 | 0.5 | No |
| 221 | aquarius intron-binding spliceosomal factor | AQR | 1.0 | 1.3 | 0.3 | 0.4 | 0.2 | No |
| 222 | androgen receptor | AR | -2.1 | -1.5 | -1.7 | -2.3 | -2.6 | No |
| 223 | archain 1 | ARCN1 | 0.5 | 0.4 | 0.5 | 1.0 | 0.7 | No |
| 224 | ADP ribosylation factor 4 | ARF4 | 1.0 | 1.1 | 0.6 | 0.3 | 0.0 | No |
| 225 | arginine and glutamate rich 1 | ARGLU1 | -1.6 | -0.9 | -1.5 | -1.1 | -1.1 | No |
| 226 | Rho/Rac guanine nucleotide exchange factor 18 | ARHGEF18 | -0.3 | -0.3 | -0.7 | -1.4 | -0.8 | No |
| 227 | Rho/Rac guanine nucleotide exchange factor 2 | ARHGEF2 | 0.4 | 0.4 | 1.0 | 1.0 | 0.5 | No |
| 228 | Rho guanine nucleotide exchange factor 39 | ARHGEF39 | -0.1 | 2.6 | 1.5 | 1.3 | 1.8 | No |
| 229 | ADP ribosylation factor like GTPase 11 | ARL11 | -0.5 | -0.2 | -1.2 | -0.7 | -0.1 | No |
| 230 | ADP ribosylation factor like GTPase 13B | ARL13B | 0.7 | 0.2 | 0.8 | 1.2 | 0.8 | No |
| 231 | ADP ribosylation factor like GTPase 14 effector protein | ARL14EP | 0.6 | 1.1 | 0.7 | 1.0 | 0.6 | No |
| 232 | ADP ribosylation factor like GTPase 4D | ARL4D | -0.8 | -1.7 | -2.1 | -1.1 | -0.9 | No |
| 233 | ADP ribosylation factor like GTPase 5C | ARL5C | -0.1 | -0.9 | -1.1 | -0.5 | -0.2 | No |
| 234 | ADP ribosylation factor like GTPase 6 interacting protein 5 | ARL6IP5 | -1.2 | -1.0 | -1.1 | -0.8 | -0.4 | No |
| 235 | aryl hydrocarbon receptor nuclear translocator like 2 | ARNTL2 | 1.6 | 0.9 | 0.8 | 0.5 | 0.4 | No |
| 236 | actin related protein 2/3 complex subunit 3 | ARPC3 | 0.6 | 1.0 | 0.9 | 0.6 | 0.3 | No |
| 237 | actin related protein 2/3 complex subunit 5 | ARPC5 | 0.8 | 1.1 | 0.6 | 0.5 | 0.4 | No |
| 238 | cAMP regulated phosphoprotein 19 | ARPP19 | 0.0 | 0.1 | 0.1 | 0.1 | 0.0 | No |
| 239 | arrestin beta 2 | ARRB2 | 0.5 | 1.0 | 0.7 | 0.7 | 0.5 | No |
| 240 | arrestin domain containing 2 | ARRDC2 | -1.2 | -1.1 | -0.8 | -0.3 | -0.8 | No |
| 241 | arrestin domain containing 3 | ARRDC3 | -0.5 | -0.1 | -0.4 | -0.2 | -0.1 | No |
| 242 | ARVCF, delta catenin family member | ARVCF | 0.6 | 1.1 | 1.0 | 1.7 | 2.5 | No |
| 243 | arsenite methyltransferase | AS3MT | -1.0 | -0.7 | -0.6 | -0.8 | -0.6 | No |
| 244 | ArfGAP with SH3 domain, ankyrin repeat and PH domain 2 | ASAP2 | 2.6 | 3.0 | 3.4 | 2.9 | 1.9 | No |
| 245 | anti-silencing function 1B histone chaperone | ASF1B | 0.6 | 1.3 | 0.5 | 0.5 | 0.4 | No |
| 246 | argininosuccinate lyase | ASL | -0.5 | -5.9 | 0.3 | -1.6 | -10.1 | No |
| 247 | arsA arsenite transporter, ATP-binding, homolog 1 (bacterial) | ASNA1 | 0.1 | 0.0 | 0.4 | 0.1 | 0.1 | No |
| 248 | activating transcription factor 3 | ATF3 | 2.0 | 2.0 | 1.3 | 0.7 | 1.0 | No |
| 249 | activating transcription factor 5 | ATF5 | 2.1 | 1.4 | 2.0 | 1.1 | 1.3 | No |
| 250 | activating transcription factor 7 | ATF7 | -1.7 | -1.0 | -1.2 | -1.0 | -3.1 | No |
| 251 | autophagy related 9A | ATG9A | -0.8 | -1.1 | -0.6 | -1.7 | -1.3 | No |
| 252 | ATPase Na+/K+ transporting subunit beta 3 | ATP1B3 | -1.7 | -1.0 | -0.8 | -0.2 | -0.6 | No |
| 253 | ATP23 metallopeptidase and ATP synthase assembly factor homolog | ATP23 | 0.8 | 1.0 | 0.5 | 0.7 | 0.0 | No |
| 254 | ATP synthase, H+ transporting, mitochondrial F1 complex, delta subunit | ATP5D | 0.7 | 1.2 | 0.3 | 0.4 | 0.3 | No |
| 255 | ATP synthase membrane subunit c locus 1 | ATP5G1 | 1.3 | 1.8 | 1.3 | 1.5 | 1.0 | No |
| 256 | ATP synthase mitochondrial F1 complex assembly factor 2 | ATPAF2 | 0.9 | 0.8 | 0.4 | 0.4 | 0.1 | No |
| 257 | ATPase inhibitor A, mitochondrial (Inhibitor of F(1)F(o)-ATPase A) (IF(1) A) (IF1 A) | ATPIF1 | 1.6 | 2.0 | 1.1 | 1.3 | 0.9 | No |
| 258 | ataxin 2 | ATXN2 | -0.9 | -0.2 | -1.2 | -4.7 | -1.5 | No |
| 259 | aurora kinase A | AURKA | 1.4 | 2.9 | 3.4 | 3.8 | 3.8 | No |
| 260 | aurora kinase C | AURKC | 0.8 | 2.6 | 2.2 | 2.8 | 2.7 | No |
| 261 | UDP-GlcNAc:betaGal beta-1,3-N-acetylglucosaminyltransferase 2 | B3GNT2 | -0.9 | -0.9 | -1.4 | -1.0 | -1.1 | No |
| 262 | UDP-GlcNAc:betaGal beta-1,3-N-acetylglucosaminyltransferase like 1 | B3GNTL1 | 0.9 | 0.8 | 0.6 | 0.8 | 0.1 | No |
| 263 | beta-1,4-galactosyltransferase 7 | B4GALT7 | 1.2 | 0.9 | 1.5 | 1.6 | 0.8 | No |
| 264 | B9 domain containing 2 | B9D2 | 1.0 | 1.3 | 1.1 | -10.6 | -0.4 | No |
| 265 | BRISC and BRCA1 A complex member 1 | BABAM1 | -1.1 | -0.3 | -0.7 | -0.4 | -0.8 | No |
| 266 | BCL2 associated agonist of cell death | BAD | -0.7 | -0.5 | -0.5 | -0.8 | -0.1 | No |
| 267 | BCL2 associated athanogene 4 | BAG4 | 0.4 | 0.6 | 0.5 | 0.5 | 0.3 | No |
| 268 | BAI1 associated protein 2 like 1 | BAIAP2L1 | -1.2 | -1.4 | -1.5 | -2.1 | -1.6 | No |
| 269 | BCL2 associated X, apoptosis regulator | BAX | 0.3 | 0.3 | 0.2 | 0.1 | -0.2 | No |
| 270 | gamma-butyrobetaine hydroxylase 1 | BBOX1 | -1.0 | -0.9 | -1.0 | 0.3 | 0.6 | No |
| 271 | Bardet-Biedl syndrome 4 | BBS4 | 1.1 | 0.8 | 0.9 | 0.8 | 0.9 | No |
| 272 | Bardet-Biedl syndrome 7 | BBS7 | 1.1 | 0.9 | 1.3 | 2.2 | 1.8 | No |
| 273 | basal cell adhesion molecule (Lutheran blood group) | BCAM | 2.1 | 0.6 | 5.2 | 4.3 | 4.0 | No |
| 274 | branched chain amino acid transaminase 1 | BCAT1 | 4.5 | 1.7 | 3.7 | 4.2 | 5.1 | No |
| 275 | branched chain keto acid dehydrogenase E1 subunit beta | BCKDHB | 1.1 | 0.8 | 1.3 | 0.8 | 0.7 | No |
| 276 | B cell CLL/lymphoma 11A | BCL11A | -1.1 | -1.6 | -1.3 | -2.3 | -1.8 | No |
| 277 | biglycan | BGN | -0.9 | -1.5 | -0.9 | -1.0 | -1.3 | No |
| 278 | basic helix-loop-helix family member e40 | BHLHE40 | -2.6 | -1.2 | -1.0 | -1.7 | -1.3 | No |
| 279 | betaine--homocysteine S-methyltransferase 2 | BHMT2 | -1.3 | -1.5 | -1.8 | -1.8 | -1.3 | No |
| 280 | bridging integrator 2 | BIN2 | 0.4 | 1.4 | 2.2 | 0.5 | 2.2 | No |
| 281 | bone morphogenetic protein 1 | BMP1 | 0.3 | 0.7 | 0.3 | 0.2 | 0.1 | No |
| 282 | bone morphogenetic protein 2 | BMP2 | -0.2 | 0.4 | 0.6 | 0.6 | 1.2 | No |
| 283 | bone morphogenetic protein 6 | BMP6 | 1.1 | 0.3 | 1.4 | 3.0 | 2.0 | No |
| 284 | BCL2 interacting protein 3 | BNIP3 | -1.4 | -1.5 | -0.5 | 0.1 | -0.1 | No |
| 285 | BCL2 interacting protein 3 like | BNIP3L | -0.9 | -1.5 | -0.3 | -0.7 | -0.7 | No |
| 286 | block of proliferation 1 | BOP1 | 1.8 | 1.4 | 0.8 | 0.9 | 0.2 | No |
| 287 | B-Raf proto-oncogene, serine/threonine kinase | BRAF | -0.7 | -0.7 | -0.2 | -0.6 | -0.7 | No |
| 288 | BTB domain containing 6 | BTBD6 | 3.8 | 3.3 | 1.0 | 1.7 | 0.5 | No |
| 289 | B-cell translocation gene 1 (B-cell translocation gene 1) | BTG1-PS1 | -0.9 | -0.5 | -0.7 | -1.0 | -0.9 | No |
| 290 | BTG anti-proliferation factor 2 | BTG2 | -1.0 | -0.8 | -2.4 | -0.7 | -1.7 | No |
| 291 | BUD23, rRNA methyltransferase and ribosome maturation factor | BUD23 | 1.5 | 1.1 | 0.5 | 0.7 | 0.6 | No |
| 292 | BUD31 homolog | BUD31 | 1.1 | 1.0 | 0.9 | 0.6 | 0.3 | No |
| 293 | bystin like | BYSL | 1.4 | 0.9 | 0.4 | -0.2 | -0.1 | No |
| 294 | basic leucine zipper and W2 domains 1 | BZW1 | 1.2 | 1.2 | 0.8 | 1.2 | 0.7 | No |
| 295 | C2CD2 like | C2CD2L | -0.9 | -1.3 | -1.3 | -1.0 | -0.2 | No |
| 296 | complement C6 | C6 | 0.8 | 1.8 | 1.3 | 1.9 | 1.5 | No |
| 297 | calcium/calmodulin dependent protein kinase II beta | CAMK2B | -0.8 | 0.2 | -0.6 | -0.1 | -0.9 | No |
| 298 | calcium/calmodulin dependent protein kinase II inhibitor 2 | CAMK2N2 | 1.0 | 0.7 | 1.7 | 1.0 | 1.3 | No |
| 299 | calmodulin regulated spectrin associated protein 1 | CAMSAP1 | 0.5 | 0.6 | 1.4 | 1.5 | 1.7 | No |
| 300 | calcium activated nucleotidase 1 | CANT1 | -0.7 | -0.3 | -0.1 | 0.3 | -2.3 | No |
| 301 | calnexin | CANX | 0.9 | 1.1 | 0.8 | 0.6 | 0.6 | No |
| 302 | calpain 12 | CAPN12 | -1.4 | -1.6 | 0.3 | 0.9 | 0.6 | No |
| 303 | cell cycle associated protein 1 | CAPRIN1 | -1.3 | -1.8 | -1.4 | -1.8 | -1.6 | No |
| 304 | Carbonic Anhydrase 10 | CAR10 | 2.8 | 1.4 | 2.9 | 3.5 | 2.5 | No |
| 305 | Carbonic Anhydrase 13 | CAR13 | 0.8 | 1.1 | 1.0 | 0.9 | 1.0 | No |
| 306 | coactivator associated arginine methyltransferase 1 | CARM1 | 0.9 | 1.3 | 0.6 | 0.7 | 0.3 | No |
| 307 | cysteinyl-tRNA synthetase | CARS | 1.9 | 1.6 | 1.5 | 1.3 | 1.0 | No |
| 308 | CASC3, exon junction complex subunit | CASC3 | 0.3 | 0.2 | 0.2 | 0.0 | 0.0 | No |
| 309 | caspase 8 | CASP8 | 1.3 | 1.5 | 0.3 | 1.7 | 0.7 | No |
| 310 | caspase 9 | CASP9 | 0.7 | 0.4 | 0.6 | 0.6 | 0.4 | No |
| 311 | cystathionine-beta-synthase | CBS | 2.1 | 1.3 | 0.5 | -0.6 | -0.9 | No |
| 312 | coiled-coil domain containing 12 | CCDC12 | 1.1 | 1.3 | 0.5 | 0.4 | 0.1 | No |
| 313 | coiled-coil domain containing 125 | CCDC125 | -1.5 | -2.0 | -1.4 | -1.6 | -2.3 | No |
| 314 | coiled-coil domain containing 151 | CCDC151 | 0.6 | -1.8 | -0.7 | -0.5 | 0.0 | No |
| 315 | coiled-coil domain containing 24 | CCDC24 | -1.4 | -0.7 | -1.0 | -2.6 | -0.8 | No |
| 316 | coiled-coil domain containing 43 | CCDC43 | 1.4 | 1.4 | 0.6 | 0.4 | 0.0 | No |
| 317 | coiled-coil domain containing 51 | CCDC51 | 1.3 | 1.4 | 0.8 | 0.8 | 0.7 | No |
| 318 | coiled-coil domain containing 85B | CCDC85B | -0.1 | -0.5 | 1.5 | 2.2 | 2.1 | No |
| 319 | cyclin A2 | CCNA2 | 0.8 | 2.1 | 2.0 | 2.8 | 2.2 | No |
| 320 | cyclin B1 | CCNB1 | 0.6 | 2.8 | 2.8 | 4.0 | 3.3 | No |
| 321 | cyclin B2 | CCNB2 | 1.2 | 3.2 | 3.0 | 4.2 | 4.2 | No |
| 322 | chaperonin containing TCP1 subunit 3 | CCT3 | 1.0 | 0.9 | 0.5 | 0.6 | 0.3 | No |
| 323 | CD22 molecule | CD22 | 1.6 | 0.7 | 0.9 | 1.3 | 0.6 | No |
| 324 | CD2 associated protein | CD2AP | -6.1 | -0.1 | -5.3 | 0.0 | -9.4 | No |
| 325 | CD2 cytoplasmic tail binding protein 2 | CD2BP2 | 1.0 | 1.1 | 0.7 | 0.6 | 0.3 | No |
| 326 | CD63 molecule | CD63 | 1.5 | 0.9 | 0.7 | 0.5 | 0.3 | No |
| 327 | CD74 molecule | CD74 | -1.5 | -1.9 | -1.1 | -1.5 | -1.3 | No |
| 328 | CD99 molecule like 2 | CD99L2 | 0.8 | 1.5 | 2.2 | 1.9 | 2.3 | No |
| 329 | cytidine and dCMP deaminase domain containing 1 | CDADC1 | -0.5 | -0.3 | -0.5 | -0.3 | -0.2 | No |
| 330 | cell division cycle 23 | CDC23 | 0.8 | 1.6 | 1.2 | 1.4 | 0.9 | No |
| 331 | CDC42 effector protein 2 | CDC42EP2 | -1.4 | -0.9 | -2.1 | -1.0 | -1.8 | No |
| 332 | cell division cycle 45 | CDC45 | 0.8 | 2.4 | 1.4 | 2.0 | 1.7 | No |
| 333 | cell division cycle 6 | CDC6 | 1.4 | 1.7 | 1.7 | 1.0 | 0.8 | No |
| 334 | cell division cycle 7 | CDC7 | 2.6 | 4.1 | 2.8 | 5.0 | 4.8 | No |
| 335 | cell division cycle associated 2 | CDCA2 | 0.7 | 1.7 | 1.4 | 2.2 | 2.0 | No |
| 336 | cadherin 4 | CDH4 | -0.9 | -1.3 | -2.3 | -0.7 | -0.5 | No |
| 337 | CDP-diacylglycerol--inositol 3-phosphatidyltransferase | CDIPT | 0.6 | 0.9 | 0.4 | 0.6 | 0.5 | No |
| 338 | cyclin dependent kinase 10 | CDK10 | 0.8 | 0.9 | 0.9 | 1.0 | 0.7 | No |
| 339 | cyclin dependent kinase 2 | CDK2 | 1.3 | 2.1 | 1.1 | 1.6 | 1.3 | No |
| 340 | cyclin dependent kinase 8 | CDK8 | 0.8 | 1.2 | 0.8 | 1.2 | 0.5 | No |
| 341 | cyclin dependent kinase inhibitor 1B | CDKN1B | -1.4 | -0.8 | -0.4 | -0.1 | -0.3 | No |
| 342 | CDKN2A interacting protein N-terminal like | CDKN2AIPNL | 0.8 | 0.9 | 0.4 | 0.2 | -0.1 | No |
| 343 | CDV3 homolog | CDV3 | 1.2 | 1.3 | 0.8 | 0.5 | 0.5 | No |
| 344 | CCAAT enhancer binding protein gamma | CEBPG | -0.1 | -0.7 | -0.4 | -1.3 | -0.6 | No |
| 345 | CUGBP Elav-like family member 3 | CELF3 | -1.4 | -1.4 | -1.1 | -0.6 | -0.5 | No |
| 346 | centromere protein M | CENPM | 0.7 | 2.3 | 1.6 | 2.5 | 2.3 | No |
| 347 | centromere protein N | CENPN | 1.0 | 2.8 | 2.0 | 2.1 | 1.4 | No |
| 348 | centrosomal protein 135 | CEP135 | 1.2 | 2.5 | 1.4 | 2.0 | 2.0 | No |
| 349 | centrosomal protein 57 like 1 | CEP57L1 | 0.7 | 1.6 | 0.9 | 0.9 | 0.0 | No |
| 350 | ceramide synthase 4 | CERS4 | -1.9 | -1.4 | -1.6 | -1.9 | -2.9 | No |
| 351 | centrin 3 | CETN3 | 1.1 | 1.7 | 1.5 | 1.8 | 1.4 | No |
| 352 | cofilin 2 | CFL2 | 1.5 | 2.1 | 1.9 | 2.8 | 2.6 | No |
| 353 | cingulin | CGN | -0.5 | -0.9 | -0.7 | -1.0 | -0.5 | No |
| 354 | chromatin assembly factor 1 subunit A | CHAF1A | 0.3 | 1.2 | 0.7 | 1.1 | 0.8 | No |
| 355 | coiled-coil-helix-coiled-coil-helix domain containing 1 | CHCHD1 | 1.5 | 1.6 | 0.6 | 1.1 | 0.4 | No |
| 356 | coiled-coil-helix-coiled-coil-helix domain containing 10 | CHCHD10 | 2.7 | 2.8 | 1.6 | 1.4 | 1.4 | No |
| 357 | coiled-coil-helix-coiled-coil-helix domain containing 2 | CHCHD2 | 1.0 | 1.1 | 0.4 | 0.3 | 0.6 | No |
| 358 | checkpoint kinase 2 | CHEK2 | 1.0 | 2.1 | 0.8 | 1.2 | 0.6 | No |
| 359 | chromogranin A | CHGA | -1.5 | -1.6 | -1.5 | -2.0 | -3.3 | No |
| 360 | charged multivesicular body protein 4C | CHMP4C | -1.3 | -0.9 | -1.4 | -1.4 | -0.9 | No |
| 361 | chondrolectin | CHODL | -0.6 | 0.1 | -1.9 | -0.3 | -0.7 | No |
| 362 | calcineurin like EF-hand protein 2 | CHP2 | -1.3 | -1.4 | -1.1 | -1.7 | -1.3 | No |
| 363 | carbohydrate sulfotransferase 11 | CHST11 | 1.6 | 1.3 | 1.1 | 1.1 | 1.3 | No |
| 364 | carbohydrate sulfotransferase 14 | CHST14 | 0.3 | 0.1 | 1.4 | 1.7 | 1.4 | No |
| 365 | chromosome transmission fidelity factor 18 | CHTF18 | 2.0 | 3.1 | 2.2 | 3.0 | 2.3 | No |
| 366 | cytosolic iron-sulfur assembly component 1 | CIAO1 | 0.7 | 1.0 | 0.5 | 0.7 | 0.5 | No |
| 367 | cytokine induced apoptosis inhibitor 1 | CIAPIN1 | 1.2 | 1.0 | 0.8 | 0.2 | 0.6 | No |
| 368 | CDGSH iron sulfur domain 2 | CISD2 | 0.6 | 0.9 | 0.6 | 0.5 | 0.5 | No |
| 369 | cytokine inducible SH2 containing protein | CISH | -1.0 | -1.0 | -2.1 | -1.7 | -1.5 | No |
| 370 | CDC28 protein kinase regulatory subunit 1B | CKS1B | 1.0 | 2.4 | 2.1 | 2.8 | 2.6 | No |
| 371 | claudin 1 | CLDN1 | -0.4 | -0.9 | -0.5 | -1.2 | -0.4 | No |
| 372 | claudin 3 | CLDN3 | -0.7 | -0.7 | -1.3 | -1.1 | -0.7 | No |
| 373 | claudin 4 | CLDN4 | -1.5 | -1.8 | -0.7 | -1.2 | -1.8 | No |
| 374 | C-type lectin domain family 4 | CLEC4A4 | 5.2 | 3.8 | 1.8 | 2.1 | 0.5 | No |
| 375 | CDC like kinase 2 | CLK2 | -1.2 | -0.5 | -1.1 | -1.0 | -0.5 | No |
| 376 | CLPTM1, transmembrane protein | CLPTM1 | 1.0 | 0.4 | 0.7 | 0.3 | 0.3 | No |
| 377 | calsyntenin 1 | CLSTN1 | -1.6 | -2.1 | -0.5 | -0.1 | -0.7 | No |
| 378 | clathrin light chain A | CLTA | 1.3 | 1.1 | 1.1 | 1.4 | 0.7 | No |
| 379 | calponin 2 | CNN2 | 0.6 | 0.5 | 0.4 | 0.1 | 0.0 | No |
| 380 | CCR4-NOT transcription complex subunit 6 like | CNOT6L | -1.2 | -1.4 | -1.1 | -1.2 | -1.5 | No |
| 381 | CCR4-NOT transcription complex subunit 7 | CNOT7 | -6.8 | -0.8 | -0.6 | -1.7 | -0.6 | No |
| 382 | canopy FGF signaling regulator 2 | CNPY2 | 1.6 | 1.9 | 2.3 | 3.1 | 2.6 | No |
| 383 | consortin, connexin sorting protein | CNST | -1.5 | -1.2 | -1.5 | -0.9 | -1.7 | No |
| 384 | component of oligomeric golgi complex 2 | COG2 | 0.4 | 0.6 | 0.1 | 0.2 | 0.1 | No |
| 385 | collagen type XI alpha 2 chain | COL11A2 | -1.9 | -1.5 | 1.7 | 4.2 | 4.6 | No |
| 386 | collagen type XVII alpha 1 chain | COL17A1 | -2.6 | -3.4 | -2.1 | -3.5 | -2.1 | No |
| 387 | collagen type I alpha 1 chain | COL1A1 | -0.7 | -0.7 | 0.9 | 2.4 | 2.6 | No |
| 388 | collagen type IV alpha 5 chain | COL4A5 | 0.4 | 0.3 | 1.8 | 1.2 | 1.8 | No |
| 389 | catechol-O-methyltransferase | COMT | -1.7 | -2.8 | -0.2 | 0.3 | 0.5 | No |
| 390 | coatomer protein complex subunit beta 1 | COPB1 | 0.8 | 0.7 | 0.6 | 0.6 | 0.4 | No |
| 391 | coatomer protein complex subunit beta 2 | COPB2 | 1.0 | 0.9 | 0.9 | 0.9 | 0.8 | No |
| 392 | COP9 signalosome subunit 5 | COPS5 | 1.0 | 0.9 | 0.9 | 0.8 | 0.4 | No |
| 393 | coatomer protein complex subunit zeta 1 | COPZ1 | 1.1 | 1.2 | 0.7 | 0.8 | 0.4 | No |
| 394 | cytochrome c oxidase assembly factor heme A:farnesyltransferase COX10 | COX10 | 1.0 | 1.2 | 0.8 | 0.7 | 0.5 | No |
| 395 | cytochrome c oxidase subunit 5B | COX5B | -0.7 | 0.1 | -0.3 | -1.6 | -0.5 | No |
| 396 | cytochrome c oxidase subunit 6A1 | COX6A1 | 0.9 | 1.6 | 1.0 | 1.0 | 0.8 | No |
| 397 | carboxypeptidase M | CPM | 0.7 | 0.6 | 1.5 | 2.0 | 1.9 | No |
| 398 | copine 2 | CPNE2 | 0.2 | 0.3 | 1.3 | 2.6 | 3.0 | No |
| 399 | calcineurin like phosphoesterase domain containing 1 | CPPED1 | 0.0 | 0.6 | 0.5 | 1.0 | 0.1 | No |
| 400 | cleavage and polyadenylation specific factor 2 | CPSF2 | 1.6 | 1.4 | 1.0 | 0.8 | 0.6 | No |
| 401 | cleavage and polyadenylation specific factor 3 | CPSF3 | 1.2 | 1.3 | 0.8 | 0.8 | 0.5 | No |
| 402 | carnitine palmitoyltransferase 1B | CPT1B | -0.1 | -1.1 | -0.6 | -2.0 | 0.0 | No |
| 403 | cellular retinoic acid binding protein 2 | CRABP2 | 1.0 | 0.0 | 1.9 | 2.6 | 1.8 | No |
| 404 | carnitine O-acetyltransferase | CRAT | -0.7 | -0.7 | -0.3 | -0.7 | -0.9 | No |
| 405 | cereblon | CRBN | -0.8 | -0.6 | -0.4 | -0.1 | -0.5 | No |
| 406 | cAMP responsive element binding protein 3 like 2 | CREB3L2 | -1.0 | -1.4 | -0.9 | -0.7 | -0.3 | No |
| 407 | cysteine rich with EGF like domains 2 | CRELD2 | 0.5 | 0.9 | 0.6 | 1.6 | 1.5 | No |
| 408 | corticotropin releasing hormone binding protein | CRHBP | -0.3 | -1.5 | -1.1 | -2.4 | -1.8 | No |
| 409 | cardiolipin synthase 1 | CRLS1 | 0.8 | 1.3 | 0.6 | 0.3 | -0.8 | No |
| 410 | carnitine O-octanoyltransferase | CROT | -0.7 | -1.0 | 0.4 | -0.1 | -0.6 | No |
| 411 | cartilage associated protein | CRTAP | 1.8 | 1.6 | 2.0 | 2.7 | -6.3 | No |
| 412 | crystallin beta B3 | CRYBB3 | -1.2 | -1.1 | -1.6 | -0.8 | -0.5 | No |
| 413 | chromosome segregation 1 like | CSE1L | 1.0 | 1.3 | 0.8 | 0.9 | 0.5 | No |
| 414 | cysteine and glycine rich protein 1 | CSRP1 | 1.0 | 0.8 | 0.9 | 0.6 | 0.4 | No |
| 415 | cystatin E/M | CST6 | 1.7 | 1.5 | 2.2 | 2.8 | 2.4 | No |
| 416 | catenin beta 1 | CTNNB1 | 2.1 | 1.4 | 2.4 | 2.0 | 1.8 | No |
| 417 | catenin beta interacting protein 1 | CTNNBIP1 | 1.7 | 1.7 | 1.5 | 1.8 | 1.1 | No |
| 418 | catenin beta like 1 | CTNNBL1 | 1.7 | 1.7 | 1.5 | 1.4 | 1.0 | No |
| 419 | C-X-C motif chemokine ligand 12 | CXCL12 | -1.1 | -2.6 | -2.0 | -0.9 | -1.6 | No |
| 420 | C-X-C motif chemokine receptor 4 | CXCR4 | 0.4 | 1.0 | 0.1 | -0.2 | 0.4 | No |
| 421 | cytochrome b561 family member D2 | CYB561D2 | 1.0 | 0.8 | 0.8 | 0.8 | 0.4 | No |
| 422 | cytochrome P450 family 20 subfamily A member 1 | CYP20A1 | 0.6 | 0.7 | 0.5 | 1.0 | 0.6 | No |
| 423 | cytochrome P450 family 26 subfamily A member 1 | CYP26A1 | -1.1 | -2.1 | -0.8 | 0.1 | 0.1 | No |
| 424 | cytochrome P450 family 26 subfamily C member 1 | CYP26C1 | 1.4 | -0.3 | 1.8 | 1.9 | 1.5 | No |
| 425 | Cytochrome P450, family 2, subfamily j, polypeptide 12 | CYP2J12 | -0.7 | -0.8 | -0.4 | -0.1 | -0.2 | No |
| 426 | Leukotriene-B4 omega-hydroxylase 3 | CYP4F14 | -5.4 | 1.0 | 0.6 | -14.9 | 0.5 | No |
| 427 | None | D8ERTD738E | -1.1 | -0.9 | -0.9 | -0.8 | -1.1 | No |
| 428 | dachshund family transcription factor 1 | DACH1 | 2.0 | 2.4 | 4.4 | 4.8 | 4.1 | No |
| 429 | dual adaptor of phosphotyrosine and 3-phosphoinositides 1 | DAPP1 | -0.8 | -1.1 | -0.5 | -0.9 | -0.8 | No |
| 430 | DBF4 zinc finger | DBF4 | 0.3 | 1.3 | 0.8 | 1.0 | 0.7 | No |
| 431 | DDB1 and CUL4 associated factor 13 | DCAF13 | 1.4 | 1.1 | 0.5 | 0.4 | 0.1 | No |
| 432 | DDB1 and CUL4 associated factor 4 | DCAF4 | -1.2 | -1.0 | -1.0 | -0.9 | -1.2 | No |
| 433 | deoxycytidine kinase | DCK | 1.1 | 2.5 | 1.7 | 2.0 | 1.3 | No |
| 434 | DNA cross-link repair 1B | DCLRE1B | 0.1 | 1.4 | 1.0 | 1.5 | 0.8 | No |
| 435 | decapping enzyme, scavenger | DCPS | 2.0 | 1.8 | 1.6 | 1.8 | 1.7 | No |
| 436 | dCMP deaminase | DCTD | 1.0 | 1.9 | 1.4 | 1.5 | 0.2 | No |
| 437 | damage specific DNA binding protein 2 | DDB2 | -0.3 | -0.4 | -0.5 | -0.7 | -1.4 | No |
| 438 | DNA damage inducible transcript 3 | DDIT3 | -0.8 | -1.2 | -0.8 | -0.5 | -0.8 | No |
| 439 | DExD-box helicase 21 | DDX21 | -1.0 | -1.5 | -1.5 | -2.2 | -1.5 | No |
| 440 | DEAD-box helicase 43 | DDX43 | 0.2 | 1.8 | 1.6 | 1.9 | 0.6 | No |
| 441 | DEAD-box helicase 49 | DDX49 | 1.1 | 0.5 | 0.3 | 0.1 | 0.0 | No |
| 442 | DEAD-box helicase 51 | DDX51 | 1.3 | 1.0 | 0.4 | 0.3 | 0.0 | No |
| 443 | death effector domain containing 2 | DEDD2 | -2.6 | -2.0 | -1.4 | -1.1 | -1.3 | No |
| 444 | DENN domain containing 5B | DENND5B | 0.6 | 1.7 | 2.1 | 1.2 | 0.0 | No |
| 445 | density regulated re-initiation and release factor | DENR | 1.5 | 1.8 | 0.9 | 1.0 | 0.8 | No |
| 446 | derlin 2 | DERL2 | 0.8 | 1.1 | 0.7 | 0.7 | 0.9 | No |
| 447 | desumoylating isopeptidase 2 | DESI2 | 0.7 | 0.8 | 0.8 | 1.1 | 0.7 | No |
| 448 | Dexi homolog | DEXI | -1.4 | -0.4 | -0.9 | -0.8 | -0.7 | No |
| 449 | diacylglycerol kinase alpha | DGKA | -1.1 | -0.8 | -0.4 | -0.6 | -0.6 | No |
| 450 | diablo IAP-binding mitochondrial protein | DIABLO | 1.0 | 1.1 | 1.0 | 0.9 | 0.7 | No |
| 451 | UTP25, small subunit processor component | DIEXF | 1.5 | 0.8 | 1.0 | 0.9 | 0.5 | No |
| 452 | DIS3 like exosome 3'-5' exoribonuclease | DIS3L | -0.3 | -0.8 | -0.8 | -1.7 | -0.7 | No |
| 453 | dyskerin pseudouridine synthase 1 | DKC1 | 2.3 | 2.0 | 1.6 | 1.1 | 0.8 | No |
| 454 | DLG associated protein 5 | DLGAP5 | 0.2 | 2.2 | 2.0 | 2.8 | 2.5 | No |
| 455 | distal-less homeobox 3 | DLX3 | -1.1 | -0.6 | -1.0 | -0.4 | -0.3 | No |
| 456 | DMRT like family A2 | DMRTA2 | -1.0 | -1.6 | -0.8 | -1.5 | -0.5 | No |
| 457 | DnaJ heat shock protein family (Hsp40) member C1 | DNAJC1 | 1.3 | 1.5 | 1.0 | 1.4 | 0.8 | No |
| 458 | DnaJ heat shock protein family (Hsp40) member C2 | DNAJC2 | 1.4 | 1.7 | 0.8 | 0.6 | 0.1 | No |
| 459 | DnaJ heat shock protein family (Hsp40) member C25 | DNAJC25 | 1.2 | 0.5 | 1.2 | 0.9 | 0.7 | No |
| 460 | dynein axonemal light intermediate chain 1 | DNALI1 | 1.6 | 1.1 | 0.6 | -0.1 | 0.6 | No |
| 461 | DNA methyltransferase 1 | DNMT1 | 1.2 | 2.1 | 1.2 | 1.0 | 0.8 | No |
| 462 | double PHD fingers 2 | DPF2 | 0.9 | 0.8 | 1.1 | 0.8 | 0.6 | No |
| 463 | dipeptidyl peptidase 3 | DPP3 | 1.3 | 1.7 | 1.2 | 1.2 | 0.8 | No |
| 464 | dpy-19 like 2 | DPY19L2 | 1.1 | 0.6 | 1.7 | 1.8 | 1.4 | No |
| 465 | dpy-30, histone methyltransferase complex regulatory subunit | DPY30 | 0.7 | 1.2 | 1.0 | 1.2 | 0.9 | No |
| 466 | DNA damage regulated autophagy modulator 1 | DRAM1 | 1.2 | 0.7 | 1.6 | 1.7 | 1.0 | No |
| 467 | DNA replication and sister chromatid cohesion 1 | DSCC1 | 1.5 | 2.9 | 1.8 | 2.4 | 2.0 | No |
| 468 | denticleless E3 ubiquitin protein ligase homolog | DTL | 1.4 | 2.8 | 2.4 | 2.6 | 2.2 | No |
| 469 | DTW domain containing 1 | DTWD1 | 0.9 | 0.9 | 0.5 | 0.6 | 0.6 | No |
| 470 | deoxythymidylate kinase | DTYMK | 1.8 | 2.6 | 2.0 | 2.7 | 2.4 | No |
| 471 | dual specificity phosphatase 22 | DUSP22 | -0.6 | -0.7 | -0.3 | -0.3 | -0.5 | No |
| 472 | dual specificity phosphatase 4 | DUSP4 | 4.0 | 2.9 | 2.7 | 1.7 | 2.1 | No |
| 473 | dishevelled segment polarity protein 1 | DVL1 | -1.0 | -1.8 | -0.9 | -1.2 | -1.2 | No |
| 474 | epithelial cell transforming 2 | ECT2 | 1.1 | 2.2 | 2.5 | 2.8 | 2.0 | No |
| 475 | eukaryotic translation elongation factor 1 alpha 1 | EEF1A1 | -0.3 | -1.2 | -1.3 | -1.9 | -2.3 | No |
| 476 | eukaryotic elongation factor 2 kinase | EEF2K | -0.4 | -1.2 | -1.1 | -0.5 | -1.3 | No |
| 477 | elongation factor Tu GTP binding domain containing 2 | EFTUD2 | 1.7 | 1.3 | 1.2 | 0.8 | 0.3 | No |
| 478 | early growth response 1 | EGR1 | -0.5 | -0.5 | -2.0 | -1.2 | -1.7 | No |
| 479 | early growth response 2 | EGR2 | -1.1 | -1.1 | -2.8 | -1.5 | -3.4 | No |
| 480 | eukaryotic translation initiation factor 2B subunit gamma | EIF2B3 | 1.2 | 1.1 | 0.8 | 1.3 | 0.6 | No |
| 481 | eukaryotic translation initiation factor 2D | EIF2D | 0.8 | 1.0 | 0.4 | 0.6 | 0.4 | No |
| 482 | eukaryotic translation initiation factor 2 subunit beta | EIF2S2 | 1.2 | 1.3 | 0.6 | 0.4 | 0.0 | No |
| 483 | eukaryotic translation initiation factor 3 subunit C | EIF3C | -0.3 | -0.6 | -0.8 | -1.2 | -1.1 | No |
| 484 | eukaryotic translation initiation factor 3 subunit D | EIF3D | -0.7 | -0.9 | -0.9 | -1.0 | -0.9 | No |
| 485 | eukaryotic translation initiation factor 3 subunit G | EIF3G | -0.6 | -0.8 | -1.0 | -1.0 | -0.9 | No |
| 486 | eukaryotic translation initiation factor 3 subunit I | EIF3I | -0.7 | -1.1 | -0.8 | -1.0 | -0.9 | No |
| 487 | eukaryotic translation initiation factor 3 subunit M | EIF3M | -0.6 | -1.0 | -1.0 | -1.3 | -1.0 | No |
| 488 | eukaryotic translation initiation factor 4E | EIF4E | 1.3 | 1.2 | 0.8 | 0.8 | 0.8 | No |
| 489 | eukaryotic translation initiation factor 4 gamma 2 | EIF4G2 | 1.0 | 0.9 | 1.1 | 1.2 | 0.8 | No |
| 490 | eukaryotic translation initiation factor 5 | EIF5 | 0.7 | 1.0 | 0.6 | 0.7 | 0.3 | No |
| 491 | ELAV like RNA binding protein 1 | ELAVL1 | 1.7 | 1.4 | 1.6 | 0.9 | 0.1 | No |
| 492 | E74 like ETS transcription factor 2 | ELF2 | -0.7 | -0.8 | -0.5 | -0.1 | -0.8 | No |
| 493 | elongation factor for RNA polymerase II | ELL | -0.9 | -0.9 | -1.1 | -0.4 | -1.0 | No |
| 494 | ER membrane protein complex subunit 10 | EMC10 | 0.7 | 0.9 | 0.6 | 0.4 | 0.2 | No |
| 495 | ER membrane protein complex subunit 6 | EMC6 | 1.0 | 1.3 | 0.8 | 0.8 | 0.4 | No |
| 496 | echinoderm microtubule associated protein like 1 | EML1 | -5.6 | -2.1 | -6.6 | -11.2 | -1.1 | No |
| 497 | epithelial membrane protein 1 | EMP1 | 0.8 | 0.7 | -8.4 | 0.9 | 0.8 | No |
| 498 | empty spiracles homeobox 2 | EMX2 | 0.4 | 0.4 | 1.1 | 0.9 | 1.8 | No |
| 499 | endonuclease G | ENDOG | 1.5 | 1.6 | 0.7 | 0.2 | 0.0 | No |
| 500 | ectonucleotide pyrophosphatase/phosphodiesterase 1 | ENPP1 | 0.5 | 0.3 | 1.1 | 1.0 | 0.7 | No |
| 501 | ectonucleoside triphosphate diphosphohydrolase 1 | ENTPD1 | 1.4 | 0.8 | 1.1 | 0.7 | 1.0 | No |
| 502 | ectonucleoside triphosphate diphosphohydrolase 8 | ENTPD8 | -2.7 | -0.7 | -1.6 | -0.7 | -1.2 | No |
| 503 | endothelial PAS domain protein 1 | EPAS1 | -2.2 | -1.9 | -1.9 | -2.4 | -16.7 | No |
| 504 | ependymin related 1 | EPDR1 | -1.7 | -1.9 | -0.9 | -0.5 | -0.4 | No |
| 505 | EPH receptor A7 | EPHA7 | 1.2 | 1.9 | 0.8 | 0.2 | 0.8 | No |
| 506 | EPH receptor B4 | EPHB4 | 1.3 | 1.1 | 0.6 | 0.6 | 1.3 | No |
| 507 | erb-b2 receptor tyrosine kinase 3 | ERBB3 | -1.3 | -1.1 | -1.3 | -1.4 | -1.3 | No |
| 508 | establishment of sister chromatid cohesion N-acetyltransferase 2 | ESCO2 | 1.2 | 3.0 | 1.9 | 2.6 | 1.8 | No |
| 509 | epithelial splicing regulatory protein 1 | ESRP1 | -1.4 | -0.9 | -1.5 | -1.8 | -1.3 | No |
| 510 | ETHE1, persulfide dioxygenase | ETHE1 | -1.1 | -5.2 | -5.6 | -1.9 | 0.0 | No |
| 511 | ETS variant 4 | ETV4 | 0.8 | 1.5 | 3.7 | 4.1 | 3.8 | No |
| 512 | envoplakin | EVPL | -1.2 | -1.5 | -1.8 | -2.6 | -1.5 | No |
| 513 | exonuclease 1 | EXO1 | 0.8 | 2.4 | 1.8 | 2.6 | 1.9 | No |
| 514 | exosome component 5 | EXOSC5 | 1.8 | 1.8 | 1.0 | 1.0 | 0.8 | No |
| 515 | exosome component 6 | EXOSC6 | 1.3 | 1.5 | 0.9 | 0.8 | 1.0 | No |
| 516 | exosome component 9 | EXOSC9 | 2.1 | 2.2 | 1.3 | 1.0 | 0.6 | No |
| 517 | EYA transcriptional coactivator and phosphatase 1 | EYA1 | -0.6 | -1.3 | -0.5 | -0.7 | -0.7 | No |
| 518 | enhancer of zeste 1 polycomb repressive complex 2 subunit | EZH1 | -1.2 | -0.7 | -0.4 | -0.9 | -1.1 | No |
| 519 | enhancer of zeste 2 polycomb repressive complex 2 subunit | EZH2 | 0.7 | 1.0 | 0.8 | 0.9 | 0.5 | No |
| 520 | F2R like trypsin receptor 1 | F2RL1 | -0.6 | -0.6 | -0.9 | -1.2 | -1.2 | No |
| 521 | fatty acid binding protein 7 | FABP7 | 0.6 | 1.4 | 2.4 | 2.2 | 1.4 | No |
| 522 | fatty acid binding protein 9 | FABP9 | 3.1 | 2.9 | 2.6 | 0.7 | -0.1 | No |
| 523 | Fas associated factor family member 2 | FAF2 | 0.4 | 0.5 | 0.2 | 0.0 | 0.2 | No |
| 524 | fumarylacetoacetate hydrolase domain containing 1 | FAHD1 | 0.7 | 1.0 | 0.7 | 1.0 | 0.7 | No |
| 525 | family with sequence similarity 102 member B | FAM102B | -1.5 | -1.3 | -1.2 | -1.6 | -1.3 | No |
| 526 | family with sequence similarity 114 member A1 | FAM114A1 | 0.7 | 0.7 | 1.5 | 2.9 | 2.6 | No |
| 527 | family with sequence similarity 129 member A | FAM129A | -1.6 | -2.2 | -2.6 | -2.3 | -1.7 | No |
| 528 | family with sequence similarity 151 member A | FAM151A | -1.4 | -1.0 | -0.9 | -0.6 | -1.8 | No |
| 529 | family with sequence similarity 172 member A | FAM172A | -0.6 | -1.3 | -0.5 | -0.7 | -0.6 | No |
| 530 | family with sequence similarity 173 member B | FAM173B | 0.9 | 1.3 | 0.5 | 0.8 | 0.3 | No |
| 531 | family with sequence similarity 174 member B | FAM174B | -1.2 | -2.5 | -1.9 | -0.9 | -0.6 | No |
| 532 | family with sequence similarity 241 member A | FAM241A | -0.9 | -1.6 | -1.2 | -1.2 | -1.2 | No |
| 533 | family with sequence similarity 76 member B | FAM76B | 0.7 | 0.7 | 0.6 | 0.5 | 0.5 | No |
| 534 | FA complementation group G | FANCG | 0.9 | 2.5 | 1.1 | 1.4 | 0.9 | No |
| 535 | FA complementation group I | FANCI | 1.2 | 2.6 | 1.5 | 1.9 | 1.8 | No |
| 536 | FA complementation group L | FANCL | 0.9 | 1.9 | 0.8 | 1.1 | 1.1 | No |
| 537 | fatty acyl-CoA reductase 1 | FAR1 | -0.7 | -0.5 | -0.9 | -1.8 | -2.1 | No |
| 538 | phenylalanyl-tRNA synthetase subunit beta | FARSB | 1.3 | 0.9 | 0.6 | 0.4 | 0.1 | No |
| 539 | fibrillarin like 1 | FBLL1 | 1.8 | 1.7 | 0.8 | 0.8 | 0.5 | No |
| 540 | F-box protein 30 | FBXO30 | 1.0 | 0.7 | 1.4 | 1.4 | 0.9 | No |
| 541 | F-box protein 5 | FBXO5 | 0.7 | 2.9 | 2.1 | 2.8 | 2.7 | No |
| 542 | flap structure-specific endonuclease 1 | FEN1 | 1.5 | 2.6 | 1.8 | 1.9 | 1.2 | No |
| 543 | FES proto-oncogene, tyrosine kinase | FES | -4.0 | -1.0 | -0.4 | -11.8 | -12.8 | No |
| 544 | fibroblast growth factor 16 | FGF16 | -2.4 | -2.4 | -1.0 | -0.3 | -1.1 | No |
| 545 | fibroblast growth factor receptor 2 | FGFR2 | -1.1 | -1.3 | -0.7 | 0.0 | 0.0 | No |
| 546 | FK506 binding protein 10 | FKBP10 | 0.5 | 0.8 | 2.3 | 3.8 | 3.3 | No |
| 547 | FK506 binding protein 14 | FKBP14 | 0.9 | 1.0 | 2.1 | 4.2 | 3.8 | No |
| 548 | FK506 binding protein 2 | FKBP2 | 1.3 | 1.3 | 1.1 | 1.5 | 1.0 | No |
| 549 | FK506 binding protein 9 | FKBP9 | 1.5 | 1.6 | 3.4 | 5.2 | 4.5 | No |
| 550 | fibronectin 1 | FN1 | 3.6 | 5.1 | 5.2 | 4.5 | 3.5 | No |
| 551 | forkhead box I1 | FOXI1 | -0.8 | -1.7 | -2.4 | -3.3 | -2.6 | No |
| 552 | forkhead box K1 | FOXK1 | -0.6 | -1.0 | -0.6 | -0.6 | -0.5 | No |
| 553 | forkhead box N1 | FOXN1 | -2.2 | -2.1 | -1.8 | -2.7 | -2.7 | No |
| 554 | forkhead box Q1 | FOXQ1 | -2.4 | -2.7 | -2.6 | -1.8 | -1.2 | No |
| 555 | follistatin like 3 | FSTL3 | -0.2 | -1.3 | -1.1 | -1.1 | 0.0 | No |
| 556 | alpha-L-fucosidase 2 | FUCA2 | -1.3 | -0.9 | -0.5 | -0.4 | -0.2 | No |
| 557 | fucosyltransferase 9 | FUT9 | -1.0 | -2.1 | -1.8 | -1.0 | 0.8 | No |
| 558 | frizzled class receptor 3 | FZD3 | -1.6 | -0.9 | -1.2 | -1.3 | -1.6 | No |
| 559 | frizzled class receptor 8 | FZD8 | -0.8 | -1.5 | -0.4 | -1.1 | -0.1 | No |
| 560 | G2/M-phase specific E3 ubiquitin protein ligase | G2E3 | 0.5 | 2.2 | 2.8 | 3.4 | 3.2 | No |
| 561 | GABA type A receptor associated protein like 2 | GABARAPL2 | -0.8 | -1.1 | -0.8 | -1.0 | -0.6 | No |
| 562 | glutamate decarboxylase 2 | GAD2 | -1.7 | 0.0 | -1.1 | -0.7 | -3.0 | No |
| 563 | galactokinase 1 | GALK1 | 0.7 | 1.7 | 1.5 | 2.2 | 1.6 | No |
| 564 | GATA zinc finger domain containing 1 | GATAD1 | 1.1 | 1.1 | 0.9 | 0.7 | 0.6 | No |
| 565 | GATS protein like 2 | GATSL2 | -1.3 | -0.9 | -0.6 | -0.5 | -1.3 | No |
| 566 | glycine C-acetyltransferase | GCAT | 1.2 | 1.4 | 1.2 | 0.8 | 0.6 | No |
| 567 | glucagon | GCG | 6.3 | 4.4 | 1.7 | 2.4 | 4.3 | No |
| 568 | glutamate-cysteine ligase modifier subunit | GCLM | 2.6 | 2.0 | 1.9 | 1.8 | 1.9 | No |
| 569 | glial cells missing homolog 2 | GCM2 | -0.3 | -7.3 | -1.0 | -0.5 | -15.0 | No |
| 570 | glucosaminyl (N-acetyl) transferase 4, core 2 | GCNT4 | -0.3 | -0.7 | -1.3 | -1.4 | -0.7 | No |
| 571 | guanine deaminase | GDA | 2.9 | 1.7 | 1.8 | -0.4 | 0.9 | No |
| 572 | glycerophosphodiester phosphodiesterase 1 | GDE1 | 0.6 | 0.9 | 0.7 | 0.8 | 0.7 | No |
| 573 | glycerophosphodiester phosphodiesterase domain containing 3 | GDPD3 | 0.7 | 1.1 | 0.7 | 0.0 | 0.3 | No |
| 574 | GTP binding protein overexpressed in skeletal muscle | GEM | 3.3 | 3.6 | 1.4 | 2.1 | 2.1 | No |
| 575 | GEN1, Holliday junction 5' flap endonuclease | GEN1 | 1.1 | 1.2 | 0.5 | 0.6 | 0.8 | No |
| 576 | growth factor, augmenter of liver regeneration | GFER | 1.4 | 1.6 | 0.8 | 1.2 | 0.9 | No |
| 577 | G elongation factor mitochondrial 1 | GFM1 | 1.9 | 1.5 | 0.8 | 0.6 | 0.3 | No |
| 578 | G elongation factor mitochondrial 2 | GFM2 | 0.7 | 1.5 | 0.9 | 1.1 | 0.4 | No |
| 579 | glucose-fructose oxidoreductase domain containing 2 | GFOD2 | -1.1 | -1.1 | -0.8 | -0.6 | -0.7 | No |
| 580 | GINS complex subunit 3 | GINS3 | 1.1 | 2.6 | 1.8 | 2.1 | 1.7 | No |
| 581 | GIT ArfGAP 2 | GIT2 | -0.7 | -0.7 | -0.3 | -0.1 | -0.9 | No |
| 582 | gap junction protein beta 3 | GJB3 | -0.6 | -0.8 | -0.8 | -1.0 | -1.0 | No |
| 583 | glucocorticoid induced 1 | GLCCI1 | -0.7 | -0.5 | -0.8 | -0.5 | -0.6 | No |
| 584 | GLI family zinc finger 2 | GLI2 | -1.1 | -1.1 | -0.6 | -1.6 | -1.3 | No |
| 585 | GLI pathogenesis related 2 | GLIPR2 | 3.1 | 3.6 | 2.8 | 4.0 | 4.2 | No |
| 586 | glyoxalase domain containing 4 | GLOD4 | 1.1 | 0.9 | 1.0 | 0.9 | 0.4 | No |
| 587 | glutaredoxin | GLRX | 0.3 | 1.0 | 0.8 | 0.8 | 0.7 | No |
| 588 | glutaredoxin 5 | GLRX5 | 0.7 | 1.0 | 0.8 | 0.8 | 0.8 | No |
| 589 | glycosyltransferase 8 domain containing 1 | GLT8D1 | 0.8 | 0.7 | 0.8 | 1.5 | 1.3 | No |
| 590 | None | GM14399 | 0.6 | 1.0 | 0.7 | 1.4 | 1.1 | No |
| 591 | None | GM17018 | 1.3 | 1.4 | 1.0 | 1.2 | 1.0 | No |
| 592 | None | GM2022 | 1.1 | 1.0 | 0.7 | 0.7 | 0.3 | No |
| 593 | None | GM20716 | 1.0 | 0.8 | 0.8 | 1.4 | 1.1 | No |
| 594 | None | GM21708 | -0.8 | -0.6 | -0.7 | -0.1 | -1.0 | No |
| 595 | None | GM21972 | 1.4 | 1.4 | 1.3 | 1.2 | 0.8 | No |
| 596 | None | GM28308 | 0.0 | 1.5 | 2.1 | 1.8 | 2.2 | No |
| 597 | None | GM43552 | -1.8 | -2.0 | -1.2 | -0.8 | -1.1 | No |
| 598 | None | GM4846 | -2.3 | -1.2 | -1.2 | -1.9 | -1.9 | No |
| 599 | None | GM4907 | 1.0 | 0.9 | 1.6 | 1.5 | 1.1 | No |
| 600 | None | GM5145 | 1.2 | 1.2 | 0.9 | 0.8 | 0.5 | No |
| 601 | None | GM5225 | 0.1 | 0.7 | 1.0 | 1.1 | 0.6 | No |
| 602 | None | GM7879 | 2.3 | 1.1 | 2.1 | 1.5 | -0.4 | No |
| 603 | None | GM9774 | 1.0 | 1.1 | 0.8 | 0.8 | 0.4 | No |
| 604 | G protein subunit gamma 3 | GNG3 | -2.5 | -2.2 | -1.8 | 0.2 | 0.3 | No |
| 605 | glutamic-oxaloacetic transaminase 2 | GOT2 | 1.4 | 1.6 | 0.8 | 0.8 | 0.6 | No |
| 606 | glucose phosphate isomerase 1 | GPI1 | 0.9 | 1.0 | 0.5 | 0.2 | 0.0 | No |
| 607 | glycoprotein M6A | GPM6A | -2.7 | -3.3 | -2.7 | -1.4 | -1.8 | No |
| 608 | G protein-coupled receptor 157 | GPR157 | -1.0 | -0.6 | -0.5 | -0.4 | -0.7 | No |
| 609 | G protein-coupled receptor 182 | GPR182 | -0.8 | -0.9 | -0.9 | -1.0 | -0.2 | No |
| 610 | G protein signaling modulator 1 | GPSM1 | 1.4 | 0.8 | 1.2 | 0.5 | 0.5 | No |
| 611 | glutathione peroxidase 2 | GPX2 | 0.8 | 1.9 | 0.4 | 1.1 | 1.9 | No |
| 612 | glutathione peroxidase 8 (putative) | GPX8 | 0.3 | 1.1 | 1.4 | 2.4 | 2.2 | No |
| 613 | growth factor receptor bound protein 10 | GRB10 | -2.1 | -1.1 | -2.1 | -2.0 | -2.4 | No |
| 614 | glyoxylate and hydroxypyruvate reductase | GRHPR | -0.6 | -0.8 | -0.6 | -0.5 | -0.5 | No |
| 615 | glutamate ionotropic receptor AMPA type subunit 1 | GRIA1 | -2.4 | -2.0 | -2.0 | -1.3 | -4.5 | No |
| 616 | GrpE like 1, mitochondrial | GRPEL1 | 1.4 | 1.1 | 0.7 | 1.3 | 0.4 | No |
| 617 | glycine/arginine rich protein 1 | GRRP1 | 3.4 | 2.7 | 3.8 | 3.5 | 3.1 | No |
| 618 | glutathione synthetase | GSS | 1.2 | 0.6 | 0.6 | 0.3 | 0.4 | No |
| 619 | glutathione S-transferase omega 1 | GSTO1 | -1.7 | -0.5 | -1.6 | -1.2 | -0.7 | No |
| 620 | general transcription factor IIH subunit 2 | GTF2H2 | 0.8 | 0.7 | 0.7 | 1.0 | 0.6 | No |
| 621 | general transcription factor IIH subunit 4 | GTF2H4 | 0.9 | 0.9 | 1.0 | 1.4 | 0.7 | No |
| 622 | general transcription factor IIIC subunit 2 | GTF3C2 | 1.5 | 0.9 | 0.6 | 1.2 | -0.9 | No |
| 623 | GTP binding protein 3, mitochondrial | GTPBP3 | 1.3 | 0.9 | 0.7 | 0.9 | 0.4 | No |
| 624 | H2A histone family member X | H2AFX | 1.6 | 3.0 | 2.1 | 2.8 | 2.3 | No |
| 625 | histocompatibility 2, class II, locus DMa | H2-DMA | -2.8 | -1.8 | -1.8 | -1.5 | -2.4 | No |
| 626 | hydroxyacyl-CoA dehydrogenase trifunctional multienzyme complex subunit alpha | HADHA | 1.4 | 1.4 | 1.1 | 0.9 | 0.7 | No |
| 627 | histidine ammonia-lyase | HAL | -0.4 | -1.3 | -1.6 | -2.9 | -2.8 | No |
| 628 | histone acetyltransferase 1 | HAT1 | 0.8 | 1.1 | 0.7 | 1.0 | 0.7 | No |
| 629 | HAUS augmin like complex subunit 4 | HAUS4 | 2.1 | 3.0 | 2.3 | 2.3 | 1.8 | No |
| 630 | HAUS augmin like complex subunit 5 | HAUS5 | 0.9 | 2.7 | 1.8 | 3.6 | 1.7 | No |
| 631 | HMG-box transcription factor 1 | HBP1 | -1.1 | -1.5 | -1.6 | -0.5 | -15.3 | No |
| 632 | host cell factor C2 | HCFC2 | 1.1 | 0.8 | 0.7 | 1.6 | 0.9 | No |
| 633 | histone deacetylase 9 | HDAC9 | -1.4 | -1.4 | -1.2 | -2.8 | -0.4 | No |
| 634 | HD domain containing 3 | HDDC3 | 0.8 | 0.9 | 1.0 | 1.1 | 0.6 | No |
| 635 | HDGF like 2 | HDGFL2 | 1.1 | 1.3 | 1.0 | 0.5 | 0.3 | No |
| 636 | hdc homolog, cell cycle regulator | HECA | -1.8 | -1.6 | -1.3 | -0.8 | -1.1 | No |
| 637 | helicase, lymphoid specific | HELLS | 2.0 | 2.4 | 1.7 | 1.6 | 0.9 | No |
| 638 | hes family bHLH transcription factor 7 | HES7 | -0.7 | -0.7 | -1.5 | -2.0 | -0.6 | No |
| 639 | hes related family bHLH transcription factor with YRPW motif 1 | HEY1 | -1.6 | -1.1 | -0.8 | -0.4 | -0.6 | No |
| 640 | homogentisate 1,2-dioxygenase | HGD | -1.2 | -2.0 | -1.9 | -1.6 | -1.2 | No |
| 641 | 3-hydroxyisobutyrate dehydrogenase | HIBADH | 1.1 | 1.1 | 0.8 | 0.8 | 0.5 | No |
| 642 | huntingtin interacting protein 1 | HIP1 | 0.6 | -0.1 | 0.6 | 1.4 | 0.5 | No |
| 643 | homeodomain interacting protein kinase 3 | HIPK3 | -0.9 | -0.8 | -0.9 | -1.0 | -0.8 | No |
| 644 | HIRA interacting protein 3 | HIRIP3 | 0.7 | 1.4 | 1.0 | 2.1 | 1.1 | No |
| 645 | hexokinase 1 | HK1 | 1.2 | 1.0 | 0.9 | 0.5 | 0.4 | No |
| 646 | hydroxymethylbilane synthase | HMBS | 1.7 | 1.8 | 1.1 | 0.9 | 0.1 | No |
| 647 | high mobility group box 3 | HMGB3 | 0.8 | 0.8 | 0.7 | 1.1 | 0.0 | No |
| 648 | hyaluronan mediated motility receptor | HMMR | -4.6 | 2.4 | 1.7 | 3.6 | 0.7 | No |
| 649 | heme oxygenase 2 | HMOX2 | -1.6 | -1.4 | -1.2 | -0.8 | -1.0 | No |
| 650 | hepatocyte nuclear factor 4 gamma | HNF4G | 0.5 | 2.1 | 1.9 | 0.9 | 2.7 | No |
| 651 | heterogeneous nuclear ribonucleoprotein L | HNRNPL | 1.3 | 1.1 | 0.8 | 0.8 | 0.5 | No |
| 652 | hook microtubule tethering protein 1 | HOOK1 | -1.3 | -1.4 | -1.8 | -2.2 | -1.9 | No |
| 653 | homeobox C11 | HOXC11 | -0.5 | -1.2 | -1.6 | -1.6 | -1.5 | No |
| 654 | 4-hydroxyphenylpyruvate dioxygenase | HPD | -2.0 | -1.9 | -1.9 | -1.7 | -1.1 | No |
| 655 | histamine receptor H3 | HRH3 | 2.3 | 2.3 | 5.2 | 6.6 | 4.6 | No |
| 656 | hydroxysteroid 17-beta dehydrogenase 10 | HSD17B10 | 1.2 | 1.5 | 0.6 | 0.6 | 0.5 | No |
| 657 | hydroxysteroid 17-beta dehydrogenase 12 | HSD17B12 | 1.1 | 0.9 | 1.0 | 0.9 | 0.3 | No |
| 658 | hematopoietic SH2 domain containing | HSH2D | -1.2 | -1.1 | -2.2 | -2.0 | -1.3 | No |
| 659 | heat shock protein 90 beta family member 1 | HSP90B1 | 1.7 | 1.3 | 1.5 | 1.1 | 1.5 | No |
| 660 | heat shock protein family A (Hsp70) member 14 | HSPA14 | 1.5 | 1.3 | 0.6 | 0.4 | 0.2 | No |
| 661 | heat shock protein family A (Hsp70) member 4 | HSPA4 | 1.1 | 0.8 | 0.6 | 0.5 | 0.2 | No |
| 662 | 5-hydroxytryptamine receptor 2C | HTR2C | 0.1 | -1.2 | -1.4 | -1.7 | -0.8 | No |
| 663 | hydroxypyruvate isomerase (putative) | HYI | -1.0 | -1.8 | -0.6 | -0.7 | -2.0 | No |
| 664 | hypoxia up-regulated 1 | HYOU1 | 2.0 | 1.4 | 1.9 | 1.6 | 1.6 | No |
| 665 | huntingtin interacting protein K | HYPK | 1.2 | 1.4 | 0.7 | 0.8 | 0.4 | No |
| 666 | isoamyl acetate hydrolyzing esterase 1 (putative) | IAH1 | 1.0 | 1.3 | 1.3 | 1.7 | 0.9 | No |
| 667 | inhibitor of DNA binding 4, HLH protein | ID4 | -0.9 | -0.6 | -1.1 | -0.9 | -0.6 | No |
| 668 | interferon related developmental regulator 1 | IFRD1 | 0.7 | 0.9 | 0.5 | 0.3 | 0.4 | No |
| 669 | interferon related developmental regulator 2 | IFRD2 | 1.1 | 1.3 | 0.9 | 0.9 | 0.6 | No |
| 670 | intraflagellar transport 52 | IFT52 | 1.0 | 1.2 | 1.3 | 1.9 | 1.4 | No |
| 671 | intraflagellar transport 57 | IFT57 | 0.1 | 0.8 | 0.6 | 1.1 | 1.2 | No |
| 672 | insulin like growth factor 1 receptor | IGF1R | 0.2 | 2.0 | 2.4 | 3.9 | 3.3 | No |
| 673 | insulin like growth factor 2 | IGF2 | 1.1 | 0.2 | 2.8 | 3.1 | 2.4 | No |
| 674 | insulin like growth factor 2 mRNA binding protein 1 | IGF2BP1 | 3.8 | 7.4 | 7.4 | 13.7 | 14.0 | No |
| 675 | insulin like growth factor 2 mRNA binding protein 3 | IGF2BP3 | 0.3 | 0.6 | 1.0 | 1.2 | 0.5 | No |
| 676 | insulin like growth factor binding protein 2 | IGFBP2 | -1.4 | -1.6 | -1.5 | -2.0 | -1.8 | No |
| 677 | interleukin enhancer binding factor 2 | ILF2 | 1.6 | 1.7 | 1.3 | 1.4 | 1.0 | No |
| 678 | IMP3, U3 small nucleolar ribonucleoprotein | IMP3 | 1.6 | 1.1 | 0.8 | 0.6 | 0.2 | No |
| 679 | IMP4, U3 small nucleolar ribonucleoprotein | IMP4 | 1.7 | 1.4 | 1.0 | 0.6 | 0.1 | No |
| 680 | inosine monophosphate dehydrogenase 1 | IMPDH1 | -2.8 | -1.5 | -1.7 | -1.2 | -2.4 | No |
| 681 | inhibitor of growth family member 5 | ING5 | 1.2 | 2.4 | 1.2 | 1.6 | 1.3 | No |
| 682 | inhibin subunit beta A | INHBA | 4.2 | 3.5 | 2.8 | 3.4 | 3.7 | No |
| 683 | INTS3 and NABP interacting protein | INIP | 1.3 | 1.5 | 1.0 | 1.5 | 0.8 | No |
| 684 | insulin receptor | INSR | -0.9 | -0.7 | -1.1 | -1.2 | -1.3 | No |
| 685 | integrator complex subunit 9 | INTS9 | 0.7 | 0.8 | 0.9 | 1.1 | 0.8 | No |
| 686 | interferon regulatory factor 7 | IRF7 | 1.1 | 1.0 | 1.8 | 1.0 | 1.4 | No |
| 687 | immunoglobulin superfamily containing leucine rich repeat | ISLR | -1.0 | -1.4 | -0.9 | -1.3 | -2.0 | No |
| 688 | integrin subunit alpha 5 | ITGA5 | 1.4 | 2.0 | 2.4 | 1.4 | 1.7 | No |
| 689 | integrin subunit beta 3 | ITGB3 | 1.8 | 1.8 | 1.7 | 1.4 | 1.1 | No |
| 690 | IL2 inducible T cell kinase | ITK | -0.6 | -0.7 | -1.0 | -1.2 | -1.7 | No |
| 691 | integral membrane protein 2B | ITM2B | -1.0 | -1.0 | -0.5 | -0.3 | -0.6 | No |
| 692 | integral membrane protein 2C | ITM2C | 1.3 | 1.1 | 1.5 | 1.8 | 1.3 | No |
| 693 | intersectin 2 | ITSN2 | -0.8 | -0.9 | -1.3 | -1.6 | -1.0 | No |
| 694 | influenza virus NS1A binding protein | IVNS1ABP | -2.0 | -1.0 | -1.7 | -1.6 | -1.4 | No |
| 695 | jagged 1 | JAG1 | -1.3 | -1.0 | -1.5 | -1.8 | -1.8 | No |
| 696 | Janus kinase 2 | JAK2 | -1.0 | -0.7 | -0.8 | -0.6 | -1.1 | No |
| 697 | Jun proto-oncogene, AP-1 transcription factor subunit | JUN | -0.8 | -0.9 | -1.4 | -0.9 | -0.2 | No |
| 698 | potassium channel tetramerization domain containing 12 | KCTD12 | -0.1 | -1.0 | -0.5 | -0.4 | -0.5 | No |
| 699 | KDEL motif containing 2 | KDELC2 | 6.3 | 6.8 | 7.8 | 13.7 | 0.2 | No |
| 700 | KDEL endoplasmic reticulum protein retention receptor 2 | KDELR2 | 0.7 | 0.8 | 0.6 | 1.3 | 1.1 | No |
| 701 | KH RNA binding domain containing, signal transduction associated 1 | KHDRBS1 | 0.5 | 0.2 | 0.1 | 0.3 | 0.1 | No |
| 702 | KH RNA binding domain containing, signal transduction associated 2 | KHDRBS2 | -2.0 | -2.6 | -3.1 | -1.2 | -3.0 | No |
| 703 | kinesin family member 11 | KIF11 | 1.5 | 2.5 | 3.2 | 3.0 | 2.9 | No |
| 704 | kinesin family member 18A | KIF18A | 0.7 | 2.5 | 1.6 | 2.0 | 1.9 | No |
| 705 | kinesin family member 23 | KIF23 | -4.1 | 1.8 | 1.7 | 1.8 | 1.5 | No |
| 706 | Kin17 DNA and RNA binding protein | KIN | 0.7 | 0.7 | 0.4 | 0.4 | 0.6 | No |
| 707 | Kruppel like factor 9 | KLF9 | -0.1 | -2.4 | -1.7 | -2.1 | -0.4 | No |
| 708 | kelch like family member 32 | KLHL32 | -0.7 | -0.7 | -0.5 | -0.5 | -1.9 | No |
| 709 | karyopherin subunit alpha 2 | KPNA2 | 1.8 | 3.3 | 3.5 | 4.2 | 3.8 | No |
| 710 | keratin 19 | KRT19 | -1.9 | -1.8 | -1.9 | -2.0 | -1.1 | No |
| 711 | leucine aminopeptidase 3 | LAP3 | 1.5 | 1.7 | 0.7 | 0.7 | 0.4 | No |
| 712 | large tumor suppressor kinase 1 | LATS1 | -0.4 | -5.0 | -2.6 | -14.3 | -4.1 | No |
| 713 | leukocyte cell derived chemotaxin 2 | LECT2 | 4.3 | 3.1 | 3.1 | 2.8 | 1.7 | No |
| 714 | lymphoid enhancer binding factor 1 | LEF1 | -0.2 | 0.9 | 3.2 | 2.3 | 2.3 | No |
| 715 | LEO1 homolog, Paf1/RNA polymerase II complex component | LEO1 | 0.9 | 1.0 | 0.6 | 0.4 | 0.2 | No |
| 716 | leucine rich glioma inactivated 1 | LGI1 | -9.6 | -1.3 | -1.0 | -1.7 | 0.0 | No |
| 717 | lengsin, lens protein with glutamine synthetase domain | LGSN | 2.2 | 3.1 | 2.7 | 1.9 | 2.7 | No |
| 718 | LHFPL tetraspan subfamily member 3 | LHFPL3 | 1.5 | 2.4 | 2.5 | 1.5 | 0.9 | No |
| 719 | LIM homeobox 8 | LHX8 | -1.5 | -3.0 | -3.5 | -2.5 | -2.0 | No |
| 720 | LIF receptor alpha | LIFR | 2.0 | 2.4 | 2.0 | 1.2 | 1.1 | No |
| 721 | lin-52 DREAM MuvB core complex component | LIN52 | 1.5 | 1.6 | 1.2 | 1.5 | 0.8 | No |
| 722 | lipase G, endothelial type | LIPG | 0.7 | 0.8 | 0.7 | 0.3 | 0.8 | No |
| 723 | lamin B1 | LMNB1 | 1.2 | 1.5 | 1.1 | 1.3 | 1.2 | No |
| 724 | lysyl oxidase like 1 | LOXL1 | -0.6 | -2.1 | -1.6 | -0.6 | -0.3 | No |
| 725 | lysyl oxidase like 3 | LOXL3 | -1.7 | -0.8 | -0.7 | -0.2 | 0.0 | No |
| 726 | lysophosphatidylcholine acyltransferase 4 | LPCAT4 | -1.0 | -0.7 | -1.1 | -0.8 | -0.3 | No |
| 727 | LDL receptor related protein 11 | LRP11 | -1.7 | -2.0 | -2.5 | -1.7 | -1.8 | No |
| 728 | leucine rich repeat containing 47 | LRRC47 | 1.3 | 1.0 | 0.8 | 0.0 | -0.9 | No |
| 729 | leucine rich repeat neuronal 1 | LRRN1 | 2.2 | 2.3 | 3.6 | 3.8 | 3.7 | No |
| 730 | LSM1 homolog, mRNA degradation associated | LSM1 | 1.2 | 1.8 | 0.9 | 1.3 | 0.7 | No |
| 731 | LSM12 homolog | LSM12 | 1.2 | 1.6 | 0.9 | 0.5 | 0.5 | No |
| 732 | LSM family member 14B | LSM14B | -0.8 | -0.9 | -1.1 | -0.9 | -1.1 | No |
| 733 | LSM4 homolog, U6 small nuclear RNA and mRNA degradation associated | LSM4 | 0.8 | 0.9 | 0.8 | 1.2 | 0.6 | No |
| 734 | LSM7 homolog, U6 small nuclear RNA and mRNA degradation associated | LSM7 | 1.1 | 1.6 | 0.8 | 1.0 | 0.8 | No |
| 735 | leukotriene A4 hydrolase | LTA4H | 1.3 | 1.1 | 1.3 | 1.2 | 0.6 | No |
| 736 | latexin | LXN | -1.1 | -2.0 | -2.0 | -2.1 | -1.9 | No |
| 737 | LysM domain containing 4 | LYSMD4 | 0.5 | 0.3 | 0.2 | 0.4 | 0.6 | No |
| 738 | mitotic arrest deficient 2 like 1 | MAD2L1 | 1.1 | 2.3 | 2.7 | 3.0 | 2.7 | No |
| 739 | MAGI family member, X-linked | MAGIX | -1.3 | -1.0 | -1.4 | -2.7 | -0.8 | No |
| 740 | mago homolog, exon junction complex subunit | MAGOH | 1.5 | 1.5 | 1.2 | 1.0 | 0.5 | No |
| 741 | male germ cell associated kinase | MAK | -0.4 | -0.6 | -0.4 | -0.2 | -0.4 | No |
| 742 | mesencephalic astrocyte derived neurotrophic factor | MANF | 1.8 | 1.8 | 1.7 | 1.3 | 1.7 | No |
| 743 | mitogen-activated protein kinase kinase kinase 5 | MAP3K5 | -1.5 | -1.6 | -1.4 | -1.0 | -1.0 | No |
| 744 | mitogen-activated protein kinase 14 | MAPK14 | -1.4 | -0.9 | -1.4 | -1.3 | -1.0 | No |
| 745 | mitogen-activated protein kinase 4 | MAPK4 | 1.4 | 1.6 | 3.1 | -0.4 | 2.6 | No |
| 746 | MARCKS like 1 | MARCKSL1 | 2.8 | 2.5 | 1.4 | 1.4 | 1.5 | No |
| 747 | microtubule associated serine/threonine kinase like | MASTL | 0.0 | 1.1 | 1.1 | 1.5 | 1.7 | No |
| 748 | matrilin 4 | MATN4 | -2.4 | -2.4 | -3.3 | -2.8 | -0.2 | No |
| 749 | methyl-CpG binding domain protein 3 | MBD3 | 1.3 | 0.4 | 1.9 | 1.9 | 1.5 | No |
| 750 | muscleblind like splicing regulator 3 | MBNL3 | -0.1 | -0.4 | -0.3 | -1.7 | -13.3 | No |
| 751 | membrane bound O-acyltransferase domain containing 1 | MBOAT1 | -1.7 | -1.3 | -2.2 | -2.7 | -1.5 | No |
| 752 | membrane bound O-acyltransferase domain containing 4 | MBOAT4 | -1.8 | -1.3 | -2.5 | -1.0 | -0.7 | No |
| 753 | multiple coagulation factor deficiency 2 | MCFD2 | 1.3 | 1.4 | 1.4 | 2.2 | 1.5 | No |
| 754 | minichromosome maintenance complex component 3 | MCM3 | 1.3 | 1.5 | 1.2 | 1.3 | 1.1 | No |
| 755 | minichromosome maintenance complex component 4 | MCM4 | 2.6 | 3.1 | 2.2 | 2.2 | 1.8 | No |
| 756 | minichromosome maintenance complex component 5 | MCM5 | 2.8 | 3.2 | 2.5 | 2.6 | 2.0 | No |
| 757 | MCTS1, re-initiation and release factor | MCTS1 | 1.3 | 1.4 | 0.8 | 1.0 | 0.4 | No |
| 758 | midkine | MDK | 3.2 | 3.5 | 4.6 | 4.9 | 4.2 | No |
| 759 | maternal embryonic leucine zipper kinase | MELK | 6.8 | 8.4 | 8.6 | 15.2 | 14.7 | No |
| 760 | mediator of cell motility 1 | MEMO1 | 0.8 | 1.0 | 1.1 | 1.7 | 1.0 | No |
| 761 | meteorin, glial cell differentiation regulator | METRN | -0.1 | -1.1 | -0.6 | -0.1 | 0.2 | No |
| 762 | methyltransferase like 14 | METTL14 | 1.2 | 1.3 | 1.1 | 1.1 | 0.5 | No |
| 763 | methyltransferase like 2 | METTL2 | 1.5 | 1.5 | 1.1 | 1.1 | 0.5 | No |
| 764 | methyltransferase like 3 | METTL3 | 0.7 | 1.0 | 0.8 | 0.8 | 0.4 | No |
| 765 | major facilitator superfamily domain containing 2A | MFSD2A | 0.5 | 1.4 | 1.7 | 2.7 | 1.4 | No |
| 766 | major facilitator superfamily domain containing 6 like | MFSD6L | -0.7 | -0.4 | -0.7 | -0.8 | -0.6 | No |
| 767 | alpha-1,3-mannosyl-glycoprotein 4-beta-N-acetylglucosaminyltransferase B | MGAT4B | -1.9 | -1.7 | -0.9 | -0.6 | -0.8 | No |
| 768 | MICAL like 2 | MICALL2 | -0.4 | -0.5 | -1.0 | -1.0 | -0.9 | No |
| 769 | MID1 interacting protein 1 | MID1IP1 | -1.6 | -0.7 | -1.4 | 0.0 | -1.1 | No |
| 770 | migration and invasion enhancer 1 | MIEN1 | -0.7 | -0.5 | -0.1 | 0.0 | -0.3 | No |
| 771 | major intrinsic protein of lens fiber | MIP | -0.8 | -1.1 | -1.1 | -0.4 | -0.3 | No |
| 772 | MAP kinase interacting serine/threonine kinase 2 | MKNK2 | -0.6 | -0.8 | -0.1 | 0.9 | 1.1 | No |
| 773 | makorin ring finger protein 3 | MKRN3 | -1.3 | -1.0 | -0.6 | -0.4 | -0.7 | No |
| 774 | Meckel syndrome, type 1 | MKS1 | 0.4 | 0.2 | 0.8 | 1.0 | 1.4 | No |
| 775 | MLX, MAX dimerization protein | MLX | -0.1 | -0.7 | -6.8 | -2.6 | -2.1 | No |
| 776 | MNAT1, CDK activating kinase assembly factor | MNAT1 | 0.8 | 1.0 | 0.9 | 1.0 | 0.6 | No |
| 777 | MORN repeat containing 4 | MORN4 | 3.1 | 3.0 | 3.9 | 4.1 | 3.1 | No |
| 778 | Mov10 RISC complex RNA helicase | MOV10 | 1.2 | 0.6 | 1.0 | 1.1 | 0.7 | No |
| 779 | mannose-P-dolichol utilization defect 1 | MPDU1 | -0.5 | -0.8 | -0.6 | -1.5 | -1.6 | No |
| 780 | M-phase phosphoprotein 6 | MPHOSPH6 | 1.0 | 1.0 | 0.7 | 0.6 | 0.1 | No |
| 781 | metallophosphoesterase domain containing 2 | MPPED2 | 1.4 | 1.4 | 2.0 | 3.7 | 2.9 | No |
| 782 | MPV17 mitochondrial inner membrane protein like 2 | MPV17L2 | 2.3 | 2.0 | 1.6 | 0.4 | 0.2 | No |
| 783 | myelin protein zero | MPZ | -0.9 | -1.8 | -2.1 | -2.3 | -2.2 | No |
| 784 | myelin protein zero like 2 | MPZL2 | -0.8 | -0.8 | -0.6 | -1.0 | -0.9 | No |
| 785 | MRG domain binding protein | MRGBP | 0.5 | 0.7 | 0.5 | 0.6 | 0.4 | No |
| 786 | mitochondrial ribosomal protein L1 | MRPL1 | 2.0 | 2.1 | 1.6 | 2.2 | 1.7 | No |
| 787 | mitochondrial ribosomal protein L10 | MRPL10 | 1.4 | 1.8 | 1.1 | 0.9 | 0.6 | No |
| 788 | mitochondrial ribosomal protein L11 | MRPL11 | 1.3 | 1.3 | 0.3 | 0.5 | -0.2 | No |
| 789 | mitochondrial ribosomal protein L12 | MRPL12 | 1.5 | 1.7 | 0.7 | 0.4 | 0.4 | No |
| 790 | mitochondrial ribosomal protein L13 | MRPL13 | 1.6 | 1.5 | 1.0 | 1.0 | 0.7 | No |
| 791 | mitochondrial ribosomal protein L14 | MRPL14 | 1.7 | 1.7 | 1.0 | 1.1 | 0.8 | No |
| 792 | mitochondrial ribosomal protein L19 | MRPL19 | 0.7 | 0.9 | 0.3 | 0.4 | 0.2 | No |
| 793 | mitochondrial ribosomal protein L24 | MRPL24 | 1.3 | 1.4 | 0.7 | 0.8 | 0.5 | No |
| 794 | mitochondrial ribosomal protein L27 | MRPL27 | 1.4 | 1.4 | 1.1 | 0.8 | 0.6 | No |
| 795 | mitochondrial ribosomal protein L28 | MRPL28 | 1.0 | 1.1 | 0.3 | 0.2 | 0.2 | No |
| 796 | mitochondrial ribosomal protein L36 | MRPL36 | 1.0 | 1.1 | 0.7 | 0.7 | -0.1 | No |
| 797 | mitochondrial ribosomal protein L38 | MRPL38 | 1.0 | 0.9 | 0.4 | 0.2 | 0.0 | No |
| 798 | mitochondrial ribosomal protein L40 | MRPL40 | 1.2 | 1.3 | 1.0 | 1.0 | 0.6 | No |
| 799 | mitochondrial ribosomal protein L44 | MRPL44 | 1.0 | 1.1 | 0.5 | 0.5 | 0.4 | No |
| 800 | mitochondrial ribosomal protein L46 | MRPL46 | 1.4 | 1.3 | 0.9 | 0.6 | 0.3 | No |
| 801 | mitochondrial ribosomal protein L9 | MRPL9 | 1.2 | 1.3 | 0.7 | 0.5 | -0.1 | No |
| 802 | mitochondrial ribosomal protein S14 | MRPS14 | 1.0 | 1.1 | 0.3 | 0.9 | 0.6 | No |
| 803 | mitochondrial ribosomal protein S18A | MRPS18A | 0.8 | 1.3 | 0.3 | 0.6 | 0.2 | No |
| 804 | mitochondrial ribosomal protein S18B | MRPS18B | 0.8 | 1.1 | 0.6 | 0.2 | 0.4 | No |
| 805 | mitochondrial ribosomal protein S18C | MRPS18C | 1.0 | 1.3 | 0.5 | 0.4 | -0.1 | No |
| 806 | mitochondrial ribosomal protein S23 | MRPS23 | 1.1 | 1.5 | 0.6 | 1.3 | -0.8 | No |
| 807 | mitochondrial ribosomal protein S26 | MRPS26 | 1.0 | 1.0 | 0.5 | 0.4 | 0.1 | No |
| 808 | mitochondrial ribosomal protein S27 | MRPS27 | 1.1 | 1.0 | 0.4 | 0.8 | 0.1 | No |
| 809 | mitochondrial ribosomal protein S34 | MRPS34 | 1.3 | 1.5 | 0.9 | 0.7 | 0.4 | No |
| 810 | mitochondrial ribosomal protein S6 | MRPS6 | 0.8 | 1.0 | 0.7 | 0.4 | 0.1 | No |
| 811 | MRT4 homolog, ribosome maturation factor | MRTO4 | 2.4 | 2.0 | 1.1 | 0.6 | 0.1 | No |
| 812 | mutS homolog 2 | MSH2 | 1.9 | 2.4 | 1.4 | 1.7 | 1.2 | No |
| 813 | moesin | MSN | 1.2 | 0.8 | 1.2 | 1.9 | 1.5 | No |
| 814 | mitochondrial carrier 2 | MTCH2 | 0.8 | 1.0 | 0.4 | 0.5 | 0.1 | No |
| 815 | myotubularin related protein 10 | MTMR10 | -1.5 | -1.1 | -1.2 | -0.7 | -0.7 | No |
| 816 | myotubularin related protein 12 | MTMR12 | -1.3 | -0.7 | -0.8 | -1.4 | -2.3 | No |
| 817 | microsomal triglyceride transfer protein | MTTP | -1.5 | -1.5 | -2.0 | -2.8 | -0.5 | No |
| 818 | metaxin 1 | MTX1 | 1.3 | 1.9 | 0.6 | 0.6 | 0.7 | No |
| 819 | metaxin 2 | MTX2 | 1.0 | 1.2 | 0.8 | 0.6 | 0.0 | No |
| 820 | mevalonate kinase | MVK | 1.0 | 1.8 | 0.5 | 0.9 | 0.9 | No |
| 821 | myc-like oncogene, s-myc protein | MYCS | 1.5 | 1.0 | 1.8 | 0.1 | 0.4 | No |
| 822 | myosin light chain 6 | MYL6 | 0.4 | 1.0 | 2.0 | 2.4 | 2.4 | No |
| 823 | myosin regulatory light chain interacting protein | MYLIP | -1.3 | -1.5 | -0.6 | -0.4 | -1.3 | No |
| 824 | N(alpha)-acetyltransferase 40, NatD catalytic subunit | NAA40 | 0.8 | 1.5 | 1.0 | -0.1 | -0.3 | No |
| 825 | nascent polypeptide associated complex subunit alpha | NACA | -0.5 | -0.7 | -1.0 | -1.1 | -0.8 | No |
| 826 | NEDD8 activating enzyme E1 subunit 1 | NAE1 | 1.2 | 1.0 | 0.9 | 1.0 | 0.5 | No |
| 827 | nuclear prelamin A recognition factor | NARF | 0.8 | 0.8 | 0.6 | 1.0 | 0.8 | No |
| 828 | nuclear autoantigenic sperm protein | NASP | 1.1 | 1.7 | 1.3 | 0.7 | 0.1 | No |
| 829 | N-acetyltransferase 10 | NAT10 | 1.5 | 1.4 | 0.8 | 0.2 | 0.1 | No |
| 830 | neuron navigator 3 | NAV3 | -1.8 | -3.1 | -1.2 | -1.5 | -1.6 | No |
| 831 | non-SMC condensin II complex subunit D3 | NCAPD3 | 1.1 | 2.6 | 1.8 | 2.3 | 2.0 | No |
| 832 | non-SMC condensin I complex subunit H | NCAPH | 0.7 | 2.6 | 1.9 | 2.7 | 2.4 | No |
| 833 | nuclear cap binding protein subunit 2 | NCBP2 | 0.9 | 1.1 | 0.9 | 1.0 | 0.7 | No |
| 834 | non-specific cytotoxic cell receptor protein 1 homolog (zebrafish) | NCCRP1 | 2.2 | 1.7 | 0.5 | 0.2 | 0.4 | No |
| 835 | NCK adaptor protein 2 | NCK2 | -5.9 | -0.4 | -5.7 | -11.7 | -1.0 | No |
| 836 | nuclear receptor coactivator 5 | NCOA5 | -1.0 | -0.2 | -0.7 | -0.4 | -0.6 | No |
| 837 | nuclear receptor corepressor 1 | NCOR1 | -1.1 | -1.3 | -1.0 | -0.9 | -1.0 | No |
| 838 | NDC80, kinetochore complex component | NDC80 | 1.5 | 2.9 | 2.4 | 3.0 | 2.6 | No |
| 839 | NADH:ubiquinone oxidoreductase subunit AB1 | NDUFAB1 | 1.5 | 1.6 | 1.0 | 1.0 | 0.7 | No |
| 840 | NADH:ubiquinone oxidoreductase subunit B10 | NDUFB10 | 0.6 | 1.0 | 0.2 | 0.3 | 0.2 | No |
| 841 | NADH:ubiquinone oxidoreductase subunit B11 | NDUFB11 | 0.9 | 1.1 | 0.8 | 0.6 | 0.3 | No |
| 842 | NADH:ubiquinone oxidoreductase subunit S4 | NDUFS4 | 1.3 | 1.2 | 0.9 | 1.0 | 0.8 | No |
| 843 | NADH:ubiquinone oxidoreductase subunit S5 | NDUFS5 | 0.7 | 1.0 | 0.6 | 0.8 | 0.5 | No |
| 844 | NADH:ubiquinone oxidoreductase subunit S6 | NDUFS6 | 0.8 | 1.1 | 0.5 | 0.6 | 0.3 | No |
| 845 | neural precursor cell expressed, developmentally down-regulated 8 | NEDD8 | 0.7 | 1.2 | 0.8 | 0.9 | 0.6 | No |
| 846 | nei like DNA glycosylase 3 | NEIL3 | 0.3 | 2.3 | 1.1 | 2.2 | 1.2 | No |
| 847 | NIMA related kinase 2 | NEK2 | 1.2 | 2.1 | 2.8 | 3.4 | 3.1 | No |
| 848 | negative elongation factor complex member A | NELFA | -0.8 | -0.4 | -0.4 | -1.0 | -0.5 | No |
| 849 | nuclear factor, erythroid 2 like 2 | NFE2L2 | -1.3 | -1.1 | -1.2 | -0.7 | -0.3 | No |
| 850 | nuclear factor I A | NFIA | -1.1 | -1.3 | -1.4 | -1.4 | -2.0 | No |
| 851 | NHP2 ribonucleoprotein | NHP2 | 1.7 | 1.3 | 0.9 | 0.9 | 0.4 | No |
| 852 | NGG1 interacting factor 3 like 1 | NIF3L1 | 1.0 | 1.0 | 0.6 | 0.4 | 0.1 | No |
| 853 | NIP7, nucleolar pre-rRNA processing protein | NIP7 | 2.2 | 1.8 | 1.1 | 0.8 | 0.3 | No |
| 854 | NIPA magnesium transporter 1 | NIPA1 | 0.8 | 1.0 | 0.6 | 0.6 | 0.3 | No |
| 855 | sodium/potassium transporting ATPase interacting 1 | NKAIN1 | -0.4 | 1.0 | 1.9 | 3.7 | 2.8 | No |
| 856 | naked cuticle homolog 2 | NKD2 | 1.2 | 2.4 | 1.9 | 0.4 | 0.8 | No |
| 857 | N-myristoyltransferase 1 | NMT1 | 1.3 | 1.5 | 1.2 | 1.2 | 0.8 | No |
| 858 | nicotinamide nucleotide transhydrogenase | NNT | 1.3 | 1.4 | 1.1 | 1.3 | 0.8 | No |
| 859 | NOC2 like nucleolar associated transcriptional repressor | NOC2L | 1.9 | 1.2 | 0.6 | 0.6 | 0.0 | No |
| 860 | NOC3 like DNA replication regulator | NOC3L | 1.0 | 1.0 | 0.3 | 0.1 | 0.0 | No |
| 861 | noggin | NOG | -1.3 | -2.2 | -1.4 | -1.3 | -0.6 | No |
| 862 | nucleolar protein with MIF4G domain 1 | NOM1 | 1.9 | 1.7 | 1.1 | 0.9 | 0.4 | No |
| 863 | NADPH oxidase activator 1 | NOXA1 | -0.6 | -0.3 | -1.0 | -1.0 | -0.3 | No |
| 864 | nucleophosmin 1 | NPM1 | 2.4 | 2.2 | 1.2 | 1.0 | 0.5 | No |
| 865 | neuronal pentraxin 1 | NPTX1 | 2.1 | 3.7 | 3.2 | 2.3 | 1.4 | No |
| 866 | NAD(P)H quinone dehydrogenase 1 | NQO1 | -0.5 | -0.7 | -1.0 | -0.6 | -1.8 | No |
| 867 | nuclear receptor subfamily 2 group F member 6 | NR2F6 | 0.7 | 0.7 | 0.6 | 1.2 | 0.7 | No |
| 868 | nuclear receptor subfamily 4 group A member 1 | NR4A1 | -0.1 | -1.0 | -1.3 | -1.3 | -1.4 | No |
| 869 | neuregulin 2 | NRG2 | -0.7 | -1.0 | -1.0 | -1.2 | -0.5 | No |
| 870 | neuritin 1 | NRN1 | 1.7 | 2.9 | 1.5 | 2.1 | -1.4 | No |
| 871 | neurensin 1 | NRSN1 | 8.7 | 9.8 | 9.3 | 18.1 | 17.8 | No |
| 872 | neurexin 3 | NRXN3 | -1.4 | -1.9 | -0.9 | -1.0 | -0.3 | No |
| 873 | NSA2, ribosome biogenesis homolog | NSA2 | -1.2 | -1.8 | -1.6 | -1.8 | -1.2 | No |
| 874 | nuclear receptor binding SET domain protein 2 | NSD2 | 1.1 | 1.9 | 1.1 | 0.1 | 0.9 | No |
| 875 | NSL1, MIS12 kinetochore complex component | NSL1 | 1.4 | 2.4 | 2.0 | 2.3 | 1.8 | No |
| 876 | nucleobindin 2 | NUCB2 | 0.9 | 1.0 | 1.3 | 1.6 | 1.6 | No |
| 877 | nuclear casein kinase and cyclin dependent kinase substrate 1 | NUCKS1 | 0.6 | 0.9 | 1.1 | 1.2 | 0.8 | No |
| 878 | NudC domain containing 1 | NUDCD1 | 1.2 | 1.0 | 1.1 | 1.2 | 0.6 | No |
| 879 | nudix hydrolase 14 | NUDT14 | 1.1 | 0.5 | 0.8 | 1.3 | 1.0 | No |
| 880 | nucleoporin 107 | NUP107 | 1.0 | 1.2 | 0.7 | 0.8 | 0.4 | No |
| 881 | nucleoporin 133 | NUP133 | 1.2 | 1.4 | 0.7 | 0.0 | 0.0 | No |
| 882 | nucleoporin 43 | NUP43 | 1.5 | 1.2 | 0.9 | -0.5 | -0.3 | No |
| 883 | nucleoporin 88 | NUP88 | 0.9 | 1.0 | 0.5 | 0.5 | 0.3 | No |
| 884 | nucleoporin 93 | NUP93 | 1.3 | 1.2 | 0.8 | 0.7 | 0.1 | No |
| 885 | nucleolar and spindle associated protein 1 | NUSAP1 | 1.0 | 2.2 | 2.7 | 2.4 | 2.4 | No |
| 886 | ornithine decarboxylase antizyme 2 | OAZ2 | -0.7 | -1.4 | -0.7 | 0.1 | -0.5 | No |
| 887 | occludin | OCLN | -1.5 | -1.6 | -1.0 | -1.4 | -2.3 | No |
| 888 | oncomodulin | OCM | -1.0 | -1.1 | -1.6 | -1.8 | -2.5 | No |
| 889 | olfactomedin like 3 | OLFML3 | 0.7 | 0.8 | 2.1 | 3.5 | 3.3 | No |
| 890 | OPA3, outer mitochondrial membrane lipid metabolism regulator | OPA3 | 0.6 | 0.7 | 0.6 | 0.2 | 0.2 | No |
| 891 | opioid receptor delta 1 | OPRD1 | -1.5 | -2.0 | -1.0 | -1.0 | -2.3 | No |
| 892 | origin recognition complex subunit 5 | ORC5 | 1.6 | 1.8 | 0.9 | 0.8 | 0.8 | No |
| 893 | oxysterol binding protein like 7 | OSBPL7 | -0.9 | -1.2 | -1.4 | -1.9 | -1.1 | No |
| 894 | purinergic receptor P2X 3 | P2RX3 | -0.6 | -1.3 | -2.1 | -2.0 | -0.9 | No |
| 895 | purinergic receptor P2X 7 | P2RX7 | -1.4 | -1.4 | -0.5 | -0.5 | -1.0 | No |
| 896 | proliferation-associated 2G4 | PA2G4 | 1.5 | 1.4 | 0.8 | 0.8 | 0.4 | No |
| 897 | phenylalanine hydroxylase | PAH | -2.0 | -2.2 | -1.9 | -2.0 | -1.1 | No |
| 898 | poly(A) binding protein interacting protein 1 | PAIP1 | -0.8 | -0.5 | -0.3 | -1.1 | -1.0 | No |
| 899 | PAK1 interacting protein 1 | PAK1IP1 | 1.6 | 1.3 | 0.5 | 0.1 | -0.1 | No |
| 900 | 3'-phosphoadenosine 5'-phosphosulfate synthase 1 | PAPSS1 | 2.4 | 1.2 | 2.1 | 0.6 | 0.6 | No |
| 901 | progestin and adipoQ receptor family member 5 | PAQR5 | -1.4 | -2.2 | -1.6 | -2.1 | -1.1 | No |
| 902 | par-6 family cell polarity regulator alpha | PARD6A | -0.5 | -1.0 | -0.8 | -0.8 | 0.0 | No |
| 903 | presenilin associated rhomboid like | PARL | -1.1 | -0.7 | -0.8 | 0.0 | -0.2 | No |
| 904 | poly(ADP-ribose) polymerase family member 3 | PARP3 | 0.7 | 1.1 | 1.2 | 1.0 | 0.8 | No |
| 905 | PARP1 binding protein | PARPBP | 0.5 | 1.0 | 2.5 | 3.2 | 2.1 | No |
| 906 | parvin beta | PARVB | -1.7 | -1.2 | -1.2 | -3.2 | -0.5 | No |
| 907 | paired box 1 | PAX1 | 1.2 | 0.7 | 9.5 | 17.2 | 18.0 | No |
| 908 | PDZ binding kinase | PBK | 2.1 | 3.2 | 3.3 | 5.2 | 4.4 | No |
| 909 | polycomb group ring finger 5 | PCGF5 | -0.7 | -0.6 | -1.3 | -1.0 | -1.2 | No |
| 910 | polycomb group ring finger 6 | PCGF6 | 1.6 | 1.5 | 1.2 | 0.9 | 0.2 | No |
| 911 | phosphoenolpyruvate carboxykinase 2, mitochondrial | PCK2 | 0.8 | 1.0 | 1.0 | 1.1 | 0.9 | No |
| 912 | protein-L-isoaspartate (D-aspartate) O-methyltransferase domain containing 2 | PCMTD2 | -1.4 | -2.2 | -0.9 | -0.2 | -0.4 | No |
| 913 | proliferating cell nuclear antigen pseudogene 2 | PCNA-PS2 | 1.3 | 2.0 | 1.3 | 1.5 | 1.1 | No |
| 914 | PEST proteolytic signal containing nuclear protein | PCNP | -0.7 | -0.4 | -0.8 | -0.9 | -1.0 | No |
| 915 | programmed cell death 2 | PDCD2 | 1.2 | 1.2 | 0.6 | 0.5 | -0.1 | No |
| 916 | programmed cell death 4 | PDCD4 | -2.2 | -1.8 | -1.5 | -2.3 | -2.0 | No |
| 917 | phosphodiesterase 12 | PDE12 | 0.7 | 0.4 | 0.1 | 0.1 | 0.3 | No |
| 918 | pyruvate dehydrogenase E1 alpha 2 subunit | PDHA2 | -1.5 | -8.0 | -1.4 | -2.0 | -13.4 | No |
| 919 | protein disulfide isomerase family A member 4 | PDIA4 | 1.1 | 1.2 | 1.3 | 1.8 | 1.7 | No |
| 920 | PDZ and LIM domain 4 | PDLIM4 | 3.9 | 3.0 | 2.6 | 1.1 | 1.2 | No |
| 921 | 3-phosphoinositide dependent protein kinase 1 | PDPK1 | -1.9 | -1.7 | -1.0 | -0.6 | -0.8 | No |
| 922 | decaprenyl diphosphate synthase subunit 1 | PDSS1 | 2.0 | 2.4 | 1.1 | 0.8 | 0.7 | No |
| 923 | decaprenyl diphosphate synthase subunit 2 | PDSS2 | 1.8 | 1.7 | 1.2 | 0.5 | 0.5 | No |
| 924 | pancreatic and duodenal homeobox 1 | PDX1 | 0.9 | 0.2 | 1.3 | 1.4 | -0.1 | No |
| 925 | pellino E3 ubiquitin protein ligase family member 2 | PELI2 | -1.0 | -1.0 | -1.2 | -0.8 | -1.4 | No |
| 926 | pelota mRNA surveillance and ribosome rescue factor | PELO | 1.7 | 1.5 | 1.1 | 0.7 | 0.9 | No |
| 927 | proenkephalin | PENK | -1.2 | -1.8 | -1.4 | -1.1 | -2.5 | No |
| 928 | pescadillo ribosomal biogenesis factor 1 | PES1 | 1.8 | 1.3 | 0.7 | 0.3 | 0.0 | No |
| 929 | prefoldin subunit 2 | PFDN2 | 0.9 | 0.9 | 0.6 | 0.6 | 0.5 | No |
| 930 | prefoldin subunit 6 | PFDN6 | 1.0 | 1.2 | 0.9 | 1.0 | 0.4 | No |
| 931 | phosphoglycerate mutase 1 | PGAM1 | 1.3 | 1.5 | 0.7 | 1.3 | 1.1 | No |
| 932 | phosphogluconate dehydrogenase | PGD | 1.0 | 1.2 | 0.7 | 0.7 | 0.3 | No |
| 933 | phosphoglucomutase 1 | PGM1 | 1.6 | 1.7 | 1.0 | 0.9 | 0.7 | No |
| 934 | phosphoglucomutase 3 | PGM3 | 1.6 | 1.9 | 1.6 | 1.7 | 1.1 | No |
| 935 | phosphoglycolate phosphatase | PGP | 1.9 | 1.7 | 1.6 | 1.5 | 1.5 | No |
| 936 | phosphorylated adaptor for RNA export | PHAX | 0.9 | 1.1 | 0.7 | 0.7 | 0.4 | No |
| 937 | prohibitin | PHB | 2.0 | 1.9 | 1.1 | 1.1 | 0.7 | No |
| 938 | polyhomeotic homolog 2 | PHC2 | 0.8 | 1.3 | 1.0 | 0.9 | 0.6 | No |
| 939 | phosphate regulating endopeptidase homolog X-linked | PHEX | 1.4 | 1.2 | 3.3 | 4.7 | 3.3 | No |
| 940 | PHD finger protein 5A | PHF5A | 1.4 | 1.4 | 1.2 | 1.1 | 1.0 | No |
| 941 | phosphorylase kinase catalytic subunit gamma 1 | PHKG1 | -0.3 | -0.3 | -1.5 | -0.6 | -0.4 | No |
| 942 | pleckstrin homology like domain family A member 3 | PHLDA3 | 0.8 | 1.1 | 0.6 | 1.1 | 1.4 | No |
| 943 | phosphatidylinositol glycan anchor biosynthesis class F | PIGF | 0.7 | 1.0 | 0.6 | 1.1 | 0.6 | No |
| 944 | phosphatidylinositol glycan anchor biosynthesis class H | PIGH | -1.0 | -1.6 | -1.3 | -0.9 | -0.9 | No |
| 945 | phosphatidylinositol-4,5-bisphosphate 3-kinase catalytic subunit delta | PIK3CD | -0.6 | -1.1 | -0.9 | -1.3 | -1.8 | No |
| 946 | phosphoinositide-3-kinase regulatory subunit 2 | PIK3R2 | -2.1 | -0.8 | -1.1 | 0.0 | -0.8 | No |
| 947 | peptidylprolyl cis/trans isomerase, NIMA-interacting 4 | PIN4 | 1.0 | 1.3 | 1.2 | 1.1 | 0.8 | No |
| 948 | phosphatidylinositol-4-phosphate 5-kinase type 1 beta | PIP5K1B | -0.2 | -0.4 | -0.5 | -1.3 | -1.4 | No |
| 949 | phosphatidylinositol transfer protein beta | PITPNB | 7.0 | 7.0 | 6.3 | -1.3 | -2.3 | No |
| 950 | pitrilysin metallopeptidase 1 | PITRM1 | 1.3 | 1.4 | 1.1 | 0.8 | 0.5 | No |
| 951 | plasminogen activator, urokinase | PLAU | -0.4 | -1.5 | -1.1 | -1.9 | -1.0 | No |
| 952 | pleckstrin 2 | PLEK2 | 0.0 | -1.1 | -0.8 | -1.0 | 0.3 | No |
| 953 | pleckstrin homology domain containing A3 | PLEKHA3 | -0.8 | -0.5 | -0.5 | -0.3 | -0.5 | No |
| 954 | pleckstrin homology domain containing B2 | PLEKHB2 | -1.4 | -1.3 | -0.9 | -0.4 | -1.0 | No |
| 955 | pleckstrin homology, MyTH4 and FERM domain containing H1 | PLEKHH1 | 0.9 | 1.3 | 1.4 | 1.4 | 1.0 | No |
| 956 | perilipin 2 | PLIN2 | 1.9 | 0.9 | 1.4 | 2.0 | 0.2 | No |
| 957 | polo like kinase 1 | PLK1 | 0.2 | 1.6 | 2.2 | 3.2 | 2.9 | No |
| 958 | polo like kinase 4 | PLK4 | 1.1 | 2.4 | 1.8 | 2.0 | 2.0 | No |
| 959 | procollagen-lysine,2-oxoglutarate 5-dioxygenase 1 | PLOD1 | 0.8 | 0.8 | 2.3 | 4.5 | 4.3 | No |
| 960 | procollagen-lysine,2-oxoglutarate 5-dioxygenase 2 | PLOD2 | 1.2 | -4.9 | 4.1 | 4.5 | 3.5 | No |
| 961 | pleiotropic regulator 1 | PLRG1 | 1.0 | 0.9 | 0.7 | 0.5 | 0.3 | No |
| 962 | phosphomannomutase 2 | PMM2 | 1.8 | 2.0 | 1.2 | 1.2 | 1.1 | No |
| 963 | partner of NOB1 homolog | PNO1 | -0.5 | -0.9 | -0.9 | -0.9 | -1.1 | No |
| 964 | purine nucleoside phosphorylase | PNP | 1.7 | 0.5 | 1.2 | 0.5 | 1.9 | No |
| 965 | POC1 centriolar protein B | POC1B | 0.6 | 1.1 | 0.8 | 1.1 | 0.8 | No |
| 966 | DNA polymerase alpha 2, accessory subunit | POLA2 | 1.7 | 2.6 | 1.7 | 2.2 | 1.5 | No |
| 967 | DNA polymerase beta | POLB | 1.1 | 1.1 | 1.0 | 0.8 | 0.7 | No |
| 968 | DNA polymerase delta 1, catalytic subunit | POLD1 | 1.3 | 1.8 | 1.0 | 0.8 | 0.8 | No |
| 969 | DNA polymerase delta 3, accessory subunit | POLD3 | 1.6 | 2.2 | 1.4 | 1.3 | 0.8 | No |
| 970 | DNA polymerase epsilon, catalytic subunit | POLE | 1.5 | 2.5 | 1.6 | 2.0 | 1.7 | No |
| 971 | DNA polymerase epsilon 3, accessory subunit | POLE3 | 0.8 | 1.3 | 0.8 | 1.0 | 0.5 | No |
| 972 | DNA polymerase eta | POLH | 0.7 | 1.5 | 0.5 | 0.7 | 0.6 | No |
| 973 | RNA polymerase I and III subunit D | POLR1D | 1.6 | 1.4 | 0.8 | 0.4 | 0.1 | No |
| 974 | RNA polymerase II subunit C | POLR2C | 0.9 | 1.0 | 0.8 | 0.6 | 0.3 | No |
| 975 | RNA polymerase II subunit E | POLR2E | 1.5 | 1.0 | 0.6 | 0.4 | 0.1 | No |
| 976 | RNA polymerase II subunit G | POLR2G | 0.7 | 1.0 | 0.4 | 0.8 | 0.5 | No |
| 977 | RNA polymerase II subunit I | POLR2I | 1.0 | 1.0 | 0.9 | 1.0 | 0.8 | No |
| 978 | proteasome maturation protein | POMP | 1.6 | 1.5 | 1.4 | 1.2 | 0.8 | No |
| 979 | POP7 homolog, ribonuclease P/MRP subunit | POP7 | 0.7 | 0.7 | 0.7 | 1.0 | 1.0 | No |
| 980 | POU class 3 homeobox 1 | POU3F1 | -0.1 | -1.7 | -1.3 | -2.9 | -1.1 | No |
| 981 | POU class 4 homeobox 1 | POU4F1 | -1.7 | -1.7 | -1.5 | -1.5 | -2.6 | No |
| 982 | pyrophosphatase (inorganic) 1 | PPA1 | 0.9 | 0.7 | 0.3 | 0.6 | 0.4 | No |
| 983 | peter pan homolog | PPAN | 1.8 | 1.4 | 0.9 | 0.6 | 0.4 | No |
| 984 | phosphoribosyl pyrophosphate amidotransferase | PPAT | 0.9 | 1.0 | 0.9 | 0.6 | 0.3 | No |
| 985 | pancreatic progenitor cell differentiation and proliferation factor | PPDPF | 1.7 | 1.7 | 1.4 | 1.6 | 0.7 | No |
| 986 | PTPRF interacting protein alpha 2 | PPFIA2 | -1.4 | -0.8 | -0.4 | -0.2 | -1.2 | No |
| 987 | PPFIA binding protein 2 | PPFIBP2 | -0.7 | -0.2 | -0.4 | -0.7 | -1.9 | No |
| 988 | peptidylprolyl isomerase B | PPIB | 0.3 | 0.5 | 0.9 | 1.6 | 1.5 | No |
| 989 | peptidylprolyl isomerase D | PPID | 0.8 | 1.3 | 0.7 | 1.3 | 0.6 | No |
| 990 | peptidylprolyl isomerase G | PPIG | 1.0 | 1.3 | 0.7 | 0.0 | 0.0 | No |
| 991 | protein phosphatase, Mg2+/Mn2+ dependent 1E | PPM1E | -0.1 | 0.4 | 2.2 | 2.1 | 2.2 | No |
| 992 | protein phosphatase, Mg2+/Mn2+ dependent 1K | PPM1K | -1.2 | -0.8 | -0.7 | -1.1 | -0.8 | No |
| 993 | protein phosphatase 1 regulatory subunit 13B | PPP1R13B | -1.2 | -1.1 | -0.7 | -1.4 | -1.0 | No |
| 994 | protein phosphatase 1 regulatory subunit 3C | PPP1R3C | -1.6 | -1.8 | -1.8 | -2.2 | -2.2 | No |
| 995 | protein phosphatase 2 scaffold subunit Aalpha | PPP2R1A | 0.8 | 1.1 | 0.8 | 0.8 | 0.6 | No |
| 996 | protein phosphatase 3 regulatory subunit B, alpha | PPP3R1 | 0.8 | 1.0 | 0.6 | 2.4 | -0.4 | No |
| 997 | protein phosphatase 4 regulatory subunit 2 | PPP4R2 | 0.5 | 0.6 | 0.4 | 0.4 | 0.4 | No |
| 998 | PRA1 domain family member 2 | PRAF2 | 0.6 | 0.5 | 1.0 | 0.6 | 1.1 | No |
| 999 | PR/SET domain 1 | PRDM1 | 1.5 | 1.3 | 2.6 | 3.3 | 3.1 | No |
| 1000 | peroxiredoxin 6B | PRDX6B | 1.1 | 1.1 | 0.8 | 0.8 | 0.4 | No |
| 1001 | perforin 1 | PRF1 | -2.1 | -2.6 | -1.6 | -1.6 | -2.3 | No |
| 1002 | proteoglycan 4 | PRG4 | -0.3 | -1.0 | -1.6 | -1.9 | -2.6 | No |
| 1003 | prickle planar cell polarity protein 1 | PRICKLE1 | 1.8 | 2.6 | 1.9 | 2.6 | 2.2 | No |
| 1004 | protein kinase C beta | PRKCB | -0.8 | -1.0 | -0.5 | -1.2 | -0.9 | No |
| 1005 | protein kinase C eta | PRKCH | -0.6 | -1.5 | -0.9 | -1.3 | -1.6 | No |
| 1006 | protein kinase C substrate 80K-H | PRKCSH | 1.1 | 1.3 | 1.2 | 1.4 | 1.1 | No |
| 1007 | protein activator of interferon induced protein kinase EIF2AK2 | PRKRA | 1.7 | 1.5 | 1.2 | 0.6 | 0.3 | No |
| 1008 | protein arginine methyltransferase 1 | PRMT1 | 2.0 | 1.9 | 1.2 | 1.0 | 0.3 | No |
| 1009 | protein arginine methyltransferase 5 | PRMT5 | 1.8 | 2.0 | 1.2 | 1.1 | 0.7 | No |
| 1010 | protein arginine methyltransferase 7 | PRMT7 | 1.7 | 1.5 | 0.9 | 0.4 | 0.4 | No |
| 1011 | pre-mRNA processing factor 18 | PRPF18 | 0.4 | 1.5 | 0.3 | 0.1 | 0.2 | No |
| 1012 | pre-mRNA processing factor 40 homolog A | PRPF40A | 1.0 | 0.8 | 0.6 | 0.5 | 0.4 | No |
| 1013 | phosphoribosyl pyrophosphate synthetase 1-like 1 | PRPS1L1 | 0.9 | 1.4 | 1.1 | 1.2 | 0.9 | No |
| 1014 | proline rich coiled-coil 1 | PRRC1 | 0.8 | 1.0 | 0.8 | 1.4 | 1.1 | No |
| 1015 | serine protease 38 | PRSS38 | -2.1 | -3.3 | -1.5 | -2.8 | -3.1 | No |
| 1016 | serine protease 42 | PRSS42 | 0.2 | 0.7 | 0.9 | 1.4 | 1.4 | No |
| 1017 | serine protease 8 | PRSS8 | -0.5 | -0.7 | -0.7 | -0.7 | -0.5 | No |
| 1018 | pregnancy specific glycoprotein 18 | PSG18 | -0.5 | -1.1 | -2.9 | -3.2 | -3.6 | No |
| 1019 | proteasome subunit alpha 2 | PSMA2 | 1.1 | 1.2 | 1.1 | 0.8 | 0.6 | No |
| 1020 | proteasome subunit alpha 4 | PSMA4 | 1.2 | 1.2 | 1.1 | 0.9 | 0.5 | No |
| 1021 | proteasome subunit alpha 5 | PSMA5 | 1.6 | 1.5 | 1.4 | 1.3 | 0.8 | No |
| 1022 | proteasome subunit alpha 6 | PSMA6 | 1.0 | 1.0 | 0.8 | 0.1 | 0.1 | No |
| 1023 | proteasome subunit beta 1 | PSMB1 | 1.7 | 1.6 | 1.5 | 1.2 | 0.9 | No |
| 1024 | proteasome subunit beta 2 | PSMB2 | 1.1 | 1.0 | 1.3 | 0.9 | 0.5 | No |
| 1025 | proteasome subunit beta 3 | PSMB3 | 1.7 | 1.5 | 1.6 | 1.2 | 0.8 | No |
| 1026 | proteasome subunit beta 6 | PSMB6 | 1.1 | 1.3 | 1.0 | 0.9 | 0.6 | No |
| 1027 | proteasome 26S subunit, ATPase 1 | PSMC1 | 1.7 | 1.4 | 1.5 | 1.2 | 0.9 | No |
| 1028 | proteasome 26S subunit, ATPase 2 | PSMC2 | 1.4 | 1.1 | 1.1 | 0.8 | 0.5 | No |
| 1029 | proteasome 26S subunit, ATPase 3 | PSMC3 | 1.7 | 1.6 | 1.4 | 1.2 | 0.7 | No |
| 1030 | PSMC3 interacting protein | PSMC3IP | 0.9 | 2.1 | 1.3 | 2.2 | 1.4 | No |
| 1031 | proteasome 26S subunit, ATPase 4 | PSMC4 | 1.7 | 1.4 | 1.2 | 0.8 | 0.7 | No |
| 1032 | proteasome 26S subunit, ATPase 6 | PSMC6 | 1.6 | 1.4 | 1.3 | 1.2 | 0.8 | No |
| 1033 | proteasome 26S subunit, non-ATPase 10 | PSMD10 | 2.0 | 2.0 | 1.6 | 1.3 | 1.2 | No |
| 1034 | proteasome 26S subunit, non-ATPase 11 | PSMD11 | 1.6 | 1.5 | 1.2 | 1.1 | 0.8 | No |
| 1035 | proteasome 26S subunit, non-ATPase 14 | PSMD14 | 1.6 | 1.3 | 1.4 | 1.1 | 0.9 | No |
| 1036 | proteasome 26S subunit, non-ATPase 4 | PSMD4 | 1.3 | 1.2 | 1.1 | 1.0 | 0.6 | No |
| 1037 | proteasome 26S subunit, non-ATPase 5 | PSMD5 | 1.6 | 1.3 | 1.4 | 1.2 | 0.7 | No |
| 1038 | proteasome 26S subunit, non-ATPase 6 | PSMD6 | 1.5 | 1.2 | 1.2 | 0.8 | 0.6 | No |
| 1039 | proteasome 26S subunit, non-ATPase 8 | PSMD8 | 1.7 | 1.6 | 1.5 | 1.3 | 1.0 | No |
| 1040 | proteasome activator subunit 3 | PSME3 | 1.9 | 1.8 | 1.2 | 0.9 | 0.5 | No |
| 1041 | proteasome assembly chaperone 1 | PSMG1 | 0.9 | 1.3 | 0.7 | 0.7 | 0.6 | No |
| 1042 | patched 2 | PTCH2 | 1.6 | 2.1 | 2.9 | 5.9 | 6.1 | No |
| 1043 | prostaglandin E synthase | PTGES | 3.4 | 3.0 | 2.3 | -0.1 | 0.6 | No |
| 1044 | protein tyrosine phosphatase type IVA, member 1 | PTP4A1 | 0.7 | 0.8 | 0.6 | 1.0 | 0.4 | No |
| 1045 | protein tyrosine phosphatase, receptor type J | PTPRJ | -0.7 | -0.9 | -0.7 | -0.8 | -1.3 | No |
| 1046 | PWP1 homolog, endonuclein | PWP1 | 1.3 | 1.3 | 0.9 | 0.8 | 0.3 | No |
| 1047 | glycogen phosphorylase, muscle associated | PYGM | -0.9 | -0.9 | -2.0 | -1.5 | -0.4 | No |
| 1048 | glutamine rich 1 | QRICH1 | 0.9 | 1.2 | 0.9 | 0.7 | 0.2 | No |
| 1049 | queuine tRNA-ribosyltransferase catalytic subunit 1 | QTRT1 | 0.8 | 1.1 | 0.5 | 0.1 | 0.0 | No |
| 1050 | RAB10, member RAS oncogene family | RAB10 | 1.0 | 0.8 | 0.7 | 0.5 | 0.2 | No |
| 1051 | RAB11A, member RAS oncogene family | RAB11A | 0.7 | 0.7 | 0.5 | 0.3 | 0.1 | No |
| 1052 | RAB13, member RAS oncogene family | RAB13 | 0.5 | 0.9 | 1.1 | 1.5 | 1.1 | No |
| 1053 | RAB34, member RAS oncogene family | RAB34 | 1.1 | 0.9 | 2.3 | 2.5 | 2.2 | No |
| 1054 | RAB3D, member RAS oncogene family | RAB3D | 2.0 | 1.4 | 1.8 | 1.0 | 0.9 | No |
| 1055 | RAB40B, member RAS oncogene family | RAB40B | -1.2 | -1.6 | -0.9 | -0.9 | -0.8 | No |
| 1056 | ras-related protein Rab-7a | RAB7 | -2.8 | -0.5 | -6.3 | -0.9 | -12.3 | No |
| 1057 | Rab geranylgeranyltransferase subunit beta | RABGGTB | 0.9 | 1.2 | 0.9 | 0.9 | 0.8 | No |
| 1058 | RAB, member RAS oncogene family-like 2 | RABL2 | 0.8 | 1.4 | 1.2 | 1.5 | 0.6 | No |
| 1059 | Rac GTPase activating protein 1 | RACGAP1 | 1.5 | 2.4 | 2.8 | 3.0 | 2.7 | No |
| 1060 | RAD51 recombinase | RAD51 | 1.3 | 2.0 | 1.4 | 1.3 | 0.4 | No |
| 1061 | RAD51 associated protein 1 | RAD51AP1 | 0.8 | 2.0 | 1.8 | 1.1 | 1.3 | No |
| 1062 | RAD51 paralog B | RAD51B | 0.4 | 2.0 | 0.9 | 1.6 | 1.1 | No |
| 1063 | RAD52 homolog, DNA repair protein | RAD52 | 0.4 | 2.9 | 1.5 | 1.8 | 1.8 | No |
| 1064 | RAD54 like | RAD54L | 0.8 | 2.5 | 1.6 | 2.6 | 2.1 | No |
| 1065 | ribonucleic acid export 1 | RAE1 | 0.8 | 1.0 | 0.6 | 0.7 | 0.4 | No |
| 1066 | retinoic acid induced 14 | RAI14 | 3.0 | 2.9 | 2.1 | 2.0 | 1.8 | No |
| 1067 | RAN binding protein 1 | RANBP1 | 1.3 | 0.9 | 0.7 | 0.6 | 0.0 | No |
| 1068 | RAN binding protein 6 | RANBP6 | 0.8 | 1.2 | 0.3 | 0.5 | 0.4 | No |
| 1069 | retinoic acid receptor alpha | RARA | -0.5 | -0.9 | -1.5 | -1.5 | -1.3 | No |
| 1070 | RAS p21 protein activator 4 | RASA4 | 8.5 | 8.4 | 8.4 | 16.0 | 16.0 | No |
| 1071 | ras related dexamethasone induced 1 | RASD1 | -1.3 | -1.8 | -2.0 | -2.2 | -2.5 | No |
| 1072 | RB transcriptional corepressor 1 | RB1 | 1.4 | 2.0 | 1.1 | 0.9 | 0.6 | No |
| 1073 | RB binding protein 7, chromatin remodeling factor | RBBP7 | 1.3 | 1.5 | 1.1 | 1.0 | 0.6 | No |
| 1074 | RB binding protein 8, endonuclease | RBBP8 | 1.3 | 2.8 | 1.3 | 1.6 | 1.6 | No |
| 1075 | RB transcriptional corepressor like 1 | RBL1 | 1.4 | 1.3 | 1.4 | 0.9 | 0.9 | No |
| 1076 | RNA binding motif protein 19 | RBM19 | 1.2 | 1.0 | 0.9 | 0.3 | -0.1 | No |
| 1077 | RNA binding motif protein 48 | RBM48 | -1.3 | -1.5 | -0.7 | -0.9 | -0.6 | No |
| 1078 | RNA binding motif protein 8A2 | RBM8A2 | 1.1 | 1.0 | 0.7 | 1.0 | 0.6 | No |
| 1079 | RNA binding motif protein X-linked 2 | RBMX2 | 0.7 | 1.1 | 0.3 | 0.4 | 0.4 | No |
| 1080 | RNA binding protein, mRNA processing factor 2 | RBPMS2 | -1.6 | -2.0 | -1.7 | -1.0 | -0.8 | No |
| 1081 | ring-box 1 | RBX1 | 1.0 | 1.2 | 0.7 | 0.8 | 0.6 | No |
| 1082 | RCAN family member 3 | RCAN3 | -0.9 | -1.1 | -1.1 | -1.2 | -0.4 | No |
| 1083 | regulator of chromosome condensation 1 | RCC1 | 2.3 | 2.1 | 1.4 | 0.7 | 0.7 | No |
| 1084 | reticulocalbin 3 | RCN3 | 0.8 | 1.7 | 3.2 | 5.4 | 5.0 | No |
| 1085 | REST corepressor 2 | RCOR2 | 8.7 | 8.7 | 8.4 | 16.6 | 16.9 | No |
| 1086 | retinol dehydrogenase 14 | RDH14 | -0.7 | -0.9 | -0.8 | -0.6 | -0.2 | No |
| 1087 | replication factor C subunit 1 | RFC1 | 0.9 | 1.6 | 1.0 | 1.4 | 0.8 | No |
| 1088 | replication factor C subunit 3 | RFC3 | 0.9 | 1.3 | 1.0 | 1.1 | 0.9 | No |
| 1089 | replication factor C subunit 4 | RFC4 | 1.4 | 2.3 | 1.2 | 2.4 | 1.1 | No |
| 1090 | replication factor C subunit 5 | RFC5 | 0.7 | 1.4 | 0.5 | 1.0 | 0.7 | No |
| 1091 | repulsive guidance molecule BMP co-receptor a | RGMA | -1.1 | -1.0 | -0.8 | -0.9 | -1.2 | No |
| 1092 | regulator of G protein signaling 11 | RGS11 | -0.9 | -1.7 | -0.8 | 0.8 | -1.1 | No |
| 1093 | regulator of G protein signaling 5 | RGS5 | -1.4 | -1.8 | -1.8 | -2.1 | -2.1 | No |
| 1094 | Rho related BTB domain containing 2 | RHOBTB2 | -8.3 | -5.8 | -2.6 | -2.1 | -10.9 | No |
| 1095 | rhophilin Rho GTPase binding protein 2 | RHPN2 | -0.3 | -1.4 | -1.5 | -1.4 | -1.1 | No |
| 1096 | RIC8 guanine nucleotide exchange factor A | RIC8A | -1.1 | -0.8 | -1.0 | -0.5 | -0.6 | No |
| 1097 | ribosomal modification protein rimK like family member A | RIMKLA | 1.5 | 1.6 | 1.3 | 1.9 | 1.5 | No |
| 1098 | RAD50 interactor 1 | RINT1 | 0.7 | 0.7 | 0.9 | 0.8 | 1.1 | No |
| 1099 | RHO family interacting cell polarization regulator 1 | RIPOR1 | 3.0 | 2.4 | 1.7 | 0.9 | 1.0 | No |
| 1100 | ribonuclease H2 subunit A | RNASEH2A | 1.6 | 2.3 | 1.4 | 1.6 | 1.1 | No |
| 1101 | ring finger protein 11 | RNF11 | -1.5 | -1.7 | -1.2 | -0.7 | -0.9 | No |
| 1102 | ring finger protein 13 | RNF13 | -6.0 | -1.2 | -0.6 | -0.2 | -1.1 | No |
| 1103 | ring finger protein 144A | RNF144A | -1.8 | -1.4 | -1.7 | -0.9 | -0.8 | No |
| 1104 | ring finger protein 148 | RNF148 | -0.9 | -1.2 | -1.0 | -1.4 | -1.0 | No |
| 1105 | ring finger protein 24 | RNF24 | -8.8 | -1.5 | -7.8 | -0.7 | -0.7 | No |
| 1106 | arginyl aminopeptidase | RNPEP | 1.1 | 1.7 | 1.6 | 3.0 | 1.5 | No |
| 1107 | roundabout guidance receptor 3 | ROBO3 | 1.0 | 0.6 | 2.6 | 5.1 | 4.4 | No |
| 1108 | replication protein A1 | RPA1 | 1.3 | 1.9 | 1.2 | 1.7 | 1.2 | No |
| 1109 | replication protein A2 | RPA2 | 1.6 | 2.2 | 1.5 | 1.9 | 1.4 | No |
| 1110 | replication protein A3 | RPA3 | 1.6 | 2.2 | 1.6 | 2.1 | 1.5 | No |
| 1111 | ribosome production factor 2 homolog | RPF2 | 1.4 | 1.2 | 0.5 | 0.0 | -0.1 | No |
| 1112 | ribose 5-phosphate isomerase A | RPIA | 1.8 | 1.1 | 0.8 | 0.6 | -0.1 | No |
| 1113 | ribosomal protein L7 | RPL7 | -0.7 | -0.7 | -0.9 | -1.0 | -0.6 | No |
| 1114 | ribosomal protein L7 like 1 | RPL7L1 | 1.8 | 1.6 | 1.1 | 0.7 | 0.6 | No |
| 1115 | ribophorin I | RPN1 | 0.8 | 1.2 | 0.8 | 1.1 | 1.0 | No |
| 1116 | ribophorin II | RPN2 | 0.9 | 0.4 | 1.4 | 1.6 | 1.3 | No |
| 1117 | ribonuclease P/MRP subunit p14 | RPP14 | 0.9 | 1.1 | 0.3 | 0.0 | -0.2 | No |
| 1118 | ribonuclease P/MRP subunit p40 | RPP40 | 1.0 | 0.7 | 0.4 | 0.6 | 0.4 | No |
| 1119 | ribosomal protein S16 | RPS16 | -0.7 | -0.6 | -0.9 | -1.0 | -0.7 | No |
| 1120 | ribosomal RNA processing 15 homolog | RRP15 | 1.6 | 1.3 | 0.8 | 0.3 | 0.0 | No |
| 1121 | ribosomal RNA processing 36 | RRP36 | 1.2 | 0.9 | 0.6 | 0.5 | -0.1 | No |
| 1122 | radical S-adenosyl methionine domain containing 2 | RSAD2 | 1.0 | 1.5 | 2.3 | 0.4 | 2.6 | No |
| 1123 | REM2 and RAB like small GTPase 1 | RSG1 | 1.3 | 1.5 | 1.3 | 1.3 | 0.9 | No |
| 1124 | RNA 3'-terminal phosphate cyclase | RTCA | 1.4 | 1.4 | 1.1 | 1.0 | 0.8 | No |
| 1125 | RNA transcription, translation and transport factor | RTRAF | 1.0 | 0.7 | 0.7 | 0.7 | 0.3 | No |
| 1126 | RuvB like AAA ATPase 1 | RUVBL1 | 1.3 | 1.4 | 0.8 | 0.8 | 0.4 | No |
| 1127 | RuvB like AAA ATPase 2 | RUVBL2 | 1.8 | 1.6 | 1.2 | 0.8 | 0.5 | No |
| 1128 | RWD domain containing 1 | RWDD1 | 1.0 | 1.3 | 0.7 | 0.8 | 0.2 | No |
| 1129 | RWD domain containing 4A | RWDD4A | 1.1 | 1.0 | 0.8 | 0.3 | -1.0 | No |
| 1130 | retinoid X receptor gamma | RXRG | -0.9 | -0.5 | -0.7 | -0.5 | -0.3 | No |
| 1131 | sphingosine-1-phosphate receptor 5 | S1PR5 | 1.2 | 2.0 | 0.8 | 0.1 | 0.8 | No |
| 1132 | serum amyloid A2 | SAA2 | 1.6 | 2.4 | 2.1 | 1.5 | 1.4 | No |
| 1133 | SUMO1 activating enzyme subunit 1 | SAE1 | 2.9 | 3.2 | 2.6 | 0.0 | 2.0 | No |
| 1134 | scaffold attachment factor B | SAFB | -0.8 | -0.2 | -1.0 | -0.7 | -0.7 | No |
| 1135 | SAM domain, SH3 domain and nuclear localization signals 1 | SAMSN1 | 1.4 | 0.8 | 0.9 | 0.9 | 0.5 | No |
| 1136 | SAS-6 centriolar assembly protein | SASS6 | 1.2 | 2.0 | 1.0 | 0.8 | 1.1 | No |
| 1137 | scinderin | SCIN | -1.2 | -1.8 | -1.7 | -3.5 | -2.2 | No |
| 1138 | short coiled-coil protein | SCOC | 0.5 | 0.4 | 1.4 | 2.0 | 1.6 | No |
| 1139 | stromal cell derived factor 2 | SDF2 | 1.2 | 5.3 | 4.5 | 6.8 | -4.8 | No |
| 1140 | short chain dehydrogenase/reductase family 16C member 5 | SDR16C5 | 1.0 | 0.9 | 0.6 | 0.6 | 0.0 | No |
| 1141 | SEC11 homolog A, signal peptidase complex subunit | SEC11A | 0.9 | 1.1 | 1.2 | 1.9 | 1.3 | No |
| 1142 | SEC13 homolog, nuclear pore and COPII coat complex component | SEC13 | 1.3 | 1.3 | 0.9 | 1.2 | 0.9 | No |
| 1143 | selenoprotein N | SELENON | 1.3 | 1.4 | 2.7 | 4.0 | 3.9 | No |
| 1144 | selenoprotein V | SELENOV | -1.1 | -0.8 | -1.0 | -0.8 | -0.2 | No |
| 1145 | SEM1, 26S proteasome complex subunit | SEM1 | 1.0 | 1.0 | 0.8 | 0.9 | 0.8 | No |
| 1146 | semaphorin 3F | SEMA3F | -0.6 | -1.5 | -0.3 | 0.0 | -0.5 | No |
| 1147 | selenophosphate synthetase 1 | SEPHS1 | 7.7 | 1.1 | 1.7 | -0.2 | 0.3 | No |
| 1148 | Sep (O-phosphoserine) tRNA:Sec (selenocysteine) tRNA synthase | SEPSECS | 1.2 | 1.3 | 0.9 | 0.7 | 0.5 | No |
| 1149 | serine incorporator 1 | SERINC1 | -0.7 | -1.1 | -0.2 | -0.1 | -0.3 | No |
| 1150 | serpin family E member 1 | SERPINE1 | 2.9 | 2.8 | 2.2 | 1.3 | 2.5 | No |
| 1151 | sestrin 1 | SESN1 | -2.3 | -1.8 | -0.7 | -0.8 | -1.5 | No |
| 1152 | sestrin 2 | SESN2 | 1.4 | 1.0 | 1.1 | 0.1 | 0.2 | No |
| 1153 | SET nuclear proto-oncogene | SET | 0.9 | 1.0 | 0.5 | 0.7 | 0.2 | No |
| 1154 | SET domain containing 1B | SETD1B | -0.6 | -0.5 | -0.8 | -0.8 | -1.0 | No |
| 1155 | SET domain containing 6 | SETD6 | 1.2 | 1.2 | 0.5 | 0.5 | 0.1 | No |
| 1156 | SET domain containing lysine methyltransferase 7 | SETD7 | -0.7 | -0.4 | -1.2 | -0.6 | -0.3 | No |
| 1157 | splicing factor 3b subunit 4 | SF3B4 | 1.4 | 1.4 | 1.0 | 0.8 | 0.5 | No |
| 1158 | splicing factor 3b subunit 5 | SF3B5 | 1.4 | 1.7 | 1.2 | 1.4 | 0.9 | No |
| 1159 | stratifin | SFN | 1.3 | 1.7 | 1.1 | 0.4 | 0.2 | No |
| 1160 | SH3 domain binding glutamate rich protein like 2 | SH3BGRL2 | -1.1 | -1.6 | -1.6 | -1.0 | -1.0 | No |
| 1161 | SH3 domain binding protein 4 | SH3BP4 | 7.1 | 5.8 | 6.7 | 12.2 | 11.3 | No |
| 1162 | SHANK associated RH domain interactor | SHARPIN | -1.0 | -0.9 | -1.0 | -1.0 | -0.9 | No |
| 1163 | Src homology 2 domain containing transforming protein D | SHD | 1.2 | 0.6 | 1.2 | 2.3 | 1.5 | No |
| 1164 | serine hydroxymethyltransferase 1 | SHMT1 | 1.3 | 1.9 | 1.1 | 0.9 | 0.5 | No |
| 1165 | SHQ1, H/ACA ribonucleoprotein assembly factor | SHQ1 | -0.8 | -0.8 | -0.8 | -0.3 | -1.0 | No |
| 1166 | siah E3 ubiquitin protein ligase family member 3 | SIAH3 | -0.7 | -0.6 | -0.4 | -0.8 | -1.1 | No |
| 1167 | sigma non-opioid intracellular receptor 1 | SIGMAR1 | 1.1 | 1.0 | 0.3 | 0.4 | 0.4 | No |
| 1168 | SIX homeobox 1 | SIX1 | -1.2 | -1.1 | -1.5 | -2.7 | -1.7 | No |
| 1169 | SIX homeobox 4 | SIX4 | -0.8 | -1.2 | -1.7 | -2.3 | -2.1 | No |
| 1170 | S-phase kinase associated protein 2 | SKP2 | 1.1 | 2.3 | 1.6 | 2.0 | 1.5 | No |
| 1171 | solute carrier family 14 member 2 | SLC14A2 | -2.6 | -3.8 | -1.1 | -4.0 | -2.2 | No |
| 1172 | solute carrier family 17 member 9 | SLC17A9 | 0.9 | 0.7 | 0.7 | 0.5 | -0.5 | No |
| 1173 | solute carrier family 1 member 4 | SLC1A4 | 2.7 | 0.3 | 4.2 | 2.4 | 2.4 | No |
| 1174 | solute carrier family 22 member 2 | SLC22A2 | -1.0 | -1.8 | -1.2 | -1.5 | -1.1 | No |
| 1175 | solute carrier family 23 member 1 | SLC23A1 | 0.9 | 2.3 | 1.2 | 2.3 | 0.6 | No |
| 1176 | solute carrier family 24 member 5 | SLC24A5 | -1.9 | -1.2 | -1.5 | 0.4 | 1.3 | No |
| 1177 | solute carrier family 25 member 1 | SLC25A1 | 0.5 | 1.1 | 1.1 | 1.6 | 1.2 | No |
| 1178 | solute carrier family 25 member 11 | SLC25A11 | 1.0 | 1.0 | 0.8 | 1.2 | 0.7 | No |
| 1179 | solute carrier family 25 member 25 | SLC25A25 | 2.4 | 0.6 | 1.8 | 0.4 | 0.7 | No |
| 1180 | solute carrier family 25 member 26 | SLC25A26 | 1.2 | 1.0 | 0.2 | 0.4 | 0.0 | No |
| 1181 | solute carrier family 25 member 32 | SLC25A32 | 1.7 | 1.1 | 1.3 | 1.0 | 0.1 | No |
| 1182 | solute carrier family 25 member 36 | SLC25A36 | -1.7 | -0.7 | -1.3 | -1.2 | -1.2 | No |
| 1183 | solute carrier family 30 member 1 | SLC30A1 | 1.1 | 0.5 | 0.7 | 0.4 | 0.5 | No |
| 1184 | solute carrier family 34 member 2 | SLC34A2 | 0.5 | 0.9 | 1.1 | 1.0 | 0.5 | No |
| 1185 | solute carrier family 35 member E3 | SLC35E3 | 1.5 | 1.0 | 1.7 | 2.4 | 1.6 | No |
| 1186 | solute carrier family 35 member F2 | SLC35F2 | 0.5 | 0.9 | 0.4 | 0.9 | 0.2 | No |
| 1187 | solute carrier family 38 member 4 | SLC38A4 | -1.5 | -2.8 | -2.3 | -1.0 | -0.6 | No |
| 1188 | solute carrier family 38 member 5 | SLC38A5 | -0.9 | -1.1 | -0.8 | -1.3 | -1.0 | No |
| 1189 | solute carrier family 3 member 2 | SLC3A2 | 2.1 | 1.2 | 1.3 | 1.7 | 1.8 | No |
| 1190 | solute carrier family 43 member 1 | SLC43A1 | 1.3 | 1.2 | 0.9 | 0.7 | 0.7 | No |
| 1191 | solute carrier family 45 member 2 | SLC45A2 | -0.7 | -1.6 | -1.0 | -1.7 | -0.6 | No |
| 1192 | solute carrier family 6 member 4 | SLC6A4 | -1.0 | -1.3 | -1.9 | -1.6 | -1.1 | No |
| 1193 | solute carrier family 7 member 7 | SLC7A7 | 1.1 | 1.3 | 1.1 | 0.2 | -0.4 | No |
| 1194 | solute carrier organic anion transporter family member 2B1 | SLCO2B1 | -1.6 | -1.9 | -1.7 | -1.9 | -1.9 | No |
| 1195 | solute carrier organic anion transporter family member 3A1 | SLCO3A1 | 1.5 | 0.6 | 1.9 | 1.7 | 2.1 | No |
| 1196 | sarcolemma associated protein | SLMAP | -0.2 | 0.8 | 1.6 | 1.4 | 1.2 | No |
| 1197 | SWI/SNF related, matrix associated, actin dependent regulator of chromatin, subfamily e, member 1 | SMARCE1 | 0.6 | 1.0 | 1.0 | 1.5 | 0.0 | No |
| 1198 | structural maintenance of chromosomes 4 | SMC4 | 0.8 | 2.1 | 1.7 | 2.1 | 1.7 | No |
| 1199 | survival of motor neuron 1, telomeric | SMN1 | 1.1 | 1.0 | 0.5 | 0.3 | -0.2 | No |
| 1200 | small muscle protein X-linked | SMPX | -1.0 | -1.3 | -2.1 | -15.3 | -15.4 | No |
| 1201 | smoothelin like 2 | SMTNL2 | 4.0 | 4.0 | 3.8 | 2.8 | 2.6 | No |
| 1202 | SMYD family member 5 | SMYD5 | 2.6 | 2.0 | 1.3 | 0.9 | -0.1 | No |
| 1203 | snail family transcriptional repressor 3 | SNAI3 | -2.9 | -2.8 | -2.4 | -2.8 | -1.6 | No |
| 1204 | small nuclear ribonucleoprotein U5 subunit 40 | SNRNP40 | 1.4 | 1.3 | 1.4 | 1.2 | 0.3 | No |
| 1205 | small nuclear ribonucleoprotein polypeptides B and B1 | SNRPB | 1.0 | 1.2 | 0.8 | 1.0 | 0.8 | No |
| 1206 | small nuclear ribonucleoprotein polypeptide C | SNRPC | 1.2 | 1.6 | 1.1 | 1.1 | 0.7 | No |
| 1207 | small nuclear ribonucleoprotein D1 polypeptide | SNRPD1 | 1.9 | 2.1 | 1.4 | 1.4 | 1.2 | No |
| 1208 | small nuclear ribonucleoprotein D2 polypeptide | SNRPD2 | 1.9 | 1.9 | 1.3 | 1.3 | 0.7 | No |
| 1209 | small nuclear ribonucleoprotein D3 polypeptide | SNRPD3 | 1.3 | 1.2 | 0.9 | 1.1 | 0.3 | No |
| 1210 | small nuclear ribonucleoprotein polypeptide E | SNRPE | 1.9 | 2.0 | 1.2 | 1.5 | 1.1 | No |
| 1211 | sorting nexin 18 | SNX18 | 1.6 | 0.9 | 1.8 | 1.9 | 1.3 | No |
| 1212 | sorting nexin 24 | SNX24 | 1.4 | 0.8 | 1.3 | 0.8 | 0.1 | No |
| 1213 | suppressor of cytokine signaling 3 | SOCS3 | 2.4 | 2.4 | 2.5 | 2.3 | 1.8 | No |
| 1214 | sclerostin domain containing 1 | SOSTDC1 | -2.0 | -2.8 | -1.4 | -3.1 | -3.9 | No |
| 1215 | SRY-box 1 | SOX1 | 0.0 | -1.7 | -1.0 | -2.0 | -1.7 | No |
| 1216 | SRY-box 3 | SOX3 | -1.9 | -1.8 | -2.5 | -1.7 | -0.8 | No |
| 1217 | SPC24, NDC80 kinetochore complex component | SPC24 | 0.7 | 2.3 | 1.8 | 2.4 | 2.0 | No |
| 1218 | signal peptidase complex subunit 1 | SPCS1 | 0.3 | 0.6 | 0.4 | 1.0 | 0.8 | No |
| 1219 | spire type actin nucleation factor 1 | SPIRE1 | -1.1 | -1.0 | -1.4 | -1.4 | -1.2 | No |
| 1220 | sphingolipid transporter 2 | SPNS2 | -0.6 | -0.9 | -1.3 | -1.4 | -0.9 | No |
| 1221 | sprouty RTK signaling antagonist 4 | SPRY4 | 1.3 | 0.5 | 1.7 | 2.3 | 1.6 | No |
| 1222 | SPRY domain containing 4 | SPRYD4 | 1.3 | 1.5 | 0.9 | 1.2 | 0.5 | No |
| 1223 | SPT2 chromatin protein domain containing 1 | SPTY2D1 | -2.2 | -1.2 | -1.4 | -1.8 | -1.5 | No |
| 1224 | SRC proto-oncogene, non-receptor tyrosine kinase | SRC | 1.3 | 0.8 | 1.5 | 0.2 | 0.6 | No |
| 1225 | steroid 5 alpha-reductase 2 | SRD5A2 | -0.4 | -0.5 | -0.7 | -1.3 | -0.7 | No |
| 1226 | steroid 5 alpha-reductase 3 | SRD5A3 | 1.1 | 0.8 | 0.5 | 0.7 | 0.2 | No |
| 1227 | splicing regulatory glutamic acid and lysine rich protein 1 | SREK1 | -1.1 | -0.6 | -1.1 | -2.3 | -1.0 | No |
| 1228 | SREK1 interacting protein 1 | SREK1IP1 | 0.6 | 1.0 | 0.8 | 1.1 | 0.7 | No |
| 1229 | sorcin | SRI | 1.0 | 1.1 | 0.9 | 0.9 | 0.6 | No |
| 1230 | signal recognition particle 19 | SRP19 | 0.9 | 0.9 | 0.7 | 0.9 | 0.2 | No |
| 1231 | signal recognition particle 9 | SRP9 | 0.8 | 1.0 | 0.6 | 0.7 | 0.4 | No |
| 1232 | SRSF protein kinase 1 | SRPK1 | -1.6 | -1.1 | -1.2 | -0.7 | -0.8 | No |
| 1233 | serine and arginine rich splicing factor 6 | SRSF6 | -2.2 | -0.7 | -1.7 | -0.8 | -1.1 | No |
| 1234 | Sjogren syndrome antigen B | SSB | 1.7 | 1.3 | 0.9 | 0.8 | 0.1 | No |
| 1235 | single stranded DNA binding protein 1 | SSBP1 | 1.3 | 1.2 | 0.6 | 0.8 | 0.2 | No |
| 1236 | signal sequence receptor subunit 3 | SSR3 | 1.3 | 1.2 | 1.1 | 1.5 | 1.3 | No |
| 1237 | signal sequence receptor subunit 4 | SSR4 | 0.9 | 1.0 | 0.9 | 1.7 | 1.3 | No |
| 1238 | suppression of tumorigenicity 14 | ST14 | -1.5 | -1.4 | -1.5 | -2.3 | -1.9 | No |
| 1239 | signal transducing adaptor family member 2 | STAP2 | 1.8 | 0.8 | 1.6 | 0.7 | 0.5 | No |
| 1240 | signal transducer and activator of transcription 1 | STAT1 | -1.1 | -1.2 | -0.8 | -0.8 | -0.7 | No |
| 1241 | stanniocalcin 2 | STC2 | 1.4 | 0.8 | 0.5 | -0.3 | 0.4 | No |
| 1242 | STIL, centriolar assembly protein | STIL | 0.9 | 1.9 | 8.8 | 2.9 | 0.1 | No |
| 1243 | stress induced phosphoprotein 1 | STIP1 | 0.9 | 0.8 | 0.5 | 0.0 | 0.3 | No |
| 1244 | serine/threonine kinase 16 | STK16 | 1.1 | 0.5 | 0.5 | 0.8 | 0.7 | No |
| 1245 | serine/threonine kinase 35 | STK35 | -1.6 | -1.4 | -1.0 | -0.8 | -1.1 | No |
| 1246 | stathmin 1 | STMN1 | 0.4 | 1.2 | 0.8 | 0.8 | 0.6 | No |
| 1247 | syntaxin 11 | STX11 | -0.7 | -0.7 | -0.9 | -1.6 | -0.9 | No |
| 1248 | syntaxin binding protein 1 | STXBP1 | -1.0 | -0.3 | -0.8 | -0.9 | -1.8 | No |
| 1249 | succinate-CoA ligase ADP-forming beta subunit | SUCLA2 | 1.2 | 1.1 | 0.7 | 0.7 | 0.1 | No |
| 1250 | succinate-CoA ligase alpha subunit | SUCLG1 | 1.1 | 1.1 | 0.6 | 0.6 | 0.4 | No |
| 1251 | sulfatase 1 | SULF1 | 1.2 | 0.9 | 1.9 | 1.6 | 1.9 | No |
| 1252 | sulfotransferase family, cytosolic, 1C, member 1 | SULT1C1 | -2.2 | -2.5 | -2.2 | -1.2 | -2.5 | No |
| 1253 | sulfotransferase family 1C member 2 | SULT1C2 | -2.0 | -1.7 | -1.0 | -0.6 | -2.6 | No |
| 1254 | small ubiquitin-like modifier 3 | SUMO3 | 0.4 | 0.8 | 0.5 | 1.0 | 0.6 | No |
| 1255 | supervillin | SVIL | -0.9 | -1.6 | -1.4 | -1.0 | -1.2 | No |
| 1256 | switching B cell complex subunit SWAP70 | SWAP70 | -1.0 | -1.3 | -0.8 | -0.4 | -1.3 | No |
| 1257 | synaptophysin like 2 | SYPL2 | 1.5 | 0.7 | 1.3 | 1.1 | 0.8 | No |
| 1258 | synaptotagmin 11 | SYT11 | -0.8 | -1.5 | -1.3 | -1.7 | -2.2 | No |
| 1259 | tachykinin receptor 3 | TACR3 | -0.7 | -2.4 | -1.5 | -0.7 | -0.7 | No |
| 1260 | transgelin | TAGLN | 1.2 | 1.4 | 0.4 | -0.1 | 0.0 | No |
| 1261 | transport and golgi organization 2 homolog | TANGO2 | 2.2 | 2.0 | 1.5 | 2.2 | -6.7 | No |
| 1262 | transporter 2, ATP binding cassette subfamily B member | TAP2 | -0.6 | -0.7 | 0.0 | -0.2 | -1.3 | No |
| 1263 | Tax1 binding protein 3 | TAX1BP3 | 1.0 | 0.8 | 0.8 | 0.0 | 0.0 | No |
| 1264 | TBC1 domain family member 19 | TBC1D19 | 1.3 | 0.9 | 1.2 | 2.3 | 0.9 | No |
| 1265 | tubulin folding cofactor A | TBCA | 1.0 | 1.1 | 0.9 | 0.9 | 0.7 | No |
| 1266 | transducin beta like 3 | TBL3 | 1.7 | 1.6 | 1.2 | -0.3 | -0.4 | No |
| 1267 | T-box 2 | TBX2 | -1.0 | -1.2 | -1.1 | -0.9 | -0.7 | No |
| 1268 | transcription factor 7 | TCF7 | 0.7 | 0.7 | 2.1 | 3.1 | 2.7 | No |
| 1269 | T cell leukemia translocation altered | TCTA | 0.9 | 1.5 | 1.0 | 1.1 | 0.9 | No |
| 1270 | transcription factor A, mitochondrial | TFAM | 0.7 | 1.0 | 1.0 | 1.2 | 0.6 | No |
| 1271 | transcription factor AP-2 alpha | TFAP2A | -1.0 | -0.7 | -0.6 | 0.1 | -0.5 | No |
| 1272 | transcription factor AP-2 epsilon | TFAP2E | -1.3 | -2.8 | -1.7 | -2.7 | -2.1 | No |
| 1273 | TGFB induced factor homeobox 1 | TGIF1 | 0.6 | 0.8 | 1.0 | 1.0 | 0.9 | No |
| 1274 | Threonine aldolase 1 | THA1 | 0.5 | 0.8 | 0.7 | 2.0 | 1.4 | No |
| 1275 | THAP domain containing 1 | THAP1 | -0.9 | -1.1 | -0.8 | -0.4 | -0.5 | No |
| 1276 | thrombospondin 3 | THBS3 | 1.5 | 1.2 | 1.8 | 2.1 | 2.8 | No |
| 1277 | THO complex 3 | THOC3 | 0.9 | 0.8 | 0.6 | 0.7 | 0.4 | No |
| 1278 | THO complex 6 | THOC6 | 3.4 | 3.9 | 3.4 | 4.1 | 0.8 | No |
| 1279 | thimet oligopeptidase 1 | THOP1 | 9.0 | 1.8 | 1.6 | 16.9 | 15.3 | No |
| 1280 | thymocyte nuclear protein 1 | THYN1 | 2.2 | 2.2 | 2.3 | 3.5 | 3.4 | No |
| 1281 | TIA1 cytotoxic granule associated RNA binding protein like 1 | TIAL1 | -1.2 | -0.7 | -0.9 | -0.8 | -0.8 | No |
| 1282 | TP53 induced glycolysis regulatory phosphatase | TIGAR | 1.6 | 1.6 | 1.5 | 0.6 | -0.5 | No |
| 1283 | translocase of inner mitochondrial membrane 10 | TIMM10 | 2.3 | 2.3 | 1.5 | 1.3 | 0.8 | No |
| 1284 | translocase of inner mitochondrial membrane 13 | TIMM13 | 1.7 | 2.1 | 1.1 | 1.1 | 0.6 | No |
| 1285 | translocase of inner mitochondrial membrane 17A | TIMM17A | 1.2 | 1.1 | 0.5 | 0.2 | 0.3 | No |
| 1286 | translocase of inner mitochondrial membrane 8 homolog B | TIMM8B | 1.6 | 1.6 | 0.9 | 0.4 | 0.2 | No |
| 1287 | TIMP metallopeptidase inhibitor 2 | TIMP2 | 5.1 | 5.4 | 3.7 | 4.3 | 3.4 | No |
| 1288 | tubulointerstitial nephritis antigen like 1 | TINAGL1 | 3.1 | 2.7 | 3.5 | 2.3 | 2.9 | No |
| 1289 | translation machinery associated 16 homolog | TMA16 | 1.0 | 0.8 | 0.3 | 0.6 | 0.0 | No |
| 1290 | transmembrane p24 trafficking protein 10 | TMED10 | 0.5 | 0.9 | 0.4 | 0.6 | 0.5 | No |
| 1291 | transmembrane p24 trafficking protein 2 | TMED2 | 0.6 | 0.8 | 0.5 | 0.6 | 0.5 | No |
| 1292 | transmembrane p24 trafficking protein 5 | TMED5 | 1.1 | 0.9 | 0.8 | 0.8 | 0.8 | No |
| 1293 | transmembrane protein 144 | TMEM144 | -0.8 | -1.0 | -0.6 | -0.3 | -0.3 | No |
| 1294 | transmembrane protein 147 | TMEM147 | 1.0 | 1.2 | 0.4 | 0.7 | 0.5 | No |
| 1295 | transmembrane protein 161A | TMEM161A | 1.2 | 0.8 | 1.5 | 1.1 | 0.7 | No |
| 1296 | transmembrane protein 167 | TMEM167 | 0.5 | 0.9 | 0.4 | 0.9 | 0.5 | No |
| 1297 | transmembrane protein 183A | TMEM183A | 1.1 | 1.4 | 0.9 | 0.8 | 0.8 | No |
| 1298 | transmembrane protein 199 | TMEM199 | 0.7 | 0.9 | 0.9 | 0.5 | 0.4 | No |
| 1299 | transmembrane protein 258 | TMEM258 | 0.6 | 1.1 | 0.6 | 1.2 | 0.9 | No |
| 1300 | transmembrane protein 38A | TMEM38A | -2.4 | -2.8 | -1.7 | -2.3 | -1.0 | No |
| 1301 | transmembrane protein 39B | TMEM39B | -1.2 | -0.8 | -1.0 | -1.1 | -1.0 | No |
| 1302 | transmembrane protein 44 | TMEM44 | 2.2 | 2.9 | 1.0 | 0.0 | 1.9 | No |
| 1303 | transmembrane protein 59 | TMEM59 | -1.0 | -0.8 | -0.5 | -0.5 | -0.9 | No |
| 1304 | transmembrane protein 9 | TMEM9 | -6.5 | -6.5 | -5.9 | -13.3 | -15.7 | No |
| 1305 | transmembrane serine protease 13 | TMPRSS13 | -0.6 | -0.8 | -1.6 | -1.7 | -1.1 | No |
| 1306 | TNF receptor superfamily member 19 | TNFRSF19 | 0.6 | 0.5 | 1.6 | 2.0 | 1.1 | No |
| 1307 | TNF superfamily member 10 | TNFSF10 | -1.2 | -1.1 | -0.3 | -0.2 | -0.6 | No |
| 1308 | TRAF2 and NCK interacting kinase | TNIK | -1.6 | -2.2 | -2.6 | 0.0 | -2.8 | No |
| 1309 | tenascin N | TNN | -3.4 | -3.0 | -2.9 | -1.7 | 0.4 | No |
| 1310 | troponin C1, slow skeletal and cardiac type | TNNC1 | 2.6 | 2.0 | 3.1 | 2.2 | 3.1 | No |
| 1311 | troponin T2, cardiac type | TNNT2 | 1.5 | 1.7 | 0.7 | -0.2 | 0.3 | No |
| 1312 | transducer of ERBB2, 1 | TOB1 | -0.7 | -1.2 | -1.8 | -1.5 | -0.7 | No |
| 1313 | translocase of outer mitochondrial membrane 22 | TOMM22 | 1.3 | 1.3 | 0.4 | 0.5 | 0.2 | No |
| 1314 | translocase of outer mitochondrial membrane 40 | TOMM40 | 2.6 | 2.4 | 1.7 | 1.8 | 1.1 | No |
| 1315 | torsin family 1 member A | TOR1A | 1.6 | 1.1 | 1.4 | 1.0 | 1.7 | No |
| 1316 | TPD52 like 1 | TPD52L1 | -1.6 | -2.1 | -1.7 | -1.5 | -1.3 | No |
| 1317 | tryptophan hydroxylase 1 | TPH1 | -1.8 | -0.9 | -1.5 | -1.9 | -3.0 | No |
| 1318 | tryptophan hydroxylase 2 | TPH2 | -1.2 | -1.1 | -0.9 | -1.3 | -0.1 | No |
| 1319 | tubulin polymerization promoting protein family member 3 | TPPP3 | -1.3 | -0.7 | -0.7 | -0.6 | 0.9 | No |
| 1320 | TP53RK binding protein | TPRKB | 0.8 | 0.4 | 0.7 | 1.1 | 0.6 | No |
| 1321 | transmembrane phosphatase with tensin homology | TPTE | 0.4 | 0.6 | 0.7 | 1.1 | 1.1 | No |
| 1322 | transformer 2 alpha homolog | TRA2A | -0.7 | -0.6 | -0.6 | -1.3 | -0.8 | No |
| 1323 | TraB domain containing | TRABD | 1.1 | 0.9 | 0.9 | 0.8 | 0.4 | No |
| 1324 | TRAF interacting protein | TRAIP | 1.1 | 2.4 | 1.5 | 2.7 | 2.3 | No |
| 1325 | translocation associated membrane protein 1 like 1 | TRAM1L1 | 0.8 | 1.1 | 0.7 | 0.5 | 0.7 | No |
| 1326 | translocation associated membrane protein 2 | TRAM2 | -0.3 | 0.6 | 0.5 | 1.4 | 1.0 | No |
| 1327 | trafficking protein particle complex 6A | TRAPPC6A | 0.5 | 0.8 | 0.8 | 1.2 | 0.4 | No |
| 1328 | TP53 regulated inhibitor of apoptosis 1 | TRIAP1 | 0.7 | 1.3 | 0.7 | 0.9 | 0.4 | No |
| 1329 | tripartite motif containing 35 | TRIM35 | -0.3 | -0.6 | -0.2 | -2.4 | -0.7 | No |
| 1330 | thyroid hormone receptor interactor 13 | TRIP13 | 1.7 | 3.4 | 2.9 | 3.4 | 2.1 | No |
| 1331 | tRNA methyltransferase 2 homolog A | TRMT2A | 1.2 | 1.3 | 1.1 | 1.4 | 0.8 | No |
| 1332 | tRNA methyltransferase 61A | TRMT61A | 1.1 | 0.4 | 0.7 | 0.5 | 0.2 | No |
| 1333 | TruB pseudouridine synthase family member 1 | TRUB1 | 1.6 | 1.3 | 1.2 | 0.9 | 0.3 | No |
| 1334 | Ts translation elongation factor, mitochondrial | TSFM | 1.8 | 1.7 | 1.3 | 1.1 | 0.5 | No |
| 1335 | translin associated factor X interacting protein 1 | TSNAXIP1 | -2.5 | -2.2 | -2.2 | -1.3 | -1.4 | No |
| 1336 | tetraspanin 12 | TSPAN12 | -1.4 | -1.4 | -1.8 | -1.8 | -1.2 | No |
| 1337 | tetraspanin 3 | TSPAN3 | -0.9 | -1.0 | -0.8 | -1.0 | -1.1 | No |
| 1338 | tetraspanin 6 | TSPAN6 | 0.7 | 0.5 | 1.4 | 2.4 | 1.9 | No |
| 1339 | tetraspanin 7 | TSPAN7 | -1.6 | -2.2 | -2.2 | -2.0 | -2.4 | No |
| 1340 | TSR1, ribosome maturation factor | TSR1 | 2.3 | 1.8 | 1.5 | 0.7 | -0.1 | No |
| 1341 | tetratricopeptide repeat domain 14 | TTC14 | -1.3 | -1.0 | -0.9 | -0.4 | -0.8 | No |
| 1342 | TTK protein kinase | TTK | 1.4 | 2.8 | 2.7 | 2.9 | 1.1 | No |
| 1343 | tweety family member 3 | TTYH3 | -0.3 | 0.2 | 2.1 | 3.3 | 2.4 | No |
| 1344 | tubulin, beta 5 class I | TUBB5 | 8.8 | 2.6 | 11.1 | 17.7 | 17.0 | No |
| 1345 | tubulin gamma complex associated protein 3 | TUBGCP3 | 1.4 | 1.8 | 1.4 | 2.9 | 2.6 | No |
| 1346 | terminal uridylyl transferase 1, U6 snRNA-specific | TUT1 | 1.1 | 1.2 | 1.0 | 1.3 | 0.4 | No |
| 1347 | thioredoxin domain containing 17 | TXNDC17 | 0.7 | 0.9 | 0.9 | 1.2 | 1.0 | No |
| 1348 | thioredoxin domain containing 2 | TXNDC2 | 3.2 | 3.2 | 2.9 | 2.6 | 2.4 | No |
| 1349 | thymidylate synthetase | TYMS | 1.5 | 2.6 | 2.0 | 2.4 | 2.0 | No |
| 1350 | U2 snRNP associated SURP domain containing | U2SURP | -1.4 | -0.7 | -1.4 | -1.0 | -1.1 | No |
| 1351 | UBA domain containing 1 | UBAC1 | -0.9 | -0.8 | -1.2 | -0.8 | -0.9 | No |
| 1352 | ubiquitin conjugating enzyme E2 C | UBE2C | 1.1 | 2.3 | 2.1 | 3.7 | 3.2 | No |
| 1353 | ubiquitin conjugating enzyme E2 N | UBE2N | 0.8 | 1.3 | 0.7 | 0.6 | 0.4 | No |
| 1354 | ubiquilin 4 | UBQLN4 | 0.8 | 1.0 | 0.6 | 0.3 | 0.4 | No |
| 1355 | ubiquitin protein ligase E3 component n-recognin 7 (putative) | UBR7 | 0.8 | 1.1 | 0.8 | 0.9 | 0.4 | No |
| 1356 | UBX domain protein 6 | UBXN6 | -0.9 | -0.8 | -0.3 | -0.5 | -0.8 | No |
| 1357 | uridine-cytidine kinase 2 | UCK2 | 1.1 | 1.0 | 0.7 | 1.0 | 0.9 | No |
| 1358 | upper zone of growth plate and cartilage matrix associated | UCMA | -1.6 | -2.8 | -1.7 | -2.8 | -1.1 | No |
| 1359 | ubiquitin-fold modifier conjugating enzyme 1 | UFC1 | 1.1 | 1.0 | 0.5 | 1.0 | 0.6 | No |
| 1360 | ubiquitin fold modifier 1 | UFM1 | 1.0 | 1.1 | 0.5 | 0.7 | 0.3 | No |
| 1361 | unc-119 lipid binding chaperone B | UNC119B | -0.8 | -1.1 | -1.3 | -1.2 | -1.1 | No |
| 1362 | uroplakin 1A | UPK1A | -1.2 | -0.6 | -1.1 | -1.4 | -1.3 | No |
| 1363 | ubiquinol-cytochrome c reductase, Rieske iron-sulfur polypeptide 1 | UQCRFS1 | 0.9 | 1.0 | 0.5 | 0.6 | 0.3 | No |
| 1364 | ubiquinol-cytochrome c reductase hinge protein | UQCRH | 1.3 | 1.6 | 0.6 | 0.8 | 0.5 | No |
| 1365 | ubiquitin specific peptidase 14 | USP14 | 1.8 | 1.7 | 1.4 | 1.3 | 1.1 | No |
| 1366 | ubiquitin specific peptidase 39 | USP39 | 1.1 | 1.4 | 0.7 | 0.9 | 0.6 | No |
| 1367 | UTP11, small subunit processome component | UTP11 | 1.3 | 1.0 | 0.7 | 0.5 | 0.4 | No |
| 1368 | vesicle associated membrane protein 8 | VAMP8 | -1.0 | -0.9 | -1.1 | -0.8 | -0.6 | No |
| 1369 | von Hippel-Lindau tumor suppressor | VHL | -1.5 | -1.3 | -1.4 | -0.5 | -0.6 | No |
| 1370 | vimentin | VIM | -1.0 | -1.7 | -1.4 | -3.2 | -2.7 | No |
| 1371 | Wiskott-Aldrich syndrome | WAS | -0.9 | -1.1 | -1.1 | -1.2 | -1.0 | No |
| 1372 | Wiskott-Aldrich syndrome like | WASL | -1.0 | -0.8 | -0.7 | -0.2 | -0.8 | No |
| 1373 | WD repeat domain 12 | WDR12 | 1.5 | 1.1 | 0.9 | 0.9 | 0.2 | No |
| 1374 | WD repeat domain 18 | WDR18 | 1.9 | 1.7 | 1.2 | 1.6 | 0.9 | No |
| 1375 | WD repeat domain 5 | WDR5 | 1.0 | 1.2 | 0.8 | 0.9 | 0.5 | No |
| 1376 | WD repeat domain 55 | WDR55 | 1.2 | 1.0 | 0.5 | 0.2 | -0.3 | No |
| 1377 | WD repeat domain 74 | WDR74 | 1.3 | 0.7 | 0.6 | 0.5 | 0.0 | No |
| 1378 | WD repeat domain 83 | WDR83 | 0.6 | 0.8 | 0.8 | 1.4 | 1.0 | No |
| 1379 | WDYHV motif containing 1 | WDYHV1 | -0.7 | -1.0 | -0.2 | 0.2 | -0.6 | No |
| 1380 | WNT1 inducible signaling pathway protein 2 | WISP2 | -1.2 | -1.1 | -0.6 | -2.0 | -6.6 | No |
| 1381 | Werner helicase interacting protein 1 | WRNIP1 | 0.4 | 0.8 | 0.8 | 1.0 | 0.4 | No |
| 1382 | WD repeat and SOCS box containing 1 | WSB1 | -1.4 | -0.9 | -1.2 | -1.4 | -1.3 | No |
| 1383 | Xrcc1 N-terminal domain containing 1 | XNDC1 | 1.3 | 2.1 | 1.6 | 0.8 | -1.3 | No |
| 1384 | X-ray repair cross complementing 1 | XRCC1 | 0.7 | 1.0 | 0.7 | 1.0 | 0.5 | No |
| 1385 | X-ray repair cross complementing 4 | XRCC4 | 1.1 | 1.6 | 0.6 | 0.7 | -0.7 | No |
| 1386 | YY1 associated factor 2 | YAF2 | 1.2 | 0.8 | 0.7 | 0.4 | -0.4 | No |
| 1387 | tyrosyl-tRNA synthetase | YARS | 1.9 | 1.8 | 1.0 | 0.8 | 0.8 | No |
| 1388 | YdjC chitooligosaccharide deacetylase homolog | YDJC | 1.7 | 0.9 | 0.9 | 1.5 | 1.2 | No |
| 1389 | YEATS domain containing 4 | YEATS4 | 1.1 | 1.2 | 1.0 | 1.0 | 0.6 | No |
| 1390 | tyrosine 3-monooxygenase/tryptophan 5-monooxygenase activation protein epsilon | YWHAE | 0.6 | 0.9 | 0.4 | 0.5 | 0.4 | No |
| 1391 | tyrosine 3-monooxygenase/tryptophan 5-monooxygenase activation protein theta | YWHAQ | 1.6 | 2.4 | 0.9 | 0.8 | 0.2 | No |
| 1392 | YY1 transcription factor | YY1 | -0.4 | -0.7 | -0.5 | -0.4 | -10.7 | No |
| 1393 | zinc finger and BTB domain containing 16 | ZBTB16 | -3.6 | -3.4 | -3.1 | -3.0 | -4.8 | No |
| 1394 | zinc finger and BTB domain containing 2 | ZBTB2 | -0.7 | -1.5 | -1.2 | -1.8 | -2.5 | No |
| 1395 | zinc finger C3HC-type containing 1 | ZC3HC1 | 0.7 | 1.1 | 0.8 | 0.7 | 0.6 | No |
| 1396 | zinc finger C4H2-type containing | ZC4H2 | 0.2 | 0.4 | 1.0 | 1.7 | 1.2 | No |
| 1397 | zinc finger DHHC-type containing 2 | ZDHHC2 | -1.2 | -1.7 | -1.8 | -1.8 | -0.5 | No |
| 1398 | zinc finger AN1-type containing 1 | ZFAND1 | 1.0 | 1.2 | 0.8 | 0.9 | 0.6 | No |
| 1399 | zinc finger protein 143 | ZFP143 | -1.3 | -0.9 | -1.2 | -1.0 | -0.9 | No |
| 1400 | zinc finger protein 330 | ZFP330 | 0.9 | 0.9 | 0.5 | 0.5 | 0.2 | No |
| 1401 | zinc finger protein 346 | ZFP346 | 0.8 | 0.9 | 0.5 | 0.7 | 0.1 | No |
| 1402 | zinc finger protein 503 | ZFP503 | -0.8 | -0.6 | -0.2 | -0.3 | -0.7 | No |
| 1403 | zinc finger protein 592 | ZFP592 | -0.5 | -0.4 | -0.6 | -0.5 | -0.5 | No |
| 1404 | zinc finger FYVE-type containing 21 | ZFYVE21 | -1.1 | -0.9 | -1.4 | -1.4 | -1.3 | No |
| 1405 | zona pellucida glycoprotein 3 | ZP3 | 1.5 | 0.7 | 1.7 | 1.5 | 1.0 | No |
| 1406 | zinc finger RANBP2-type containing 1 | ZRANB1 | -1.2 | -1.3 | -0.9 | -0.7 | -1.0 | No |
| 1407 | zw10 kinetochore protein | ZW10 | 0.6 | 1.0 | 0.8 | 0.9 | 0.5 | No |
| 1408 | zwilch kinetochore protein | ZWILCH | 1.6 | 2.8 | 2.0 | 2.4 | 1.8 | No |
