## Supplementary Table No. 2 for "Understanding the complexity of Epimorphic Regeneration in zebrafish: A Transcriptomic and Proteomic approach"

Supplementary Table 2: List of Proteins differentially expressed based on Proteomic analysis.

| **S. No** | **Accession** | **Protein Name** | **Symbol** | **12hpa** | **1dpa** | **2dpa** | **3dpa** | **7dpa** | **ΣCoverage** | **Σ# Peptides** | **Σ# PSMs** |
| --- | --- | --- | --- | --- | --- | --- | --- | --- | --- | --- | --- |
| 1 | 40538786 | septin-9 | sept9 | 3.9 | -1.6 | -0.1 | 0.1 | -0.1 | 6.51 | 3 | 4 |
| 2 | 292623901 | si:ch211-160j6.2 | si:ch211-160j6.2 | 0.5 | 0.8 | -0.1 | -2.3 | 4.2 | 6.95 | 8 | 11 |
| 3 | 28278387 | 4-aminobutyrate aminotransferase | ABAT | 0.1 | -0.5 | 0.9 | 1.1 | 1.0 | 11.27 | 3 | 3 |
| 4 | 323510624 | ATP-binding cassette, sub-family A (ABC1), member 1B | ABCA1 | 1.0 | 1.2 | 0.4 | 0.7 | 0.2 | 2.51 | 6 | 12 |
| 5 | 62955065 | ATP-binding cassette sub-family B member 8, mitochondrial | ABCB8 | 0.6 | 1.1 | -0.2 | 0.1 | -0.5 | 13.17 | 5 | 12 |
| 6 | 148540042 | apoptotic chromatin condensation inducer 1a | acin1a | -0.1 | 3.0 | -0.8 | 0.5 | 0.4 | 3.83 | 2 | 12 |
| 7 | 54400372 | peroxisomal acyl-coenzyme A oxidase 1 | ACOX1 | 0.8 | 2.7 | -0.6 | 0.0 | -0.2 | 5.45 | 2 | 9 |
| 8 | 321400067 | aarF domain containing kinase 5 | ADCK5 | 0.8 | 0.6 | 0.6 | 1.1 | 0.6 | 2.76 | 2 | 22 |
| 9 | 326678705 | adenylate cyclase type 9-like | ADCY9 | 0.1 | -1.2 | -0.4 | -0.2 | -0.2 | 5.17 | 3 | 5 |
| 10 | 121583960 | arf-GAP with GTPase, ANK repeat and PH domain-containing protein 2 | AGAP2 | 1.7 | 1.7 | 0.3 | 0.6 | 0.2 | 4.84 | 5 | 6 |
| 11 | 323690136 | A-kinase anchor protein 2 | AKAP2 | 1.8 | 2.8 | -0.5 | -0.2 | 0.0 | 7.64 | 4 | 4 |
| 12 | 326676339 | a-kinase anchor protein 9 | Akap9 | 0.2 | 0.0 | 0.7 | 1.3 | 1.1 | 6.68 | 9 | 18 |
| 13 | 292624810 | adenylate kinase domain-containing protein 1 | AKD1 | 0.9 | 1.5 | 0.0 | 0.6 | 0.5 | 3.93 | 5 | 8 |
| 14 | 326670523 | collagen alpha-2(IV) chain, partial | alpha 2(IV) | 2.0 | 2.6 | 0.4 | 0.5 | -1.0 | 7.59 | 8 | 12 |
| 15 | 326677775 | ankyrin-1-like | Ank1 | -0.7 | 0.2 | 0.5 | 1.0 | 0.4 | 3.18 | 5 | 5 |
| 16 | 68404160 | ankyrin repeat domain-containing protein 33B | ANKRD33B | 0.5 | 1.0 | -0.7 | -0.4 | 0.4 | 5.88 | 2 | 5 |
| 17 | 334683133 | Annexin A6 | ANXA6 | 0.4 | 0.2 | 0.5 | 0.8 | 0.6 | 14.98 | 5 | 6 |
| 18 | 147902822 | adaptor-related protein complex 1, sigma-like | AP1S1 | -1.2 | -0.7 | -0.2 | 0.4 | 0.4 | 4.97 | 2 | 20 |
| 19 | 326675669 | apolipoprotein B-100 | APOB | 1.1 | 1.8 | -0.8 | 0.0 | -0.2 | 4.76 | 13 | 19 |
| 20 | 319738605 | intron-binding protein aquarius | AQR | -0.3 | 1.6 | 0.1 | 0.1 | 0.2 | 7.87 | 8 | 9 |
| 21 | 326666129 | rho GTPase-activating protein 23-like | ARHGAP23 | 0.8 | 1.5 | -0.6 | -0.4 | -0.7 | 8.03 | 8 | 11 |
| 22 | 189534278 | rho GTPase-activating protein 24 | ARHGAP24 | 1.4 | 1.1 | 0.5 | 0.9 | 0.2 | 3.99 | 3 | 6 |
| 23 | 131887346 | AT-rich interactive domain-containing protein 3B | arid3b | 1.8 | 2.0 | 0.9 | 1.1 | 1.1 | 9.12 | 3 | 3 |
| 24 | 27545261 | V-type proton ATPase subunit E 1 | ATP6V1E1 | 1.2 | 1.1 | -0.4 | -1.7 | -0.6 | 16.81 | 3 | 4 |
| 25 | 292610566 | serine/threonine-protein kinase ATR | atr | 0.7 | -0.2 | -0.2 | 1.2 | 1.8 | 3.18 | 6 | 9 |
| 26 | 319655732 | transcriptional regulator ATRX | ATRX | 1.4 | 2.6 | -0.1 | 0.4 | 0.4 | 1.79 | 4 | 8 |
| 27 | 94536611 | advillin | AVIL | 0.0 | -0.6 | 1.1 | 1.3 | 0.6 | 7.15 | 4 | 13 |
| 28 | 326672544 | 5-azacytidine-induced protein 1-like | AZI1 | 0.8 | 1.0 | -0.1 | -0.1 | 0.3 | 10.26 | 9 | 12 |
| 29 | 76253873 | UDP-GlcNAc:betaGal beta-1,3-N-acetylglucosaminyltransferase-like protein 1 | B3GNTL1 | -0.3 | 0.6 | -0.6 | -0.1 | -0.1 | 17.16 | 3 | 6 |
| 30 | 189521621 | n-acetyl-beta-glucosaminyl-glycoprotein 4-beta-N-acetylgalactosaminyltransferase 1-like | B4galnt4 | -0.8 | 2.4 | 0.2 | 0.8 | 0.4 | 8.41 | 4 | 5 |
| 31 | 317374851 | RecName: Full=Large proline-rich protein BAG6; AltName: Full=BCL2-associated athanogene 6; AltName: Full=HLA-B-associated transcript 3 | Bag6 | 0.5 | 1.1 | -1.1 | 0.4 | -0.1 | 3.88 | 3 | 8 |
| 32 | 326676328 | brain-specific angiogenesis inhibitor 1 | BAIAP2L1 | 6.6 | 6.0 | -0.5 | -0.2 | -0.3 | 8.67 | 7 | 8 |
| 33 | 41055604 | HLA-B associated transcript 1 | BAT1 | 1.8 | 1.6 | 0.0 | 0.5 | 0.7 | 10.11 | 4 | 10 |
| 34 | 326670845 | bromodomain adjacent to zinc finger domain protein 2B-like | BAZ2B | 0.8 | 1.2 | 2.4 | -0.2 | 0.6 | 4.38 | 3 | 3 |
| 35 | 190337626 | Bcl9l protein | BCL9L | -0.1 | 0.1 | 0.0 | 0.3 | -4.7 | 6.40 | 4 | 5 |
| 36 | 125841732 | baculoviral IAP repeat-containing protein 6 | BIRC6 | 2.0 | -0.2 | -0.4 | -0.3 | -0.7 | 5.13 | 10 | 10 |
| 37 | 34365524 | TPA_exp: replicase/helicase/endonuclease | BK001160.1 | -1.1 | 2.4 | 0.0 | -0.1 | 0.3 | 2.69 | 5 | 6 |
| 38 | 189527306 | Bloom syndrome protein homolog, partial | Blm | 0.3 | 0.4 | -0.2 | -0.3 | 0.3 | 4.20 | 4 | 5 |
| 39 | 326669898 | bromodomain and PHD finger-containing protein 3 | BRPF3 | -3.4 | -3.3 | -0.1 | 0.4 | 0.5 | 2.32 | 3 | 18 |
| 40 | 326670398 | protein bassoon-like | BSN | -0.4 | 0.1 | 0.3 | 0.4 | -0.1 | 6.39 | 16 | 17 |
| 41 | 229485221 | RecName: Full=Ankyrin repeat and BTB/POZ domain-containing protein BTBD11-A; AltName: Full=BTB/POZ domain-containing protein 11-A | btbd11a | -0.6 | 1.6 | -0.2 | 0.1 | 0.1 | 2.94 | 4 | 5 |
| 42 | 118150594 | uncharacterized protein LOC777745 | C12orf4 | 0.8 | 0.1 | 0.4 | 1.1 | 0.1 | 7.44 | 3 | 5 |
| 43 | 224747096 | complement C1q subcomponent subunit A | C1QA | 0.4 | 6.6 | -0.6 | -0.5 | -0.9 | 36.44 | 4 | 6 |
| 44 | 323422944 | complement C3-H1-like | C3 | 1.9 | 2.3 | 3.5 | 3.4 | 3.2 | 3.90 | 4 | 17 |
| 45 | 326670920 | uncharacterized protein C5orf42-like, partial | C5orf42 | 0.3 | 0.1 | -1.4 | 0.5 | -1.5 | 2.60 | 4 | 24 |
| 46 | 292621938 | uncharacterized protein C6orf174 homolog | C6orf174 | 1.6 | 1.5 | -2.3 | 0.7 | -1.0 | 10.77 | 5 | 11 |
| 47 | 71679707 | LOC570832 protein, partial | C7 | 0.7 | 1.2 | 0.3 | 0.6 | -0.1 | 12.30 | 5 | 9 |
| 48 | 181344480 | aminopeptidase O | C9orf3 | 1.0 | 1.1 | -0.6 | -0.1 | 0.0 | 2.80 | 2 | 24 |
| 49 | 292626858 | CDK5 and ABL1 enzyme substrate 2 | CABLES2 | -0.2 | 0.0 | 0.4 | 1.2 | 0.9 | 24.95 | 4 | 22 |
| 50 | 292625337 | voltage-dependent P/Q-type calcium channel subunit alpha-1A | CACNA1A | -0.4 | 1.2 | 0.4 | 0.9 | -0.1 | 5.74 | 9 | 40 |
| 51 | 160774064 | LOC100001129 protein | CADPS | 3.7 | 4.1 | 0.3 | 0.2 | 1.0 | 9.75 | 5 | 7 |
| 52 | 159155216 | Calcium/calmodulin-dependent protein kinase (CaM kinase) II delta 2 | CAMK2D | 4.7 | 5.3 | 0.0 | 0.7 | 0.4 | 7.91 | 2 | 3 |
| 53 | 189533822 | calpain-5-like | CAPN5 | 1.1 | 0.5 | 0.2 | 0.7 | -0.1 | 4.05 | 3 | 35 |
| 54 | 189514414 | caskin-1-like | CASKIN1 | -0.5 | 2.5 | -0.8 | -0.4 | -1.8 | 4.11 | 4 | 34 |
| 55 | 125829566 | cas scaffolding protein family member 4-like | cass4 | 1.2 | -0.3 | -0.8 | -0.3 | -0.6 | 5.94 | 3 | 4 |
| 56 | 190339564 | Zgc:194249 protein | cast | 0.1 | -0.1 | 1.1 | 1.7 | 0.5 | 7.46 | 3 | 3 |
| 57 | 326679199 | zinc finger protein castor homolog 1 | CASZ1 | -0.1 | 0.0 | -1.3 | -1.4 | -1.7 | 2.98 | 4 | 6 |
| 58 | 162287275 | cystathionine-beta-synthase a | cbsa | 0.6 | 1.0 | -1.0 | 0.6 | -0.2 | 8.06 | 4 | 18 |
| 59 | 302318876 | coiled-coil domain containing 120 | CCDC120 | 3.0 | 3.3 | -1.3 | -0.1 | -0.1 | 11.22 | 5 | 5 |
| 60 | 131889584 | coiled-coil domain-containing protein 146 | CCDC146 | -0.6 | -1.0 | 0.8 | 1.1 | 0.7 | 8.60 | 4 | 5 |
| 61 | 116004549 | coiled-coil domain-containing protein 61 | CCDC61 | 2.8 | 0.5 | 0.1 | 0.6 | 0.5 | 10.37 | 4 | 5 |
| 62 | 189531040 | protein Daple | CCDC88C | 0.2 | 1.3 | -1.4 | -0.1 | -0.3 | 8.61 | 11 | 13 |
| 63 | 375298703 | cyclin-T1 | CCNT1 | 0.1 | -1.5 | -0.7 | 0.1 | -0.5 | 6.82 | 3 | 35 |
| 64 | 40807135 | Cct4 protein | CCT4 | 1.4 | 2.5 | -0.4 | 0.1 | 0.1 | 6.94 | 2 | 4 |
| 65 | 146149105 | CD2-associated protein | CD2AP | 1.1 | 2.0 | 0.4 | 1.3 | 0.1 | 5.63 | 2 | 4 |
| 66 | 292622519 | neural-cadherin-like | CDH2 | 1.4 | 1.4 | 0.1 | -0.5 | 0.2 | 5.28 | 8 | 9 |
| 67 | 326666670 | cadherin EGF LAG seven-pass G-type receptor 1 | Celsr1 | 1.5 | 1.6 | -0.8 | -0.3 | 0.1 | 2.50 | 7 | 15 |
| 68 | 123704351 | cadherin EGF LAG seven-pass G-type receptor 2 precursor | celsr2 | 1.4 | 0.8 | 0.2 | 0.4 | -0.3 | 3.71 | 8 | 10 |
| 69 | 256000751 | histone H3-like centromeric protein A | cenpa | 1.0 | 1.8 | -0.3 | -0.2 | 0.2 | 8.97 | 2 | 5 |
| 70 | 115313145 | LOC563066 protein | CEP131 | -0.4 | 0.2 | 2.8 | -0.9 | 2.3 | 4.21 | 2 | 17 |
| 71 | 317008586 | centrosomal protein of 135 kDa | CEP135 | 0.2 | 1.0 | 0.1 | 0.1 | 0.5 | 9.54 | 6 | 11 |
| 72 | 189537675 | centrosomal protein of 152 kDa-like | Cep152 | 0.0 | -0.3 | 1.0 | 1.0 | 0.6 | 4.23 | 7 | 11 |
| 73 | 189529885 | cingulin-like | CGN | -0.3 | 0.4 | 1.9 | 2.1 | -1.4 | 6.26 | 5 | 19 |
| 74 | 326664806 | chloride channel protein 2 | CLCN2 | -0.1 | 1.5 | 0.1 | 0.5 | 0.2 | 3.32 | 5 | 6 |
| 75 | 118341447 | Wu:fb78c02 protein | CLCN3 | -0.5 | 0.0 | -0.1 | 0.7 | -2.6 | 2.20 | 2 | 27 |
| 76 | 189522889 | chloride transport protein 6 | CLCN6 | 0.2 | 0.0 | -0.4 | -0.2 | 0.1 | 3.36 | 2 | 7 |
| 77 | 292621733 | calsyntenin-2-like | CLSTN2 | 0.3 | 0.6 | -3.5 | -0.2 | -2.9 | 10.62 | 6 | 6 |
| 78 | 378556117 | cytidine monophosphate sialic acid synthetase 2 | CMAS | 0.5 | -0.1 | -0.5 | -1.0 | -0.3 | 9.30 | 2 | 4 |
| 79 | 41055983 | cap-specific mRNA (nucleoside-2'-O-)-methyltransferase 1 | Cmtr1 | 0.5 | 1.6 | 0.5 | 0.6 | 0.5 | 9.29 | 4 | 5 |
| 80 | 302318888 | cardiomyopathy associated 5 like | cmya5 | 0.7 | 1.1 | -0.1 | -0.1 | 0.0 | 3.01 | 3 | 3 |
| 81 | 292627548 | contactin-associated protein-like 2-like, partial | Cntnap2 | 2.5 | 3.3 | -0.5 | 0.0 | -0.5 | 5.30 | 3 | 8 |
| 82 | 326677363 | collagen alpha-1(XII) chain-like | col12a1 | 0.9 | 1.3 | -0.1 | 0.3 | 0.2 | 4.93 | 8 | 11 |
| 83 | 115635736 | collagen XV alpha 1 chain | col15a1a | -0.2 | 1.2 | 0.1 | 0.4 | -3.2 | 8.48 | 4 | 4 |
| 84 | 326674791 | collagen alpha-1(XIV) chain | COL16A1 | 1.3 | 1.5 | -0.1 | 0.6 | 0.5 | 7.72 | 8 | 12 |
| 85 | 292619906 | collagen alpha-1(IX) chain-like | COL1A1 | 0.5 | 1.5 | -0.4 | 0.1 | -0.1 | 13.31 | 4 | 6 |
| 86 | 48762667 | collagen alpha-2(I) chain precursor | COL1A2 | 3.7 | 4.0 | 0.2 | 0.7 | 0.1 | 17.31 | 16 | 28 |
| 87 | 41393113 | collagen, type I, alpha 1b precursor | col1a3 | 0.3 | 0.6 | 3.0 | 3.2 | 3.6 | 16.91 | 11 | 21 |
| 88 | 292620605 | collagen alpha-1(XXIII) chain | COL23A1 | -0.1 | 0.0 | -4.4 | -4.2 | 0.3 | 17.32 | 3 | 3 |
| 89 | 326667849 | collagen alpha-1(XXIV) chain | COL24A1 | 1.7 | 0.8 | 1.8 | -0.3 | 0.0 | 5.37 | 7 | 41 |
| 90 | 254028223 | collagen alpha-1(XXVII) chain A precursor | col27a1a | 3.4 | 2.0 | -0.1 | 0.3 | 0.4 | 14.30 | 10 | 13 |
| 91 | 225310541 | collagen type XXVIII alpha 1 b precursor | col28a1l | 1.2 | 1.8 | -0.2 | 0.1 | 0.2 | 9.93 | 5 | 8 |
| 92 | 326674345 | collagen alpha-1(IV) chain-like | col4a1 | -0.2 | 0.0 | -0.2 | 1.1 | -0.1 | 10.59 | 5 | 5 |
| 93 | 224809395 | collagen alpha-2(V) chain precursor | COL5A2 | -2.1 | -0.7 | 0.8 | 0.9 | 0.7 | 4.76 | 5 | 6 |
| 94 | 299782471 | collagen, type V, alpha 3b precursor | col5a3b | 1.3 | 2.0 | -0.5 | 0.0 | 0.0 | 10.80 | 12 | 17 |
| 95 | 125829720 | collagen alpha-1(VI) chain | COL6A1 | 1.9 | -0.2 | -0.9 | -0.3 | -0.9 | 6.20 | 6 | 10 |
| 96 | 326668779 | collagen alpha-1(VII) chain-like | COL7A1 | 1.0 | 0.2 | 0.0 | 0.6 | 0.0 | 10.79 | 14 | 20 |
| 97 | 116812896 | coproporphyrinogen-III oxidase, mitochondrial | CPOX | 1.5 | -0.2 | 0.4 | 0.5 | 0.7 | 2.67 | 2 | 33 |
| 98 | 51859073 | Carnitine acetyltransferase | cratb | -0.9 | -1.4 | 0.2 | -0.4 | -0.3 | 10.18 | 4 | 36 |
| 99 | 308248929 | hypothetical protein CRE_10162 [Caenorhabditisremanei] | CRE_10162 | -1.0 | -1.1 | -1.0 | -0.3 | -0.7 | 3.23 | 2 | 10 |
| 100 | 308259292 | hypothetical protein CRE_28233 [Caenorhabditisremanei] | CRE_28233 | 0.2 | 1.0 | 0.7 | 0.8 | 0.5 | 2.78 | 3 | 3 |
| 101 | 308257042 | CRE-PIF-1 protein [Caenorhabditisremanei] | Cre-pif-1 | 1.6 | 1.2 | 6.3 | 6.3 | 6.6 | 5.29 | 3 | 3 |
| 102 | 56090491 | crystallin, gamma M4 | crygm4 | 2.3 | 3.4 | 0.1 | 1.1 | 0.4 | 11.49 | 3 | 5 |
| 103 | 326680066 | chondroitin sulfate proteoglycan 4 | CSPG4 | 2.2 | 0.6 | -0.4 | -0.1 | -0.3 | 2.65 | 6 | 9 |
| 104 | 54400554 | cysteine and glycine-rich protein 3 | CSRP3 | 0.1 | -0.1 | -1.3 | -0.3 | 0.1 | 11.92 | 2 | 8 |
| 105 | 47086527 | cytoplasmic FMR1-interacting protein 1 homolog | CYFIP1 | 3.0 | 2.8 | -0.5 | 0.6 | -0.8 | 2.23 | 3 | 4 |
| 106 | 42476256 | cytochrome P450, subfamily XIA, polypeptide 1 | CYP11A1 | -0.5 | -1.5 | -0.3 | 0.3 | 0.1 | 15.72 | 4 | 9 |
| 107 | 119943089 | cytochrome P450-like | cyp2aa9 | -0.1 | -0.8 | 1.5 | 2.2 | 1.3 | 9.02 | 3 | 48 |
| 108 | 52219020 | cytochrome P450, family 8, subfamily B, polypeptide 3 | CYP8B1 | 1.3 | 1.3 | 0.2 | 0.9 | 0.8 | 14.06 | 2 | 3 |
| 109 | 292625250 | cytospin-A | CYTSA | -0.4 | 2.3 | -0.3 | 0.4 | 0.7 | 4.82 | 5 | 6 |
| 110 | 48391003 | dapper1 | DACT1 | 0.4 | 0.1 | -0.2 | -0.7 | 0.2 | 5.15 | 4 | 6 |
| 111 | 223029395 | DDB1- and CUL4-associated factor 6 | DCAF6 | 1.1 | 1.1 | 0.7 | 0.4 | 0.1 | 4.85 | 4 | 7 |
| 112 | 189526704 | dynactin subunit 1 | Dctn1 | -0.2 | 0.0 | 0.0 | 1.0 | 0.1 | 4.16 | 4 | 32 |
| 113 | 108742054 | LOC556764 protein | DDX17 | 0.0 | -0.1 | 0.9 | 1.2 | 2.1 | 5.39 | 2 | 12 |
| 114 | 120537661 | Vasa protein | DDX4 | 4.2 | 4.3 | -0.1 | -0.1 | 0.0 | 8.87 | 5 | 27 |
| 115 | 47086965 | peroxisomal 2,4-dienoyl-CoA reductase | DECR2 | 1.8 | 1.8 | -0.5 | 0.9 | -0.2 | 18.67 | 2 | 5 |
| 116 | 116267981 | DENN/MADD domain containing 2D | DENND2D | 0.7 | 1.8 | 0.1 | -0.1 | 0.7 | 7.97 | 2 | 2 |
| 117 | 18858539 | desmin a | desma | 1.8 | 2.8 | -0.2 | 0.3 | 0.0 | 12.47 | 6 | 13 |
| 118 | 41053698 | pre-mRNA-splicing factor ATP-dependent RNA helicase PRP16 | DHX38 | 1.1 | 0.8 | -0.1 | 0.0 | 0.3 | 6.20 | 7 | 8 |
| 119 | 114797050 | diaphanous 2 | DIAPH2 | 1.3 | 1.9 | -0.1 | -0.1 | 0.0 | 5.41 | 2 | 38 |
| 120 | 224496092 | disks large homolog 5 | DLG5 | 1.2 | 0.1 | -0.6 | -0.3 | -0.7 | 3.74 | 6 | 6 |
| 121 | 326666549 | deleted in malignant brain tumors 1 protein | DMBT1 | 0.7 | 0.1 | -0.3 | 2.9 | 2.7 | 5.54 | 6 | 9 |
| 122 | 292614668 | dynein beta chain, ciliary | dnah | 0.2 | 0.0 | -0.8 | -0.4 | -0.9 | 3.55 | 9 | 13 |
| 123 | 326671940 | dynein heavy chain 1, axonemal | DNAH1 | 1.1 | 1.0 | 2.5 | 0.8 | 0.9 | 4.35 | 9 | 9 |
| 124 | 292613807 | dynein heavy chain 10, axonemal | DNAH10 | 1.6 | 1.3 | -0.3 | -0.2 | -0.8 | 6.62 | 16 | 18 |
| 125 | 326669322 | dynein heavy chain 2, axonemal-like | DNAH2 | -0.5 | 0.1 | -0.5 | -0.6 | -0.8 | 5.06 | 13 | 15 |
| 126 | 189523512 | dynein heavy chain 7, axonemal | DNAH7 | -1.6 | -1.5 | -0.1 | 0.1 | -0.1 | 7.37 | 14 | 20 |
| 127 | 326672103 | dynein heavy chain 9, axonemal, partial | DNAH9 | -0.2 | -0.2 | 1.5 | 1.8 | 1.9 | 2.07 | 4 | 31 |
| 128 | 41055508 | dynamin-1-like protein | DNM1L | 5.8 | 6.5 | -0.3 | -0.1 | -0.4 | 8.25 | 4 | 56 |
| 129 | 326668898 | deoxynucleotidyltransferase terminal-interacting protein 1-like, partial | DNTTIP1 | 0.4 | -0.9 | -1.2 | -0.6 | -0.9 | 13.64 | 2 | 17 |
| 130 | 158517915 | dedicator of cytokinesis protein 1 | DOCK1 | -0.2 | 1.9 | 0.2 | 0.7 | 0.4 | 10.72 | 8 | 10 |
| 131 | 326673981 | dedicator of cytokinesis protein 10-like | DOCK10 | -0.1 | 6.6 | -0.6 | -0.1 | 0.3 | 5.31 | 8 | 10 |
| 132 | 292615010 | dedicator of cytokinesis protein 7 | Dock7 | -0.3 | 1.7 | -0.5 | 1.0 | -0.2 | 3.82 | 6 | 7 |
| 133 | 326670154 | dedicator of cytokinesis protein 8 | DOCK8 | 0.0 | -1.0 | -0.1 | 0.3 | -0.4 | 4.30 | 6 | 7 |
| 134 | 292616778 | cAMP-specific 3',5'-cyclic phosphodiesterase 4D | DPDE3 | 0.5 | -0.5 | -0.5 | -0.1 | -6.1 | 8.32 | 3 | 4 |
| 135 | 169146773 | Similar to human D4, zinc and double PHD fingers family 1 (DPF1) | DPF1 | 0.5 | 1.4 | 0.0 | -0.8 | 0.6 | 14.62 | 3 | 3 |
| 136 | 66392186 | dihydropyrimidinase-related protein 5 | DPYSL5 | 0.1 | 1.0 | -0.1 | 0.1 | 0.0 | 19.86 | 4 | 14 |
| 137 | 326673807 | dystrophin-related protein 2-like | DRP2 | -0.2 | 0.2 | -1.2 | -0.8 | -1.0 | 6.53 | 4 | 4 |
| 138 | 292621329 | Down syndrome cell adhesion molecule like 1 | DSCAML1 | 0.8 | 0.3 | 0.4 | 1.3 | 0.4 | 4.67 | 5 | 23 |
| 139 | 292618623 | e3 ubiquitin-protein ligase DTX3L | DTX3L | -0.7 | -4.3 | 0.7 | 0.8 | 0.7 | 6.95 | 3 | 3 |
| 140 | 326680379 | dual oxidase 1, partial | duox | -1.0 | 0.8 | 2.2 | -0.8 | 2.7 | 5.26 | 5 | 13 |
| 141 | 45387793 | dyslexia susceptibility 1 candidate 1 | DYX1C1 | 0.8 | 0.9 | -0.2 | 1.3 | -0.4 | 10.71 | 5 | 8 |
| 142 | 326677658 | ER degradation-enhancing alpha-mannosidase-like 3-like | EDEM3 | 6.6 | 6.3 | 0.1 | 0.3 | 0.1 | 8.36 | 3 | 25 |
| 143 | 28278942 | Eukaryotic translation elongation factor 2, like | EEF2 | -1.2 | 0.0 | -0.3 | -0.1 | -0.6 | 10.14 | 7 | 9 |
| 144 | 197246961 | Eif4bb protein | EIF4B | 0.2 | 0.1 | 4.6 | 5.0 | 4.6 | 11.04 | 5 | 15 |
| 145 | 309384237 | eukaryotic translation initiation factor 4H isoform 2 | EIF4H | -0.5 | 2.5 | -0.7 | -0.3 | -0.8 | 23.05 | 6 | 9 |
| 146 | 190339262 | Eph-like kinase 1 | ek1 | 0.6 | 1.0 | 0.3 | 0.8 | 0.3 | 12.03 | 5 | 8 |
| 147 | 292625836 | ETS-related transcription factor Elf-3 | ELF3 | 1.8 | 1.1 | 0.6 | 0.4 | 0.4 | 4.80 | 3 | 20 |
| 148 | 106635740 | tropoelastin 1 | eln1 | -0.1 | 6.6 | -0.2 | 0.1 | 0.2 | 2.23 | 2 | 18 |
| 149 | 41054802 | ectonucleotidepyrophosphatase/phosphodiesterase family member 2 precursor | ENPP2 | 1.9 | 1.1 | -1.8 | 0.2 | -1.1 | 7.53 | 3 | 20 |
| 150 | 28201964 | erythrocyte membrane protein band 4.1b (elliptocytosis 1, RH-linked) | epb41b | 0.2 | -0.3 | -0.1 | -0.1 | 0.0 | 5.02 | 6 | 13 |
| 151 | 326672585 | enhancer of polycomb homolog 1 | EPC1 | -0.5 | -1.6 | -0.5 | 0.3 | 0.3 | 10.93 | 7 | 11 |
| 152 | 326670579 | ephrin type-A receptor 3, partial | EPHA3 | 0.7 | 1.6 | -0.1 | -0.1 | 0.0 | 5.20 | 3 | 4 |
| 153 | 326671085 | ephrin type-A receptor 5-like | EPHA5 | 1.3 | 1.4 | 0.5 | 1.1 | 0.3 | 8.75 | 7 | 7 |
| 154 | 326665330 | epidermal growth factor receptor substrate 15-like 1-like, partial | EPS15 | 0.7 | -0.1 | -0.1 | 0.9 | -0.1 | 17.79 | 6 | 8 |
| 155 | 41053845 | epidermal growth factor receptor kinase substrate 8 | EPS8 | 1.1 | 0.5 | -0.1 | 0.3 | -0.3 | 7.21 | 4 | 7 |
| 156 | 125833380 | DNA excision repair protein ERCC-6 | ERCC6 | 0.1 | 1.1 | -0.2 | 0.2 | 0.1 | 6.47 | 4 | 4 |
| 157 | 124481637 | Zgc:158613 protein | erich3 | -0.2 | 0.5 | 0.4 | 0.6 | 0.8 | 7.38 | 3 | 6 |
| 158 | 326668660 | espin-like protein-like | ESPNL | 1.9 | 1.6 | 0.5 | 0.5 | 0.0 | 4.19 | 4 | 4 |
| 159 | 167555226 | exonuclease 3'-5' domain-containing protein 2 | EXD2 | 1.1 | 1.9 | -0.8 | -0.4 | -1.4 | 16.86 | 5 | 6 |
| 160 | 118150550 | protein FAM160B2 | FAM160B2 | 0.4 | -0.2 | 1.2 | 1.7 | 1.2 | 3.13 | 2 | 5 |
| 161 | 125842521 | protein FAM161B-like | FAM161B | -0.2 | -0.1 | 0.7 | 1.0 | 0.8 | 8.93 | 3 | 37 |
| 162 | 125818405 | protein FAM171B-like | FAM171B | 1.5 | 0.5 | -0.2 | 0.2 | 0.3 | 6.49 | 4 | 6 |
| 163 | 292621368 | uncharacterized protein C17orf63 homolog | FAM222B | 1.8 | 1.2 | 0.1 | 0.4 | 0.4 | 4.33 | 2 | 5 |
| 164 | 292624941 | protein FAM59B-like | FAM59B | 1.8 | 1.9 | 0.2 | 0.3 | -0.2 | 10.62 | 4 | 5 |
| 165 | 113681812 | uncharacterized protein LOC566573 | FAM65A | 0.7 | 0.7 | -0.9 | 0.2 | -0.8 | 3.05 | 3 | 5 |
| 166 | 157279721 | fanconi-associated nuclease 1 | FAN1 | 2.2 | 2.8 | -0.4 | 0.1 | -0.5 | 4.55 | 4 | 13 |
| 167 | 442746617 | Putative fanconi anemia complementation group a danio rerio fanconi anemia a | fancd2 | 1.0 | -0.1 | -0.5 | 0.1 | 0.5 | 5.52 | 3 | 5 |
| 168 | 47085727 | protocadherin Fat 1 precursor | fat1 | 1.4 | 1.4 | -1.2 | 0.1 | -0.4 | 1.13 | 5 | 5 |
| 169 | 292620786 | protocadherin Fat 2 | FAT2 | -0.2 | 1.1 | -0.1 | 0.1 | -0.6 | 4.17 | 11 | 15 |
| 170 | 326671264 | protocadherin Fat 3-like, partial | FAT3 | 3.1 | 0.7 | -0.3 | 0.4 | 0.5 | 3.33 | 6 | 11 |
| 171 | 292623023 | f-box only protein 34 | FBXO34 | -0.3 | -0.6 | -0.4 | -0.2 | -0.6 | 7.16 | 3 | 3 |
| 172 | 125836388 | f-box only protein 38-like | FBXO38 | -0.1 | -1.1 | -0.6 | 0.9 | -0.4 | 9.12 | 6 | 8 |
| 173 | 292619674 | f-box/WD repeat-containing protein 10-like | FBXW10 | 0.0 | 0.6 | 0.4 | 0.9 | 1.0 | 6.22 | 3 | 13 |
| 174 | 326671800 | FYVE, RhoGEF and PH domain-containing protein 5-like | FGD5 | 0.6 | 0.9 | -1.0 | -0.8 | -1.1 | 4.82 | 7 | 8 |
| 175 | 189529697 | FH1/FH2 domain-containing protein 3 | Fhod3 | 0.9 | 1.1 | 0.3 | 0.4 | 0.1 | 5.36 | 5 | 6 |
| 176 | 62955561 | adenosine monophosphate-protein transferase FICD | FICD | -0.4 | -1.3 | 0.0 | 0.1 | 0.1 | 16.70 | 3 | 6 |
| 177 | 158254320 | Fit2 protein | FIT2 | 0.9 | 0.9 | 1.1 | 1.8 | 1.1 | 8.73 | 2 | 8 |
| 178 | 6019201 | fli1 protein | FLI1 | -0.1 | -0.1 | -1.0 | 0.0 | 0.0 | 11.84 | 3 | 3 |
| 179 | 56548636 | Flk1b | flk1b | 1.4 | 2.1 | 0.3 | 1.1 | 1.2 | 4.59 | 3 | 5 |
| 180 | 326679837 | formin-like protein 2-like | FMNL2 | 0.5 | 0.4 | 0.4 | 0.9 | 0.6 | 6.00 | 4 | 7 |
| 181 | 61651682 | fibronectin precursor | fn1 | -0.1 | 0.8 | -1.5 | 0.9 | -1.0 | 3.57 | 4 | 9 |
| 182 | 188536034 | folliculin-interacting protein 1 | FNIP1 | 1.9 | 1.5 | 0.0 | 0.1 | -0.6 | 4.64 | 5 | 6 |
| 183 | 41055369 | forkhead box protein M1 | FOXM1 | 1.0 | -0.5 | 0.1 | 0.4 | 0.1 | 11.56 | 5 | 7 |
| 184 | 212549694 | Fras-related extracellular matrix protein 1b precursor | frem1b | -0.2 | 1.5 | -0.1 | 0.4 | 0.8 | 1.02 | 2 | 4 |
| 185 | 113678387 | fibroblast growth factor receptor substrate 2 | FRS2 | 1.2 | 1.2 | -0.1 | -0.2 | -0.4 | 13.81 | 3 | 4 |
| 186 | 227908759 | protein furry homolog-like | FRYL | 1.8 | 2.1 | -0.2 | -0.1 | -0.2 | 4.90 | 9 | 11 |
| 187 | 4587298 | alpha(1,3)fucosyltransferase | ft2 | 0.7 | 1.6 | 0.1 | 0.5 | 0.4 | 7.65 | 2 | 4 |
| 188 | 292616102 | alpha-ketoglutarate-dependent dioxygenase FTO | Fto | -0.2 | -2.8 | 0.0 | 0.4 | 0.3 | 6.02 | 3 | 3 |
| 189 | 326679310 | FYVE and coiled-coil domain-containing protein 1-like | FYCO1 | 0.9 | 1.1 | 0.8 | 0.9 | 0.2 | 7.03 | 5 | 14 |
| 190 | 403399386 | Tyrosine-protein kinase fynb; AltName: Full=Proto-oncogene c-Fynb | FYN | 0.7 | 0.8 | 1.3 | 1.7 | 1.4 | 10.11 | 4 | 5 |
| 191 | 326664121 | protein G7c | g7c | 1.8 | 2.8 | -0.3 | -0.9 | 0.0 | 8.62 | 3 | 20 |
| 192 | 292611987 | glucagon receptor-like | Gcgr | 1.0 | 0.3 | -0.2 | -0.2 | 0.0 | 10.92 | 3 | 4 |
| 193 | 326486577 | Gig2-like protein DreB | Gig2 | 0.7 | 0.7 | -0.2 | 0.9 | 0.3 | 10.79 | 2 | 11 |
| 194 | 71834468 | PERQ amino acid-rich with GYF domain-containing protein 2 | GIGYF2 | 0.8 | 1.7 | -0.4 | -0.1 | -0.6 | 4.12 | 4 | 4 |
| 195 | 125833735 | GTPase IMAP family member 4-like | GIMAP4 | 2.2 | 2.1 | -1.0 | 0.0 | 0.1 | 8.14 | 3 | 7 |
| 196 | 326680477 | GTPase IMAP family member 8-like, partial | GIMAP8 | 6.2 | 6.3 | 3.7 | -1.1 | 0.3 | 2.85 | 2 | 32 |
| 197 | 165972439 | uncharacterized protein LOC569254 | gkup | 0.4 | -2.7 | 4.2 | 4.3 | -1.2 | 6.88 | 3 | 15 |
| 198 | 19881338 | glycine receptor alphaZ1L subunit | GLRA1 | 0.3 | 0.6 | -0.1 | -0.3 | 0.6 | 7.63 | 3 | 3 |
| 199 | 28502941 | Germ cell-less homolog 1 (Drosophila) | GMCL1 | 4.5 | -5.3 | 0.1 | 0.6 | 0.7 | 5.77 | 2 | 7 |
| 200 | 326666232 | GEM-interacting protein-like | GMIP | 1.1 | 2.0 | 0.0 | 0.3 | -0.2 | 6.36 | 3 | 9 |
| 201 | 49899190 | Gnl3 protein, partial | GNL3 | -1.1 | -0.9 | -0.1 | -0.1 | 0.0 | 8.08 | 4 | 4 |
| 202 | 326670214 | G-protein coupled receptor 124, partial | GPR124 | 0.8 | 1.3 | 0.6 | 1.1 | 1.0 | 3.88 | 4 | 4 |
| 203 | 190339228 | Glutamate receptor, ionotropic, delta 2 | GRID2 | -1.2 | 0.4 | 0.2 | 0.4 | -0.6 | 11.50 | 5 | 5 |
| 204 | 113677636 | delphilin | grid2ip | 1.4 | 0.7 | -0.6 | -0.2 | -0.6 | 4.06 | 3 | 4 |
| 205 | 326678808 | glutamate receptor-interacting protein 2 | Grip2 | 0.5 | 3.3 | -0.4 | -0.3 | -0.3 | 4.46 | 3 | 4 |
| 206 | 326667890 | glucocorticoid receptor DNA-binding factor 1 | Grlf1 | -0.1 | 3.3 | 1.2 | 1.2 | 0.3 | 6.45 | 5 | 5 |
| 207 | 255003839 | glutathione S-transferase C-terminal domain-containing protein | GSTCD | 1.0 | 0.9 | -0.5 | 0.0 | 0.0 | 8.31 | 3 | 18 |
| 208 | 326664592 | general transcription factor 3C polypeptide 1-like | GTF3C1 | 2.0 | 2.1 | -1.7 | -0.5 | -0.9 | 5.22 | 5 | 6 |
| 209 | 122937177 | general transcription factor 3C polypeptide 2 | GTF3C2 | -0.1 | 2.5 | 0.0 | 0.1 | -0.1 | 4.10 | 2 | 8 |
| 210 | 326676497 | interferon-induced very large GTPase 1-like | Gvin1 | 0.5 | 1.3 | -0.2 | 0.7 | 0.2 | 5.30 | 5 | 33 |
| 211 | 66392235 | glycogen [starch] synthase, liver | GYS2 | 0.9 | 0.8 | -1.1 | -0.3 | 0.1 | 5.28 | 2 | 2 |
| 212 | 189525800 | HEAT repeat-containing protein 1-like | HEATR1 | 1.0 | 2.2 | -0.6 | -0.1 | -0.6 | 5.14 | 8 | 10 |
| 213 | 50838804 | HEAT repeat-containing protein 5A | HEATR5A | -0.2 | -1.1 | -0.1 | -0.1 | 0.0 | 4.85 | 5 | 18 |
| 214 | 292619872 | HEAT repeat-containing protein 5B-like | HEATR5B | -0.1 | 0.0 | -0.1 | 0.8 | 0.7 | 6.63 | 6 | 10 |
| 215 | 224809217 | E3 ubiquitin-protein ligase HECW1 | HECW1 | 3.9 | 3.9 | 3.3 | 3.8 | 2.7 | 5.03 | 4 | 5 |
| 216 | 47086099 | probable helicase with zinc finger domain | HELZ | -0.4 | 0.1 | 0.6 | 2.0 | -0.1 | 6.94 | 10 | 12 |
| 217 | 134026300 | LOC100002738 protein | HHIPL1 | 0.0 | 0.1 | 0.1 | 0.7 | 0.6 | 3.54 | 3 | 3 |
| 218 | 326675060 | transcription factor HIVEP2-like | HIVEP2 | 0.5 | 0.3 | 1.1 | 1.3 | 1.3 | 4.76 | 4 | 5 |
| 219 | 125805885 | helicase-like transcription factor-like | HLTF | 1.2 | 1.8 | 0.3 | 0.8 | 0.5 | 9.13 | 3 | 3 |
| 220 | 326670167 | hemicentin-1-like | HMCN1 | -0.4 | 3.1 | -0.3 | 0.3 | 0.0 | 5.45 | 8 | 10 |
| 221 | 375298754 | heterogeneous nuclear ribonucleoprotein M | hnrnpm | -0.2 | 1.1 | -1.6 | 0.0 | -0.2 | 20.26 | 9 | 10 |
| 222 | 83415126 | Hermansky-Pudlak syndrome 1 protein | HPS1 | -0.4 | 0.5 | 0.1 | 0.4 | 0.0 | 7.49 | 3 | 3 |
| 223 | 302129644 | Hermansky-Pudlak syndrome 3 | HPS3 | 1.0 | -0.2 | 0.3 | 0.8 | 0.4 | 5.88 | 5 | 6 |
| 224 | 189529013 | heat shock protein 105 kDa-like | Hsph1 | 1.0 | 1.6 | -1.7 | -1.0 | -1.2 | 8.46 | 6 | 6 |
| 225 | 189536057 | e3 ubiquitin-protein ligase HUWE1 isoform 2 | HUWE1 | -0.1 | 6.6 | -1.3 | -0.3 | -0.6 | 5.60 | 13 | 16 |
| 226 | 326670834 | interferon-induced helicase C domain-containing protein 1 | Ifih1 | -0.6 | -1.5 | -0.4 | 0.5 | 0.1 | 4.73 | 4 | 4 |
| 227 | 292614269 | interferon-induced protein with tetratricopeptide repeats 5-like | IFIT5 | 1.5 | 1.3 | -2.5 | 0.7 | -2.3 | 6.21 | 2 | 30 |
| 228 | 56693322 | intraflagellar transport protein 80 homolog | IFT80 | 1.7 | 1.2 | 0.7 | 1.4 | 0.6 | 6.69 | 3 | 6 |
| 229 | 190337705 | Igf1rb protein | IGF1R | 0.9 | 1.0 | -0.5 | 0.0 | 0.5 | 5.58 | 5 | 8 |
| 230 | 62122883 | NF-kappa-B essential modulator | IKBKG | -0.5 | -0.7 | 0.9 | 2.4 | 1.0 | 9.39 | 3 | 27 |
| 231 | 71834526 | immunoglobulin-like domain-containing receptor 2 | ILDR2 | 6.6 | 6.6 | 0.2 | 0.6 | 1.1 | 7.18 | 4 | 8 |
| 232 | 28502787 | Ilf3 protein, partial | ILF3 | 1.8 | 2.7 | -0.1 | -0.2 | 0.5 | 8.66 | 3 | 4 |
| 233 | 167860129 | acetolactate synthase-like protein | ILVBL | 1.9 | 1.2 | -0.2 | -0.5 | -0.3 | 6.12 | 3 | 8 |
| 234 | 61403300 | LOC560949 protein, partial | im:7148382 | 2.6 | 2.5 | -0.2 | -0.1 | 0.4 | 7.20 | 4 | 6 |
| 235 | 326670440 | INO80 complex subunit D-like | INO80D | 0.8 | 0.3 | -1.7 | 0.0 | -1.0 | 5.33 | 3 | 3 |
| 236 | 123704786 | INO80 complex subunit D-B | ino80db | 0.5 | 0.1 | -1.7 | -0.2 | -0.8 | 7.59 | 4 | 4 |
| 237 | 127799335 | Zgc:73290 | IRF2BPL | 0.5 | -0.5 | 0.7 | 0.5 | 0.7 | 10.59 | 3 | 3 |
| 238 | 189519226 | insulin receptor substrate 1 | IRS1 | -0.5 | 0.0 | -0.5 | 0.2 | 0.0 | 8.72 | 7 | 11 |
| 239 | 55824637 | integrin alpha V subunit | ITGAV | 0.9 | 1.1 | -0.7 | -0.3 | -0.8 | 4.21 | 4 | 45 |
| 240 | 125846722 | inositol-trisphosphate 3-kinase B | ITPKB | 0.6 | -0.3 | 0.6 | 1.1 | 0.9 | 4.87 | 2 | 8 |
| 241 | 238637318 | intersectin-2 | itsn2 | 0.8 | 1.1 | 0.5 | 0.0 | 0.7 | 3.16 | 5 | 8 |
| 242 | 226237500 | jak2 | JAK2 | 0.8 | 0.3 | -0.2 | -0.4 | -0.6 | 9.25 | 6 | 6 |
| 243 | 326674945 | protein Jumonji | Jarid2 | 0.2 | 0.9 | -1.4 | -0.9 | -1.0 | 5.94 | 6 | 6 |
| 244 | 326675738 | probable JmjC domain-containing histone demethylation protein 2C-like | JMJD1C | 1.2 | -0.1 | 0.2 | 0.6 | -0.2 | 5.61 | 9 | 22 |
| 245 | 61402674 | Jph2 protein | JPH2 | -0.3 | -0.1 | -1.6 | -1.1 | -1.3 | 6.41 | 3 | 4 |
| 246 | 326674110 | uncharacterized protein K02A2.6-like | K02A2.6 | -0.4 | -1.4 | -0.3 | 0.3 | -0.2 | 6.54 | 7 | 8 |
| 247 | 326670469 | kalirin-like | Kalrn | 0.5 | 0.8 | -0.5 | -0.1 | 0.1 | 2.40 | 7 | 11 |
| 248 | 326667988 | KN motif and ankyrin repeat domain-containing protein 1 | KANK1 | 2.7 | 0.9 | -0.5 | -0.8 | 0.3 | 6.65 | 4 | 13 |
| 249 | 82658240 | katanin p60 ATPase-containing subunit A-like 2 | KATNAL2 | 1.6 | 1.7 | -0.4 | 0.2 | 0.1 | 9.07 | 4 | 4 |
| 250 | 326665837 | potassium voltage-gated channel subfamily H member 4 | KCNH4 | -1.1 | 3.6 | -0.6 | 0.0 | 0.1 | 8.58 | 5 | 5 |
| 251 | 190337148 | Potassium voltage-gated channel, subfamily H (eag-related), member 2 | KCNH6 | 0.3 | 1.2 | 0.0 | 0.3 | 0.6 | 4.72 | 5 | 6 |
| 252 | 326670721 | potassium voltage-gated channel subfamily H member 8-like | Kcnh8 | -0.2 | 1.1 | -0.1 | 0.5 | -0.1 | 7.67 | 5 | 10 |
| 253 | 71834454 | ATP-sensitive inward rectifier potassium channel 8 | KCNJ8 | -0.1 | -3.2 | 0.1 | 0.6 | -0.1 | 7.93 | 3 | 6 |
| 254 | 292625159 | potassium channel subfamily T member 1 | Kcnt1 | 2.0 | 2.6 | -0.1 | 0.5 | -0.1 | 6.32 | 5 | 5 |
| 255 | 189527733 | lysine-specific demethylase 2A | KDM2A | 0.6 | 1.2 | -0.7 | 0.0 | -1.1 | 5.51 | 6 | 10 |
| 256 | 326673609 | lysine-specific demethylase 3B | KDM3B | 3.1 | 6.6 | -0.1 | 0.8 | 0.4 | 6.17 | 6 | 34 |
| 257 | 189515732 | lysine-specific demethylase 4A | Kdm4a | 0.5 | 1.0 | 3.2 | -0.2 | 3.5 | 5.07 | 3 | 6 |
| 258 | 326675082 | dyslexia-associated protein KIAA0319-like | KIAA0319 | 1.4 | 1.1 | 0.0 | 0.5 | 0.6 | 1.96 | 2 | 10 |
| 259 | 384551704 | protein TALPID3 | KIAA0586 | 0.2 | 6.6 | 0.0 | 0.3 | -0.2 | 2.96 | 4 | 6 |
| 260 | 292629522 | UPF0258 protein KIAA1024-like | KIAA1024 | -0.5 | 1.0 | -0.2 | 0.2 | 0.2 | 6.35 | 4 | 22 |
| 261 | 139947525 | uncharacterized protein LOC563289 | KIAA1328 | 0.5 | 0.8 | -0.1 | -0.6 | 0.5 | 6.90 | 2 | 4 |
| 262 | 326664703 | uncharacterized protein KIAA1486-like | KIAA1486 | -2.4 | -0.6 | 0.1 | 0.5 | 0.8 | 7.22 | 4 | 7 |
| 263 | 189524007 | protein KIAA1967 homolog | KIAA1967 | 4.1 | 3.9 | 2.8 | 2.3 | 0.7 | 8.59 | 5 | 5 |
| 264 | 292623315 | kinase D-interacting substrate of 220 kDa | Kidins220 | 1.3 | 1.7 | -0.2 | -0.1 | -0.5 | 5.54 | 4 | 14 |
| 265 | 292622799 | kinesin-like protein KIF13A-like, partial | KIF13A | 6.0 | 6.6 | -0.6 | 0.3 | -0.3 | 1.47 | 4 | 7 |
| 266 | 292628456 | kinesin family member 15 | KIF15 | 2.6 | 2.9 | 0.0 | 1.4 | 0.4 | 5.52 | 4 | 9 |
| 267 | 326672559 | kinesin family member 19 | KIF19 | -2.5 | -2.4 | 0.3 | 0.9 | 0.7 | 6.45 | 6 | 14 |
| 268 | 326668658 | kinesin-like protein KIF1A-like | KIF1A | -0.1 | 2.8 | -0.5 | 0.3 | -0.1 | 2.69 | 5 | 6 |
| 269 | 326677113 | kinesin family member 26Aa | KIF26A | 1.1 | 2.1 | -0.5 | -0.1 | -0.6 | 8.16 | 9 | 10 |
| 270 | 62414084 | kinesin-like protein kif7 | KIF7 | 0.4 | 0.0 | 1.3 | 0.3 | 1.2 | 2.42 | 3 | 4 |
| 271 | 326679130 | kelch-like protein 17 | KLHL17 | 0.9 | 1.6 | 0.5 | 0.3 | -0.7 | 11.51 | 4 | 7 |
| 272 | 51010971 | keratin 12 | KRT12 | -0.1 | 0.4 | -1.3 | -0.5 | -0.1 | 33.48 | 16 | 35 |
| 273 | 50539834 | type I keratin E7 | KRT17 | 1.6 | 1.1 | -1.1 | 0.2 | -0.7 | 15.98 | 7 | 10 |
| 274 | 18858425 | keratin 5 | KRT5 | 0.6 | 0.6 | -1.3 | -0.6 | 0.0 | 38.53 | 27 | 94 |
| 275 | 29335502 | keratin 8 | KRT8 | 1.2 | 1.4 | -1.0 | -0.5 | -0.2 | 21.24 | 13 | 25 |
| 276 | 66472564 | serine beta-lactamase-like protein LACTB, mitochondrial | LACTB | 2.4 | 2.5 | 0.5 | 0.7 | 0.3 | 15.86 | 4 | 6 |
| 277 | 77993334 | laminin subunit alpha-1 precursor | LAMA1 | -0.3 | 0.0 | 0.4 | 0.4 | 1.1 | 3.77 | 7 | 10 |
| 278 | 27545303 | laminin subunit beta-4 precursor | lamb4 | 1.1 | 0.7 | -0.1 | 0.6 | 1.0 | 6.62 | 6 | 8 |
| 279 | 213626113 | LATS, large tumor suppressor, homolog 1 (Drosophila) | LATS1 | 1.0 | 0.0 | -0.2 | 0.5 | 0.1 | 5.62 | 6 | 6 |
| 280 | 46329565 | Lmnb1 protein, partial | LMNB1 | 0.2 | 0.8 | 0.8 | 1.3 | 1.0 | 7.16 | 3 | 3 |
| 281 | 326672945 | hypothetical protein LOC100002463 | LOC100002463 | 0.3 | 1.4 | -0.7 | 2.4 | -1.5 | 3.28 | 3 | 24 |
| 282 | 109150074 | LOC100004107 protein | LOC100004107 | 1.0 | 0.9 | 0.3 | 0.5 | 0.8 | 6.51 | 3 | 6 |
| 283 | 326666688 | hypothetical protein LOC100034534 | LOC100034534 | -0.5 | 6.6 | 1.8 | 2.2 | 1.8 | 13.50 | 4 | 56 |
| 284 | 326667766 | hypothetical protein LOC100148510 | LOC100148510 | -0.8 | 3.3 | 0.2 | 0.9 | 0.5 | 5.32 | 9 | 12 |
| 285 | 326672965 | hypothetical protein LOC100150811 | LOC100150811 | 0.5 | 0.9 | -1.0 | -0.2 | 0.3 | 8.01 | 3 | 7 |
| 286 | 292625758 | hypothetical protein LOC100151259 | LOC100151259 | -0.4 | 4.5 | -0.4 | -0.1 | -0.7 | 15.69 | 3 | 3 |
| 287 | 326671136 | hypothetical protein LOC100329499 | LOC100329499 | 0.4 | 0.2 | 0.2 | 0.7 | 0.4 | 4.07 | 9 | 10 |
| 288 | 292628600 | hypothetical protein LOC100330801 | LOC100330801 | -0.4 | -0.2 | -0.4 | -0.2 | -0.9 | 18.24 | 3 | 18 |
| 289 | 326677269 | hypothetical protein LOC100330808 | LOC100330808 | 1.0 | 1.2 | 0.9 | 0.8 | 1.0 | 5.12 | 6 | 8 |
| 290 | 292619626 | hypothetical protein LOC100330995 | LOC100330995 | 2.3 | 3.0 | -0.4 | 0.1 | -0.4 | 5.99 | 8 | 10 |
| 291 | 326669759 | hypothetical protein LOC100331748, partial | LOC100331748 | 0.7 | -2.2 | 0.3 | 0.7 | 1.1 | 5.92 | 3 | 4 |
| 292 | 292628622 | hypothetical protein LOC100332351 | LOC100332351 | 1.0 | 1.0 | -0.5 | 0.7 | -0.2 | 17.77 | 2 | 24 |
| 293 | 326665638 | hypothetical protein LOC100332545 | LOC100332545 | 1.2 | 1.3 | -0.8 | 0.0 | 0.2 | 6.35 | 6 | 6 |
| 294 | 326667359 | hypothetical protein LOC100332651 | LOC100332651 | 0.0 | 0.3 | 2.8 | 3.0 | 2.9 | 5.60 | 4 | 5 |
| 295 | 326668978 | hypothetical protein LOC100334579 | LOC100334579 | 0.4 | 1.0 | 0.2 | 0.6 | 0.4 | 6.87 | 4 | 6 |
| 296 | 292622657 | hypothetical protein LOC100334768 | LOC100334768 | -2.7 | 0.2 | 0.7 | 1.0 | 1.0 | 4.99 | 3 | 9 |
| 297 | 326667074 | hypothetical protein LOC100534841 | LOC100534841 | 1.9 | 2.1 | -0.1 | 0.2 | -0.2 | 5.87 | 7 | 7 |
| 298 | 326669436 | hypothetical protein LOC100534893 | LOC100534893 | 3.6 | 3.5 | -0.2 | 0.4 | 0.0 | 7.19 | 6 | 6 |
| 299 | 326680639 | hypothetical protein LOC100535395 | LOC100535395 | 3.6 | 2.9 | 2.9 | 3.8 | -1.0 | 10.79 | 5 | 7 |
| 300 | 326669818 | hypothetical protein LOC100536699 | LOC100536699 | 2.2 | 1.5 | 0.1 | 1.1 | 0.1 | 2.58 | 2 | 45 |
| 301 | 326674832 | hypothetical protein LOC100537160 | LOC100537160 | -0.7 | 3.1 | -0.6 | -0.4 | -0.5 | 7.28 | 7 | 11 |
| 302 | 326674198 | hypothetical protein LOC100537179 | LOC100537179 | 0.6 | 1.0 | -0.6 | 0.2 | -0.2 | 9.13 | 5 | 6 |
| 303 | 326670626 | hypothetical protein LOC100537235 | LOC100537235 | 0.1 | 0.2 | -0.4 | 1.2 | -0.7 | 6.06 | 3 | 4 |
| 304 | 326666380 | hypothetical protein LOC100537740 | LOC100537740 | -0.1 | 1.3 | 0.0 | 0.2 | 0.3 | 2.43 | 4 | 5 |
| 305 | 326666220 | hypothetical protein LOC100538074 | LOC100538074 | 0.7 | 1.1 | 0.8 | 1.5 | 0.6 | 11.16 | 4 | 4 |
| 306 | 326670235 | hypothetical protein LOC325772 | LOC325772 | 2.1 | 0.2 | 0.2 | 0.7 | 0.7 | 7.22 | 5 | 5 |
| 307 | 326680500 | hypothetical protein LOC335285 | LOC335285 | 1.6 | 1.7 | -0.3 | 0.1 | 0.3 | 7.62 | 2 | 4 |
| 308 | 326671330 | hypothetical protein LOC402835 | LOC402835 | 0.2 | 1.5 | -0.5 | -0.2 | 0.1 | 2.20 | 6 | 7 |
| 309 | 326677626 | hypothetical protein LOC497165 | LOC497165 | -0.7 | -0.9 | -0.1 | 0.2 | 0.6 | 6.71 | 11 | 11 |
| 310 | 326664001 | hypothetical protein LOC556494 | LOC556494 | 1.4 | 0.9 | -0.4 | 0.1 | -0.4 | 3.97 | 2 | 6 |
| 311 | 292627669 | hypothetical protein LOC556561 | LOC556561 | 1.1 | 2.5 | 0.3 | 0.6 | -0.8 | 6.58 | 2 | 11 |
| 312 | 326675520 | hypothetical protein LOC557451 isoform 2 | LOC557451 | 1.1 | 1.1 | 0.1 | 0.4 | 0.0 | 3.93 | 3 | 7 |
| 313 | 292616004 | hypothetical protein LOC559035 | LOC559035 | -1.1 | -0.5 | 0.7 | 0.5 | 0.7 | 3.51 | 4 | 5 |
| 314 | 292620303 | hypothetical protein LOC559067 | LOC559067 | -0.1 | 0.0 | -1.0 | 0.0 | 0.0 | 3.96 | 13 | 19 |
| 315 | 326665199 | hypothetical protein LOC559221 | LOC559221 | 0.3 | -0.3 | -1.7 | -0.6 | -0.2 | 1.72 | 4 | 4 |
| 316 | 326677062 | hypothetical protein LOC559994 | LOC559994 | -0.1 | 1.0 | -0.1 | 0.2 | 0.2 | 2.81 | 20 | 27 |
| 317 | 326665896 | hypothetical protein LOC565612 | LOC565612 | 0.5 | 2.1 | 0.1 | 0.2 | -0.7 | 3.71 | 3 | 5 |
| 318 | 326676818 | hypothetical protein LOC566074 | LOC566074 | 0.9 | 1.2 | -0.4 | 0.3 | 0.0 | 10.57 | 3 | 5 |
| 319 | 189530379 | hypothetical protein LOC566140 | LOC566140 | 2.3 | 3.0 | 0.2 | 0.2 | -0.3 | 13.58 | 4 | 32 |
| 320 | 326673057 | hypothetical protein LOC567045 | LOC567045 | 0.1 | -0.1 | 0.6 | 1.0 | 0.7 | 4.87 | 4 | 37 |
| 321 | 326678363 | hypothetical protein LOC569436 | LOC569436 | 2.4 | 1.9 | 0.2 | 0.7 | 0.0 | 4.58 | 6 | 6 |
| 322 | 326669198 | hypothetical protein LOC569661 isoform 2 | LOC569661 | 3.0 | 3.8 | 0.3 | 0.3 | 0.6 | 4.39 | 2 | 12 |
| 323 | 326673028 | hypothetical protein LOC569976 | LOC569976 | 1.3 | 2.7 | -2.1 | 0.0 | -0.7 | 2.46 | 6 | 6 |
| 324 | 326670643 | hypothetical protein LOC570215 | LOC570215 | 1.1 | 1.4 | 0.1 | 0.5 | -0.1 | 5.95 | 4 | 4 |
| 325 | 326669712 | hypothetical protein LOC570526 | LOC570526 | -0.1 | -0.1 | -0.6 | 1.1 | -0.6 | 1.69 | 2 | 9 |
| 326 | 326674476 | hypothetical protein LOC572344 | LOC572344 | 0.1 | 0.4 | -1.6 | -0.2 | -1.1 | 6.24 | 3 | 3 |
| 327 | 125827607 | hypothetical protein LOC793455 | LOC793455 | 2.7 | 3.0 | 0.1 | 0.4 | 0.2 | 13.17 | 4 | 14 |
| 328 | 326675678 | hypothetical protein LOC796278 | LOC796278 | -0.1 | 0.3 | 3.2 | 3.6 | 3.1 | 5.06 | 5 | 6 |
| 329 | 125827065 | hypothetical protein LOC796880 | LOC796880 | 0.2 | 1.5 | 0.1 | 0.4 | 0.0 | 6.12 | 9 | 10 |
| 330 | 326671288 | hypothetical protein LOC798565 | LOC798565 | -0.1 | 3.3 | 0.0 | 0.7 | 0.4 | 3.63 | 9 | 10 |
| 331 | 326667977 | hypothetical protein LOC799338 | LOC799338 | 0.0 | 0.7 | 0.1 | 0.4 | 0.3 | 5.74 | 5 | 7 |
| 332 | 326677586 | lipoxygenase homology domain-containing protein 1-like | LOXHD1 | 1.7 | 2.7 | 0.3 | 0.5 | 0.2 | 4.43 | 4 | 5 |
| 333 | 326670797 | lactase-phlorizin hydrolase | LPH | 0.0 | 0.6 | -0.4 | 0.1 | 0.3 | 3.93 | 5 | 5 |
| 334 | 326664940 | phosphatidate phosphatase LPIN2 isoform 4 | LPIN1 | 1.5 | 1.7 | 0.9 | 1.0 | 1.1 | 5.86 | 5 | 11 |
| 335 | 326664818 | lipid phosphate phosphatase-related protein type 4 | Lppr4 | 1.8 | 1.0 | 0.1 | 0.2 | 0.0 | 13.74 | 5 | 6 |
| 336 | 131889079 | low-density lipoprotein receptor-related protein 12 | LRP12 | 1.1 | 3.4 | 0.0 | 0.1 | 0.4 | 5.88 | 4 | 6 |
| 337 | 326670753 | low-density lipoprotein receptor-related protein 1B | LRP1B | 1.4 | 1.0 | 0.2 | 0.4 | -0.4 | 1.89 | 5 | 6 |
| 338 | 41152012 | alpha-2-macroglobulin receptor-associated protein | LRPAP1 | 1.2 | 0.6 | -0.9 | -0.7 | -0.5 | 7.25 | 2 | 4 |
| 339 | 217272841 | leucine-rich PPR-motif containing | LRPPRC | -0.2 | 0.0 | 0.0 | -0.1 | -0.3 | 8.29 | 5 | 8 |
| 340 | 32766407 | Leucine rich repeat containing 40 | LRRC40 | -2.5 | 4.3 | -0.9 | -3.5 | -2.8 | 8.32 | 4 | 4 |
| 341 | 224496046 | leucine rich repeat containing 58 | LRRC58 | -0.7 | -0.8 | 0.3 | 0.5 | 0.1 | 9.97 | 2 | 7 |
| 342 | 125806066 | leucine-rich repeat and coiled-coil domain-containing protein 1 | LRRCC1 | 6.0 | 6.6 | 0.6 | 1.1 | 0.9 | 3.51 | 3 | 4 |
| 343 | 326669070 | leukotriene B4 receptor 1-like | LTB4R | 1.6 | 2.4 | 0.0 | 0.0 | -0.2 | 14.62 | 4 | 4 |
| 344 | 326671671 | lymphocyte antigen 75, partial | LY75 | -1.2 | -1.1 | 0.8 | 0.9 | 0.4 | 4.66 | 3 | 4 |
| 345 | 292628067 | alpha-mannosidase 2x-like | MAN2A2 | 6.6 | 6.3 | -0.7 | 0.6 | 0.2 | 13.83 | 7 | 12 |
| 346 | 189527122 | mitogen-activated protein kinase kinasekinase 4 | Map3k4 | 6.1 | 6.0 | 0.9 | 0.7 | 0.9 | 3.25 | 3 | 20 |
| 347 | 125833621 | mitogen-activated protein kinase kinasekinase 9 | MAP3K9 | -0.1 | 0.7 | 0.2 | 1.3 | -0.4 | 7.23 | 4 | 4 |
| 348 | 42415535 | mitogen-activated protein kinase kinasekinasekinase 5 | MAP4K5 | 0.9 | 1.2 | -0.7 | -0.2 | 0.0 | 9.00 | 5 | 8 |
| 349 | 165972405 | MAP7 domain containing 1 | map7d1b | 1.9 | 1.7 | 0.5 | 0.9 | 0.6 | 3.43 | 3 | 3 |
| 350 | 189517647 | microtubule-associated serine/threonine-protein kinase 1-like | mast1 | -0.4 | 1.4 | 0.6 | 0.3 | -1.0 | 2.88 | 4 | 24 |
| 351 | 328447229 | matrin 3-like | Matr3 | 1.0 | 0.1 | -0.1 | -0.1 | 0.0 | 4.76 | 4 | 4 |
| 352 | 148726024 | membrane-bound transcription factor protease, site 1 | mbtps1 | 0.5 | 0.5 | 1.1 | 3.3 | 1.2 | 5.68 | 4 | 4 |
| 353 | 113678532 | methylcrotonoyl-CoA carboxylase subunit alpha, mitochondrial | MCCC1 | 2.2 | 2.5 | -0.1 | -0.1 | 0.0 | 10.13 | 4 | 13 |
| 354 | 326671273 | proto-oncogene DBL-like | MCF2 | -4.5 | -1.4 | 0.2 | 0.6 | 0.4 | 5.02 | 5 | 6 |
| 355 | 27545265 | DNA replication licensing factor MCM2 | MCM2 | 2.0 | 1.8 | 0.7 | 0.5 | 0.7 | 9.20 | 5 | 9 |
| 356 | 22347793 | DNA replication licensing factor Mcm5 | MCM5 | -0.3 | 1.9 | -0.6 | 0.1 | -0.3 | 12.15 | 4 | 4 |
| 357 | 326677081 | midasin | MDN1 | -0.1 | 0.0 | -0.1 | -0.1 | -0.3 | 5.46 | 16 | 18 |
| 358 | 346421502 | mediator of RNA polymerase II transcription subunit 12 | MED12 | 1.6 | 1.6 | 0.0 | 0.3 | -0.2 | 4.98 | 5 | 7 |
| 359 | 326671046 | mitogen-activated protein kinase kinasekinase 1 | MEKK1 | 0.6 | 1.2 | -1.1 | 0.7 | -0.6 | 7.33 | 5 | 8 |
| 360 | 311901071 | molecule interacting with CasL 1 | mical1 | 0.1 | -0.1 | -1.3 | 0.1 | -0.2 | 5.55 | 3 | 52 |
| 361 | 70887663 | myeloid leukemia factor 2 | MLF2 | -0.5 | 0.1 | -0.6 | 0.6 | -0.6 | 12.00 | 3 | 3 |
| 362 | 326676474 | histone-lysine N-methyltransferase MLL4-like | MLL4 | 6.6 | 6.6 | -0.1 | -0.2 | 0.3 | 9.39 | 6 | 8 |
| 363 | 133902336 | myeloid/lymphoid or mixed-lineage leukemia | Mllt3 | 4.9 | 5.3 | 0.2 | -0.5 | -0.6 | 2.85 | 8 | 10 |
| 364 | 126631839 | Si:dkey-189p24.5 protein | MMAB | 0.9 | 1.4 | 0.0 | 0.3 | 0.4 | 35.62 | 4 | 4 |
| 365 | 292619255 | MMS19 nucleotide excision repair protein homolog | MMS19 | 1.2 | 0.5 | -0.9 | -0.8 | -1.1 | 4.53 | 3 | 11 |
| 366 | 292617923 | melanopsin-B-like | mop | 0.3 | 0.0 | -0.9 | -0.3 | 0.0 | 7.93 | 3 | 6 |
| 367 | 116268043 | MOV10-like 1 | MOV10L1 | 0.4 | 1.3 | -0.6 | 0.1 | 0.0 | 6.15 | 4 | 5 |
| 368 | 162139066 | methylthioribose-1-phosphate isomerase | MRI1 | 3.1 | 3.2 | -0.5 | -0.4 | -1.0 | 13.31 | 3 | 3 |
| 369 | 57524611 | 28S ribosomal protein S27, mitochondrial | MRPS27 | 1.6 | -0.2 | 0.4 | 1.0 | 0.5 | 10.30 | 2 | 319 |
| 370 | 23308677 | hepatocyte growth factor-like protein precursor | MST1 | 0.0 | 1.1 | -1.2 | -0.7 | -0.8 | 9.45 | 5 | 6 |
| 371 | 125842397 | metastasis-associated protein MTA1-like | MTA1 | 3.8 | 4.5 | -0.4 | -0.5 | 0.6 | 10.41 | 6 | 7 |
| 372 | 339717169 | monofunctional C1-tetrahydrofolate synthase, mitochondrial | MTHFD1L | 0.6 | 1.5 | -0.4 | 0.3 | -0.1 | 5.62 | 6 | 9 |
| 373 | 117606281 | myotubularin-related protein 2 | MTMR2 | 0.7 | 1.7 | 0.2 | 0.7 | 0.9 | 5.81 | 2 | 22 |
| 374 | 189524213 | microtubule-associated tumor suppressor candidate 2-like | mtus2 | 0.0 | -0.9 | -0.9 | -0.7 | -1.2 | 7.30 | 5 | 8 |
| 375 | 326673528 | matrix-remodeling-associated protein 5 | MXRA5 | 6.3 | 6.3 | 0.2 | 0.7 | 0.1 | 5.39 | 4 | 4 |
| 376 | 68533603 | Mybbp1a protein, partial | mybbp1a | 1.9 | -0.1 | -0.3 | -0.1 | -0.1 | 6.26 | 3 | 4 |
| 377 | 41054699 | uncharacterized protein LOC393530 | MYBPH | -0.1 | 3.0 | 0.5 | 0.1 | -1.2 | 6.40 | 3 | 4 |
| 378 | 189519129 | myosin heavy chain, fast skeletal muscle | MYH4 | 5.8 | 6.1 | 0.2 | 0.4 | 0.4 | 5.64 | 6 | 7 |
| 379 | 326679095 | myosin-7-like | MYH7 | 0.9 | 1.2 | 0.3 | 0.3 | 0.5 | 2.01 | 3 | 4 |
| 380 | 326670548 | myosin-X | myo10 | 2.2 | 1.9 | -0.7 | -0.2 | 0.0 | 5.52 | 9 | 11 |
| 381 | 326672500 | putative myosin-XVB | MYO15B | -0.2 | 1.1 | 0.7 | 1.2 | 0.7 | 4.07 | 6 | 9 |
| 382 | 326674234 | myosin-XVIIIa | MYO18A | -0.3 | 0.2 | 0.0 | 1.0 | 0.3 | 6.18 | 10 | 10 |
| 383 | 292619518 | myosin-Id | myo1d | -0.1 | 6.6 | -0.8 | 0.5 | -1.4 | 9.44 | 4 | 43 |
| 384 | 45387587 | myosin-IIIa | MYO3A | 0.6 | 1.5 | -0.5 | 0.3 | -0.7 | 2.25 | 2 | 3 |
| 385 | 326680074 | myosin-Va | MYO5A | 2.1 | 1.8 | 0.2 | 0.7 | 0.3 | 6.77 | 7 | 7 |
| 386 | 125854492 | myosin-Vc | MYO5C | 0.8 | 0.7 | -1.4 | -0.4 | 0.1 | 2.86 | 6 | 7 |
| 387 | 190339980 | Myosin VIIa | MYO7A | 0.9 | 2.0 | -0.5 | 0.2 | 0.1 | 3.58 | 7 | 8 |
| 388 | 326669624 | myosin-IXa | MYO9A | 1.7 | 1.7 | -0.6 | -0.7 | -0.6 | 6.33 | 9 | 9 |
| 389 | 326665071 | myosin-IXb | MYO9B | 1.7 | 2.0 | -0.2 | -0.1 | 0.2 | 4.23 | 5 | 6 |
| 390 | 125823175 | nucleus accumbens-associated protein 2-like | Nacc2 | 0.7 | -0.3 | 0.5 | 1.2 | 0.8 | 7.19 | 4 | 4 |
| 391 | 326680197 | neuron navigator 2 | NAV2 | 1.3 | 1.4 | -0.5 | 0.1 | 0.0 | 4.52 | 6 | 7 |
| 392 | 113676876 | neuron navigator 3 | NAV3 | 0.1 | 1.0 | -0.1 | 0.5 | -0.1 | 3.70 | 6 | 11 |
| 393 | 49618989 | nucleolin | ncl | 0.1 | 1.0 | 0.4 | 0.9 | 1.1 | 10.79 | 8 | 10 |
| 394 | 326678951 | nuclear receptor coactivator 6, partial | NCOA6 | 0.7 | 1.4 | -0.3 | -0.1 | -0.1 | 4.66 | 7 | 10 |
| 395 | 292620555 | bifunctionalheparan sulfate N-deacetylase/N-sulfotransferase 1-like | NDST1 | -2.0 | -0.7 | 2.2 | 0.6 | 2.2 | 6.10 | 3 | 30 |
| 396 | 292609653 | bifunctionalheparan sulfate N-deacetylase/N-sulfotransferase 4 | NDST4 | 2.3 | 2.4 | 0.1 | 0.7 | 0.6 | 3.89 | 3 | 4 |
| 397 | 169146098 | novel protein similar to vertebrate nebulin (NEB) | neb | 0.9 | 3.2 | 0.5 | 0.8 | 1.0 | 2.76 | 12 | 19 |
| 398 | 66910288 | LOC566027 protein, partial | NEFL | -0.1 | 0.0 | -1.4 | -0.9 | -0.3 | 14.31 | 5 | 9 |
| 399 | 66472558 | uncharacterized protein LOC553620 | neurl1b | -1.9 | -0.4 | -0.3 | -0.1 | -0.7 | 27.43 | 3 | 16 |
| 400 | 326679293 | nuclear factor of activated T-cells, cytoplasmic 2-like | NFATC2 | 1.5 | 1.5 | -0.1 | 0.5 | 0.7 | 7.45 | 5 | 5 |
| 401 | 326673182 | nidogen-1 | NID1 | 1.4 | 0.5 | 0.8 | 1.1 | 1.2 | 6.81 | 3 | 4 |
| 402 | 319738667 | NF-kappa B repressing factor | NKRF | 0.4 | 1.1 | -0.1 | 0.1 | 0.9 | 5.71 | 3 | 6 |
| 403 | 326678365 | NACHT, LRR and PYD domains-containing protein 12 | NLRP12 | 1.3 | 2.4 | 1.1 | 1.5 | 0.8 | 3.74 | 4 | 6 |
| 404 | 326667045 | NACHT, LRR and PYD domains-containing protein 3-like isoform 1 | NLRP3 | 1.3 | 1.3 | 0.2 | 1.5 | 0.3 | 9.71 | 3 | 8 |
| 405 | 326670030 | uncharacterized protein C20orf112 homolog | nol4l | 3.4 | 2.4 | 0.3 | 0.5 | -0.1 | 13.00 | 3 | 7 |
| 406 | 375331898 | nucleolar and coiled-body phosphoprotein 1 | Nolc1 | 2.4 | 3.4 | -1.4 | -0.4 | 0.2 | 3.50 | 3 | 4 |
| 407 | 116004559 | nephrocystin-1 | NPHP1 | 1.4 | 2.2 | -0.7 | 0.1 | -0.2 | 6.60 | 3 | 62 |
| 408 | 41054207 | nuclear protein localization protein 4 homolog | NPLOC4 | 0.5 | 1.3 | 0.0 | 0.9 | 0.2 | 8.33 | 3 | 4 |
| 409 | 113678661 | atrial natriuretic peptide receptor 1 precursor | NPR1 | 3.1 | 3.7 | 0.0 | 0.2 | 0.1 | 7.50 | 3 | 9 |
| 410 | 190340030 | Neuropilin 2b | NRP2 | -0.9 | -0.7 | -0.6 | 0.2 | -0.1 | 3.48 | 3 | 3 |
| 411 | 121583798 | neurexin 2b precursor | NRXN2 | 3.2 | 3.5 | -0.4 | 0.9 | -0.4 | 7.25 | 4 | 5 |
| 412 | 292621054 | hypothetical protein LOC556086 | nsd1a | 6.5 | 6.1 | -0.3 | -0.5 | -0.4 | 4.23 | 6 | 7 |
| 413 | 326674457 | nuclear mitotic apparatus protein 1, partial | NUMA1 | -2.5 | -2.4 | 0.1 | 1.2 | -0.1 | 14.50 | 4 | 5 |
| 414 | 116487854 | Nucleoporin 107 | NUP107 | 4.5 | -5.8 | 0.8 | 1.0 | 1.0 | 5.22 | 4 | 5 |
| 415 | 66392148 | nuclear pore complex protein Nup88 | NUP88 | -0.2 | 0.0 | -1.5 | 0.3 | -0.7 | 5.00 | 3 | 23 |
| 416 | 326677656 | outer dense fiber protein 2 | Odf2 | 1.4 | 2.3 | -0.5 | -0.2 | 0.0 | 9.03 | 5 | 6 |
| 417 | 220678631 | novel protein similar to vertebrate odz, odd Oz/ten-m homolog 2 (Drosophila) (ODZ2) | odz2 | 0.2 | -0.5 | -1.2 | 0.7 | -0.4 | 5.31 | 7 | 9 |
| 418 | 254028264 | oxoglutarate (alpha-ketoglutarate) dehydrogenase (lipoamide) | OGDH | -0.1 | 3.3 | -0.4 | 0.1 | 0.4 | 6.16 | 6 | 8 |
| 419 | 190339672 | One cut domain, family member, like | onecutl | 1.9 | 2.1 | -0.5 | -0.1 | -0.6 | 11.21 | 3 | 5 |
| 420 | 292625456 | olfactory receptor 2T6 | OR2T6 | 0.3 | 1.0 | -1.1 | -0.1 | 0.0 | 17.41 | 2 | 4 |
| 421 | 41053965 | origin recognition complex subunit 1 | ORC1 | 2.4 | 2.1 | -0.7 | -0.2 | -0.5 | 7.14 | 6 | 6 |
| 422 | 34392574 | gag-like protein | ORF1 | -0.1 | 0.3 | 1.6 | 1.8 | 1.3 | 8.71 | 3 | 3 |
| 423 | 224496048 | oxysterol-binding protein 1 | OSBP | 0.9 | 1.2 | 0.0 | 0.5 | 0.1 | 12.24 | 7 | 7 |
| 424 | 113462015 | otoferlin | OTOF | 1.6 | 1.4 | 0.4 | 0.7 | 1.4 | 5.62 | 6 | 6 |
| 425 | 326673979 | otolin-1-A-like | otol1a | 1.3 | 0.2 | 0.2 | 0.4 | -0.1 | 9.34 | 3 | 4 |
| 426 | 148229357 | G-protein coupled purinergic receptor P2Y10 | P2RY10 | 0.3 | 1.3 | -0.2 | -0.2 | 0.7 | 13.47 | 3 | 17 |
| 427 | 326669372 | p2Y purinoceptor 4-like | P2ry4 | 1.7 | 1.0 | 0.3 | 0.4 | 0.2 | 7.67 | 2 | 5 |
| 428 | 38488743 | paladin | PALD1 | 0.8 | 1.1 | -0.1 | 0.3 | 0.2 | 6.05 | 3 | 4 |
| 429 | 47087313 | pantothenate kinase 3 | pank1a | -0.7 | -1.6 | 0.0 | 0.5 | 0.5 | 12.87 | 3 | 12 |
| 430 | 326671190 | poly [ADP-ribose] polymerase 14-like | Parp14 | -1.1 | 0.3 | -0.5 | 0.2 | -0.1 | 3.21 | 5 | 5 |
| 431 | 131887972 | protein PAT1 homolog 1 | PATL1 | 6.3 | 6.4 | -0.1 | 0.3 | 0.8 | 6.14 | 3 | 3 |
| 432 | 326671182 | protocadherin-1 | PCDH1 | 5.9 | 5.7 | 1.0 | 0.7 | 1.0 | 3.87 | 3 | 3 |
| 433 | 292616384 | protocadherin-7, partial | PCDH7 | 1.8 | 1.1 | 1.2 | 1.2 | -0.6 | 2.63 | 4 | 5 |
| 434 | 66773380 | protocadherin 1 gamma 9 | PCDHB3 | 1.7 | 2.6 | -0.3 | -0.1 | 0.0 | 3.15 | 2 | 6 |
| 435 | 148232784 | polycomb group RING finger protein 6 | PCGF6 | 3.3 | -0.8 | -0.7 | -0.1 | -0.4 | 12.64 | 4 | 5 |
| 436 | 125819445 | phosphorylated CTD-interacting factor 1 | PCIF1 | 6.5 | 6.6 | -1.4 | -0.2 | -1.1 | 7.12 | 3 | 16 |
| 437 | 189527118 | pecanex-like protein 1 | PCNX | -1.7 | 0.0 | -0.3 | -0.4 | -0.5 | 9.02 | 11 | 14 |
| 438 | 326671610 | cAMP-specific 3',5'-cyclic phosphodiesterase 4D-like | pde4d | 0.7 | 2.0 | -0.6 | 0.8 | -0.3 | 10.58 | 4 | 6 |
| 439 | 55742280 | rod cGMP-specific 3',5'-cyclic phosphodiesterase subunit alpha | PDE6A | 1.2 | 1.4 | 0.7 | 1.0 | 1.9 | 4.78 | 3 | 6 |
| 440 | 41152317 | cone cGMP-specific 3',5'-cyclic phosphodiesterase subunit alpha' | pde6c | 1.2 | 1.5 | 0.1 | 1.1 | 0.2 | 12.68 | 5 | 43 |
| 441 | 124430733 | sister chromatid cohesion protein PDS5 homolog A | PDS5A | 3.6 | 0.3 | -0.3 | 0.5 | 0.3 | 4.70 | 5 | 6 |
| 442 | 189530829 | PDZ domain-containing protein 8-like | PDZD8 | -0.6 | -0.4 | 1.9 | 2.0 | 1.5 | 3.93 | 2 | 3 |
| 443 | 130502509 | PDZ domain containing ring finger 4 | PDZRN4 | 1.6 | 1.0 | -0.1 | -0.1 | 0.0 | 5.58 | 3 | 5 |
| 444 | 41053700 | protein pelota homolog | PELO | 1.2 | 0.6 | 0.3 | 0.6 | 0.6 | 22.86 | 4 | 4 |
| 445 | 190337432 | Period homolog 1b (Drosophila) | per1b | 1.3 | 1.4 | 0.3 | 0.2 | 0.2 | 6.58 | 8 | 10 |
| 446 | 37962936 | period 4 | per4 | 1.7 | 1.5 | 0.1 | 0.3 | 0.1 | 6.37 | 5 | 14 |
| 447 | 41053983 | peroxisome assembly protein 26 | PEX26 | -0.7 | 0.0 | -1.1 | -0.5 | -0.8 | 6.07 | 2 | 34 |
| 448 | 292626081 | piggyBac transposable element-derived protein 4-like | PGBD4 | -0.2 | 1.3 | -0.5 | 0.0 | -0.6 | 8.33 | 3 | 10 |
| 449 | 47086263 | phosphoglucomutase-2 | PGM2 | -0.2 | 0.4 | 3.0 | 3.2 | 0.0 | 8.02 | 3 | 3 |
| 450 | 117606301 | phosphorylase kinase gamma subunit 1 | PHKG1 | 1.4 | 1.4 | -1.6 | 0.2 | -0.5 | 11.42 | 3 | 5 |
| 451 | 77993310 | phosphatidylinositol 4-kinase alpha | PI4KA | 1.3 | 0.9 | 0.3 | 0.4 | 0.4 | 3.59 | 6 | 9 |
| 452 | 326678707 | serine/threonine-protein kinase pim-2 | pim2 | -0.2 | 0.4 | -0.2 | 1.6 | -0.2 | 5.49 | 2 | 24 |
| 453 | 198281999 | plakophilin-2 | PKP2 | -0.3 | 2.0 | 0.2 | 1.0 | 0.7 | 4.29 | 2 | 4 |
| 454 | 189520147 | plakophilin-4 | PKP4 | -3.4 | -3.3 | -0.8 | -0.5 | -0.3 | 7.04 | 6 | 6 |
| 455 | 18858457 | cytosolic phospholipase A2 | PLA2G4A | 1.9 | 1.5 | 0.1 | 0.8 | 0.7 | 7.15 | 3 | 4 |
| 456 | 239582779 | 1-phosphatidylinositol-4,5-bisphosphate phosphodiesterase epsilon-1 | PLCE1 | 1.1 | 0.1 | 0.3 | 0.6 | -0.3 | 6.05 | 7 | 11 |
| 457 | 190336957 | Phospholipase C, gamma 1 | PLCG1 | 0.4 | 0.1 | 1.6 | 2.1 | 1.9 | 9.07 | 5 | 5 |
| 458 | 326676167 | 1-phosphatidylinositol-4,5-bisphosphate phosphodiesterase eta-1, partial | plch1 | -0.7 | -0.3 | 0.8 | 0.9 | 0.5 | 3.52 | 4 | 7 |
| 459 | 148225260 | mitochondrial cardiolipin hydrolase | PLD6 | 1.8 | 1.8 | -0.6 | 0.1 | -0.2 | 10.57 | 2 | 3 |
| 460 | 326674641 | plectin-like | PLEC | 0.6 | 1.5 | -0.2 | 2.4 | -0.9 | 4.68 | 12 | 16 |
| 461 | 292618909 | pleckstrin homology domain-containing family A member 6-like | Plekha6 | -0.3 | 0.0 | -0.1 | 0.8 | -0.4 | 3.96 | 5 | 7 |
| 462 | 326669379 | pleckstrin homology domain-containing family A member 7 | PLEKHA7 | -0.1 | 2.9 | 3.4 | -0.8 | 0.2 | 7.81 | 5 | 7 |
| 463 | 326678944 | pleckstrin homology domain-containing family G member 1-like | PLEKHG1 | -0.3 | -1.1 | 0.4 | 0.8 | 0.1 | 2.60 | 3 | 4 |
| 464 | 326672868 | pleckstrin homology domain-containing family H member 2 | PLEKHH2 | -0.3 | 1.2 | -0.1 | 0.4 | 1.0 | 4.24 | 6 | 9 |
| 465 | 326670952 | serine/threonine-protein kinase PLK2 | PLK2 | 1.6 | 1.0 | 0.0 | 0.5 | 0.4 | 2.95 | 2 | 7 |
| 466 | 52218876 | plexin-A4 precursor | PLXNA4 | 2.4 | -0.2 | -0.8 | -0.2 | -0.1 | 3.05 | 4 | 6 |
| 467 | 326679011 | plexin-B1 | PLXNB1 | 3.0 | 2.0 | 0.1 | -0.3 | 0.7 | 3.88 | 5 | 5 |
| 468 | 158253837 | Unknown (protein for IMAGE:7216799) | PNOC | -0.3 | -0.7 | -0.2 | 0.4 | 0.2 | 8.70 | 2 | 24 |
| 469 | 189514960 | neuropathy target esterase | PNPLA6 | 0.4 | 0.0 | -0.2 | 1.0 | 1.2 | 10.39 | 8 | 10 |
| 470 | 189519296 | centrosomal protein POC5 | poc5 | 1.3 | 1.1 | -0.3 | 0.3 | 0.0 | 18.69 | 7 | 10 |
| 471 | 49618933 | DNA polymerase alpha | POLA1 | 0.2 | 0.2 | 1.0 | 0.8 | 0.4 | 2.23 | 2 | 8 |
| 472 | 326677806 | DNA polymerase theta | POLQ | 0.8 | 1.1 | -0.3 | 0.3 | 0.0 | 2.55 | 6 | 10 |
| 473 | 292620699 | DNA-directed RNA polymerase I subunit RPA1 | POLR1A | 1.4 | 1.0 | 0.7 | 0.8 | -0.2 | 4.49 | 6 | 7 |
| 474 | 451770412 | DNA-directed RNA polymerase III subunit RPC1 | POLR3A | 1.5 | 1.2 | -1.1 | -0.1 | -0.6 | 3.17 | 3 | 11 |
| 475 | 54400416 | DNA-directed RNA polymerase III subunit RPC7-like | polr3glb | -0.1 | 3.1 | -0.6 | 0.0 | 0.0 | 9.43 | 2 | 8 |
| 476 | 326675337 | ribonucleases P/MRP protein subunit POP1-like | POP1 | 0.7 | 1.3 | -0.4 | 0.0 | -0.1 | 12.66 | 6 | 7 |
| 477 | 118150590 | periostin isoform 1 precursor | POSTN | 1.3 | 1.3 | -0.1 | 0.3 | -0.1 | 14.55 | 4 | 13 |
| 478 | 260447194 | liprin-alpha-4 | Ppfia4 | -0.3 | -1.0 | 0.6 | 0.7 | 0.3 | 6.94 | 6 | 6 |
| 479 | 326680242 | inositol hexakisphosphate and diphosphoinositol-pentakisphosphate kinase 1-like | PPIP5K1 | -0.1 | 6.6 | 0.1 | 1.4 | 0.3 | 2.90 | 3 | 3 |
| 480 | 326677931 | serine/threonine-protein phosphatase 2A 55 kDa regulatory subunit B beta isoform-like | PPP2R1A | 2.4 | 2.0 | -0.2 | 0.7 | 0.0 | 9.20 | 3 | 12 |
| 481 | 326669842 | serine/threonine-protein phosphatase 2A 55 kDa regulatory subunit B alpha isoform | PPP2R2D | -0.9 | 1.6 | -1.0 | -0.3 | -0.4 | 6.05 | 3 | 6 |
| 482 | 190338318 | Prickle-like 2 (Drosophila) | PRICKLE2 | 1.6 | 1.4 | 0.2 | 0.5 | -0.3 | 10.83 | 6 | 9 |
| 483 | 326669531 | DNA-dependent protein kinase catalytic subunit | PRKDC | 0.9 | 1.3 | 0.4 | 0.5 | -0.1 | 5.79 | 15 | 17 |
| 484 | 292619982 | cGMP-dependent protein kinase 2-like | PRKG2 | 1.7 | 2.1 | 0.4 | 0.5 | 0.1 | 7.81 | 4 | 74 |
| 485 | 292613513 | proSAP-interacting protein 1-like | ProSAPiP1 | 0.6 | 0.9 | 0.2 | 0.5 | 0.4 | 6.40 | 4 | 4 |
| 486 | 284172381 | prospero-related homeobox gene 1b | prox1b | -0.2 | 0.0 | -0.8 | 0.2 | -1.0 | 6.53 | 2 | 3 |
| 487 | 169646741 | pre-mRNA processing factor 8 | PRPF8 | -1.1 | -1.0 | 0.3 | 0.8 | 0.0 | 8.88 | 10 | 14 |
| 488 | 41055708 | 26S proteasome non-ATPase regulatory subunit 3 | PSMD3 | 0.8 | 0.4 | 0.5 | 1.1 | 0.8 | 4.97 | 2 | 3 |
| 489 | 213627611 | Prostaglandin E synthase 2-like | PTGES2 | 1.6 | 1.3 | 0.8 | 1.3 | 1.0 | 10.34 | 2 | 6 |
| 490 | 326672163 | tyrosine-protein phosphatase non-receptor type 13-like, partial | Ptpn13 | 1.7 | 0.0 | 0.3 | 0.6 | 0.4 | 9.42 | 7 | 9 |
| 491 | 55742324 | tyrosine-protein phosphatase non-receptor type 4 | PTPN4 | 0.0 | 0.0 | -2.0 | 4.5 | 3.6 | 11.07 | 5 | 5 |
| 492 | 189517688 | PWWP domain-containing protein 2A-like | PWWP2A | -0.1 | 0.5 | 0.5 | 1.4 | 0.0 | 4.61 | 5 | 5 |
| 493 | 292627974 | ras-related protein Rab-19-like | RAB19 | -3.4 | -0.4 | 0.3 | 0.5 | 0.4 | 6.22 | 2 | 3 |
| 494 | 326678519 | rabGTPase-activating protein 1 | Rabgap1 | -0.4 | 0.0 | 0.6 | 0.8 | 0.5 | 7.24 | 6 | 6 |
| 495 | 326670885 | DNA repair protein RAD50 | RAD50 | 0.6 | 0.5 | 0.0 | 0.8 | 1.2 | 3.81 | 5 | 6 |
| 496 | 326670414 | e3 SUMO-protein ligase RanBP2 | RANBP2 | -0.3 | 0.1 | -0.8 | 0.2 | -0.1 | 2.77 | 5 | 6 |
| 497 | 326663944 | rap guanine nucleotide exchange factor 2 | RAPGEF2 | 4.5 | 4.9 | -0.3 | -0.3 | 0.3 | 5.84 | 7 | 10 |
| 498 | 189523269 | rap guanine nucleotide exchange factor 4 isoform 2 | Rapgef4 | -0.5 | 0.1 | 0.4 | 0.3 | 0.9 | 5.96 | 4 | 6 |
| 499 | 190194335 | ras-associated and pleckstrin homology domains-containing protein 1 | RAPH1 | 2.2 | -0.3 | -0.2 | 0.6 | 0.6 | 1.55 | 3 | 4 |
| 500 | 326664974 | rasGTPase-activating protein nGAP | RASAL2 | 0.4 | 1.2 | 0.0 | 0.2 | -0.1 | 5.39 | 6 | 18 |
| 501 | 160333122 | rasguanyl-releasing protein 3 | RASGRP3 | 0.8 | 1.5 | -0.5 | -0.6 | -1.0 | 5.08 | 3 | 5 |
| 502 | 292615125 | arginine-glutamic acid dipeptide repeats protein | RERE | -0.2 | 0.2 | 0.3 | 0.7 | 0.8 | 3.78 | 4 | 4 |
| 503 | 2102660 | receptor tyrosine kinase | RET | 1.2 | 1.7 | 1.1 | 0.9 | 0.7 | 4.97 | 3 | 3 |
| 504 | 181330398 | DNA repair protein REV1 | REV1 | -0.5 | -0.2 | -0.7 | -1.2 | -0.6 | 10.17 | 7 | 8 |
| 505 | 326676933 | DNA polymerase zeta catalytic subunit | REV3L | -1.4 | -1.3 | -0.6 | -0.5 | -0.1 | 3.72 | 4 | 5 |
| 506 | 113682006 | repulsive guidance molecule A | RGMA | -0.6 | 0.1 | 0.0 | 0.3 | -0.4 | 17.37 | 4 | 6 |
| 507 | 326667603 | rapamycin-insensitive companion of mTOR, partial | RICTOR | 1.4 | 1.5 | -0.6 | 0.1 | 0.2 | 3.03 | 5 | 6 |
| 508 | 326670268 | RIMS-binding protein 2 | RIMBP2 | 0.7 | -0.3 | -1.3 | -0.5 | -1.0 | 4.50 | 6 | 23 |
| 509 | 326675614 | zinc finger protein Rlf | RLF | 1.2 | 1.4 | -0.2 | 0.7 | 0.7 | 3.36 | 6 | 8 |
| 510 | 371874078 | E3 ubiquitin-protein ligase BRE1A | RNF20 | 0.3 | -1.2 | 1.0 | 1.4 | 0.8 | 5.63 | 5 | 6 |
| 511 | 326666269 | RING finger protein 213-like | RNF213 | 2.3 | 3.1 | 0.0 | 0.8 | 0.2 | 4.22 | 9 | 9 |
| 512 | 375298761 | Rho-associated, coiled-coil containing protein kinase 1 | ROCK1 | 1.0 | 1.4 | 0.0 | 0.3 | -0.1 | 9.86 | 9 | 12 |
| 513 | 41054591 | 60S ribosomal protein L7-like 1 | RPL7L1 | 1.4 | 0.2 | 0.1 | 0.5 | 0.0 | 6.07 | 2 | 7 |
| 514 | 63101235 | Zgc:110579 | RRNAD1 | 1.0 | 1.9 | -0.1 | -0.1 | 0.0 | 10.12 | 3 | 5 |
| 515 | 136255554 | RNA polymerase-associated protein RTF1 homolog | RTF1 | 3.2 | 3.3 | 5.0 | 5.6 | 5.4 | 4.26 | 3 | 4 |
| 516 | 326672661 | ryanodine receptor 2-like, partial | RYR2 | 1.4 | 1.0 | -0.2 | 0.2 | -0.2 | 10.74 | 18 | 22 |
| 517 | 326677163 | ryanodine receptor 3 | RYR3 | 0.4 | 0.2 | -1.5 | -0.6 | -1.1 | 7.10 | 18 | 29 |
| 518 | 123705627 | sal-like protein 4 | SALL4 | 1.8 | 1.6 | -1.1 | -0.9 | -0.9 | 1.37 | 2 | 34 |
| 519 | 134025269 | Sb:cb54 protein | sb:cb54 | 1.2 | 0.7 | -1.1 | -0.6 | -0.8 | 13.90 | 3 | 4 |
| 520 | 315583015 | protein strawberry notch homolog 1 | SBNO1 | -0.4 | -0.2 | -1.8 | -0.9 | -0.7 | 6.42 | 5 | 6 |
| 521 | 176866331 | S phase cyclin A-associated protein in the endoplasmic reticulum | SCAPER | 1.8 | 1.6 | -0.6 | -0.2 | -0.6 | 3.34 | 5 | 5 |
| 522 | 326663860 | sodium channel and clathrin linker 1 | Sclt1 | 1.4 | 1.1 | -1.5 | 0.0 | 0.0 | 11.37 | 4 | 24 |
| 523 | 326671905 | sex comb on midleg-like protein 2-like | SCML2 | 0.3 | 0.7 | 1.1 | 0.6 | 0.9 | 8.47 | 3 | 3 |
| 524 | 113675683 | sodium channel, voltage-gated, type VIII, alpha b | SCN8A | -0.4 | -2.1 | 0.5 | 1.4 | 1.1 | 5.26 | 6 | 13 |
| 525 | 110626183 | protein scribble homolog | SCRIB | 1.2 | 0.6 | 0.4 | 0.8 | 0.4 | 4.81 | 7 | 7 |
| 526 | 41053873 | succinate dehydrogenase [ubiquinone] flavoprotein subunit, mitochondrial precursor | SDHA | 1.3 | 0.4 | -3.3 | -0.2 | 0.2 | 13.46 | 3 | 13 |
| 527 | 50540242 | translocation protein SEC63 homolog | SEC63 | 0.6 | 1.0 | 0.2 | 0.2 | 0.1 | 5.73 | 4 | 4 |
| 528 | 189441654 | Zgc:172141 protein | SECISBP2L | 2.5 | 2.8 | -0.7 | -0.5 | -0.3 | 5.81 | 2 | 8 |
| 529 | 285026514 | selenoprotein O | SELO | 3.6 | 1.1 | 0.3 | 0.7 | 0.0 | 7.95 | 2 | 12 |
| 530 | 239985641 | semaphorin-3B precursor | SEMA3B | 0.6 | 3.8 | -0.9 | -0.4 | -0.8 | 10.32 | 5 | 5 |
| 531 | 41053391 | semaphorin-6A precursor | SEMA6A | 0.8 | 0.3 | 0.2 | 0.3 | -0.1 | 4.65 | 5 | 5 |
| 532 | 292616794 | semaphorin-6B | SEMA6B | -0.1 | 0.0 | -0.5 | -0.1 | -0.5 | 14.66 | 6 | 8 |
| 533 | 326666050 | hypothetical protein LOC556535 | setd1a | 6.4 | 5.9 | 0.2 | 1.1 | 0.5 | 6.57 | 8 | 8 |
| 534 | 255069736 | phosphatidylcholine:ceramidecholinephosphotransferase 2 | SGMS2 | -0.3 | 1.8 | 0.2 | 0.7 | 0.0 | 22.79 | 3 | 11 |
| 535 | 326667644 | SH2B adapter protein 3 | SH2B3 | 3.3 | -3.3 | 0.5 | 0.3 | -0.2 | 7.62 | 3 | 16 |
| 536 | 326677967 | SH3 and PX domain-containing protein 2B | SH3PXD2B | 3.8 | 3.1 | -0.6 | -0.1 | -0.2 | 12.70 | 7 | 10 |
| 537 | 27802860 | novel protein similar to DNA polymerases | si:busm1-180o5.3 | 2.8 | 1.7 | -0.6 | 0.4 | -0.1 | 4.81 | 5 | 6 |
| 538 | 292613604 | low-density lipoprotein receptor-related protein 2 isoform 5 | si:ch211-12e13.2 | -0.1 | -0.1 | 3.5 | 3.7 | 3.1 | 2.88 | 4 | 4 |
| 539 | 94732340 | novel immune type receptor protein | si:ch211-154o6.8 | -0.1 | -3.3 | 0.5 | 1.0 | 0.7 | 28.38 | 2 | 4 |
| 540 | 292627601 | hypothetical protein LOC569837 | si:ch211-161h7.4 | -0.2 | 0.1 | -1.5 | -0.3 | -1.0 | 6.11 | 5 | 7 |
| 541 | 165972487 | uncharacterized protein LOC571755 precursor | si:ch211-170d8.2 | 0.0 | 0.4 | 0.9 | 1.4 | 0.9 | 11.45 | 2 | 2 |
| 542 | 292617529 | hypothetical protein LOC558404 | si:ch211-67e16.4 | 0.3 | 0.6 | 0.5 | 1.0 | 0.7 | 14.52 | 3 | 9 |
| 543 | 292625508 | hypothetical protein LOC100006560 | si:ch73-112l6.1 | -0.2 | 1.3 | 0.9 | 1.2 | 0.5 | 5.46 | 4 | 12 |
| 544 | 125806140 | zinc finger protein 600 | si:dkey-154p10.3 | -0.1 | 0.1 | 0.1 | 1.1 | 0.6 | 4.02 | 3 | 3 |
| 545 | 189523351 | hypothetical protein LOC564101 | si:dkey-230p4.1 | 1.9 | 2.2 | -0.9 | -0.1 | -0.1 | 3.60 | 10 | 12 |
| 546 | 94732798 | novel protein | si:dkey-27c15.3 | 0.7 | 1.4 | 0.3 | 0.4 | 0.3 | 6.33 | 4 | 19 |
| 547 | 292621401 | rho GTPase-activating protein 32 | si:dkey-287f10.3 | -1.0 | 0.7 | 0.2 | 0.6 | 0.4 | 3.52 | 6 | 8 |
| 548 | 320089554 | intermediate filament protein-like | si:dkey-33c12.3 | -0.1 | -1.1 | -1.4 | 0.9 | -1.1 | 10.31 | 3 | 4 |
| 549 | 56207720 | novel protein containing a beta-ketoacyl synthase, N-terminal domain | si:dkey-61p9.11 | 0.5 | 4.4 | -0.9 | 2.1 | -1.5 | 2.57 | 3 | 5 |
| 550 | 194353937 | uncharacterized protein LOC792544 | si:dkey-67c22.2 | -0.8 | -1.4 | -0.6 | 0.5 | -0.1 | 4.12 | 5 | 7 |
| 551 | 113678676 | paired amphipathic helix protein Sin3b | SIN3B | 0.5 | 1.1 | 0.2 | 0.7 | 0.5 | 9.53 | 12 | 14 |
| 552 | 125821751 | solute carrier family 12 member 4 | SLC12A4 | -0.2 | -0.1 | 0.9 | 1.5 | 0.9 | 4.47 | 4 | 7 |
| 553 | 182888608 | Zgc:91959 protein | SLC1A1 | -1.0 | 0.2 | -0.2 | -0.1 | 0.1 | 12.55 | 2 | 10 |
| 554 | 296455205 | excitatory amino acid transporter SLC1A8a | slc1a8a | 1.3 | 1.2 | 0.3 | -0.4 | 0.8 | 11.52 | 3 | 6 |
| 555 | 326670104 | solute carrier family 26 member 6 | Slc26a6 | -0.2 | -0.7 | 1.0 | 1.9 | 0.6 | 9.26 | 3 | 5 |
| 556 | 189523618 | sodium/myo-inositol cotransporter-like | Slc5a3 | -2.0 | 4.2 | -0.1 | -0.1 | 0.0 | 4.45 | 2 | 10 |
| 557 | 125851107 | high affinity choline transporter 1-like | SLC5A7 | 0.2 | -0.8 | 0.3 | 1.3 | -0.2 | 9.88 | 3 | 4 |
| 558 | 125995400 | SWI/SNF-related matrix-associated actin-dependent regulator of chromatin subfamily A member 5 | SMARCA5 | -0.4 | -0.6 | 0.2 | 0.5 | 0.0 | 4.67 | 6 | 12 |
| 559 | 302129641 | structural maintenance of chromosomes protein 5 | SMC5 | 0.8 | 0.4 | -0.1 | 3.1 | 2.8 | 4.94 | 5 | 7 |
| 560 | 292622980 | structural maintenance of chromosomes protein 6 | SMC6 | 1.5 | 3.0 | -0.2 | 0.0 | -0.1 | 8.23 | 4 | 5 |
| 561 | 181331982 | U5 small nuclear ribonucleoprotein 200 kDa helicase | SNRNP200 | 0.5 | -0.5 | -0.3 | 1.0 | -1.2 | 1.45 | 2 | 2 |
| 562 | 292620207 | sorbin and SH3 domain-containing protein 1 | Sorbs1 | -0.5 | 1.7 | -0.8 | -0.4 | -0.8 | 4.30 | 4 | 5 |
| 563 | 326668207 | cytospin-B | SPECC1 | 1.0 | 1.0 | -1.8 | 0.1 | -0.1 | 5.24 | 4 | 5 |
| 564 | 326670781 | striated muscle-specific serine/threonine-protein kinase | Speg | 0.1 | 0.5 | -0.7 | 0.3 | -0.1 | 5.07 | 12 | 16 |
| 565 | 326679167 | msx2-interacting protein | Spen | 0.8 | 1.4 | -0.8 | -0.3 | 0.3 | 6.48 | 12 | 13 |
| 566 | 37359664 | Shadoo protein | sprn | 1.8 | 1.9 | 2.7 | 3.7 | 3.4 | 12.88 | 2 | 5 |
| 567 | 326673173 | spectrin beta chain, brain 4-like | sptbn4 | -1.1 | -1.0 | 2.8 | 3.0 | 0.8 | 4.05 | 3 | 7 |
| 568 | 148235634 | sterol regulatory element-binding protein 2 | SREBF2 | -0.2 | 1.6 | -0.3 | 0.5 | 0.1 | 12.01 | 7 | 10 |
| 569 | 41055859 | signal recognition particle 54 kDa protein | SRP54 | 0.7 | 1.2 | -0.6 | -0.3 | -0.7 | 12.90 | 4 | 8 |
| 570 | 319918875 | serine/arginine repetitive matrix 2 | srrm2 | 0.6 | 1.1 | -0.1 | -0.1 | 0.0 | 9.59 | 11 | 11 |
| 571 | 148277652 | serrate RNA effector molecule homolog | srrt | -0.1 | 1.2 | -0.5 | 0.2 | -0.1 | 5.25 | 4 | 4 |
| 572 | 190609992 | TPA: SCO-spondin precursor | SSPO | 0.4 | 1.7 | 0.6 | 1.1 | 0.7 | 2.40 | 7 | 9 |
| 573 | 64966362 | alpha-2,3-sialyltransferase | st3gal5l | -0.1 | -0.1 | 2.2 | 1.5 | -0.4 | 13.65 | 3 | 4 |
| 574 | 284005073 | uncharacterized protein LOC797439 | st6gal2b | -1.2 | -0.5 | -0.3 | 0.2 | -0.1 | 3.53 | 2 | 4 |
| 575 | 125805891 | cohesin subunit SA-1 | STAG1 | 4.5 | 4.9 | -0.3 | -0.1 | -0.2 | 6.68 | 4 | 5 |
| 576 | 187608279 | stAR-related lipid transfer protein 8 | STARD8 | 0.5 | 1.2 | -0.7 | -0.2 | 0.2 | 7.25 | 3 | 3 |
| 577 | 326675708 | stAR-related lipid transfer protein 9-like | STARD9 | -0.1 | 1.0 | -0.5 | 0.1 | -0.2 | 2.77 | 6 | 6 |
| 578 | 326663880 | stromal interaction molecule 2-like | STIM2 | 1.1 | 1.9 | -0.9 | -0.1 | 0.0 | 5.65 | 5 | 15 |
| 579 | 41054445 | serine/threonine-protein kinase 3 | STK3 | 0.9 | 1.4 | -0.2 | 0.2 | 3.5 | 7.52 | 3 | 5 |
| 580 | 292625366 | stromelysin-3 | STMY3 | -0.1 | 1.1 | -0.5 | 0.6 | -0.3 | 14.92 | 4 | 7 |
| 581 | 315507141 | sulfotransferase family 1, cytosolic sulfotransferase 5 | SULT1C4 | -1.1 | -0.9 | 0.1 | 0.7 | 0.7 | 4.78 | 2 | 2 |
| 582 | 11527858 | transcription elongation regulator FOGGY | SUPT5H | 5.8 | 5.9 | -0.4 | 1.4 | 0.3 | 7.29 | 4 | 9 |
| 583 | 21539661 | transcription elongation factor SPT6 | SUPT6H | 1.8 | 2.2 | 0.4 | 0.5 | 0.1 | 5.56 | 7 | 9 |
| 584 | 143586162 | RecName: Full=Histone-lysine N-methyltransferase SUV420H1; AltName: Full=Suppressor of variegation 4-20 homolog 1; Short=Su(var)4-20 homolog 1; Short=Suv4-20h1 | SUV420H1 | 0.0 | -0.1 | 0.9 | 1.4 | 0.5 | 8.79 | 6 | 32 |
| 585 | 302316220 | Swap70b, partial | swap70b | 1.4 | 1.5 | 0.1 | 0.4 | 0.6 | 4.82 | 2 | 4 |
| 586 | 326670849 | protein TANC1-like | TANC1 | 0.9 | 1.5 | -0.8 | -0.1 | 0.0 | 4.87 | 6 | 7 |
| 587 | 125812260 | protein TANC2 | TANC2 | 0.8 | 1.6 | 0.2 | 0.5 | -0.3 | 4.47 | 6 | 9 |
| 588 | 326663920 | TBC1 domain family member 1 | TBC1D1 | -0.4 | -0.4 | -0.1 | -0.1 | 0.0 | 4.47 | 4 | 5 |
| 589 | 371940897 | TBC1 domain family member 16 | TBC1D16 | 0.2 | -0.1 | 0.9 | 0.7 | 1.1 | 3.21 | 2 | 4 |
| 590 | 149383922 | tudor domain containing protein 7 | tdrd7a | 1.7 | 1.5 | 0.2 | 0.3 | 0.0 | 1.48 | 2 | 6 |
| 591 | 18859469 | teneurin-3 | TENM3 | 1.9 | 2.5 | -0.2 | 0.2 | -0.2 | 1.97 | 5 | 6 |
| 592 | 326671251 | retrotransposable element Tf2 155 kDa protein type 1-like | Tf2 | 0.7 | 2.1 | 0.1 | 0.5 | 0.1 | 7.54 | 6 | 8 |
| 593 | 292627661 | transferrin receptor protein 1-like | TFRC | 4.6 | 4.2 | -0.1 | -0.1 | 0.0 | 8.30 | 3 | 3 |
| 594 | 326671554 | protein timeless homolog | Timeless | 1.1 | 1.3 | -0.7 | -0.4 | -0.6 | 6.34 | 5 | 7 |
| 595 | 320118871 | tight junction protein ZO-2 isoform 2 | TJP2 | -0.5 | -0.7 | -0.2 | 0.3 | 0.3 | 8.51 | 7 | 10 |
| 596 | 57222259 | talin-1 | TLN1 | 0.1 | -0.8 | 0.7 | 1.1 | 0.6 | 5.56 | 7 | 7 |
| 597 | 326672343 | trinucleotide repeat-containing gene 6B protein | TNRC6B | 0.0 | -0.4 | -0.8 | -0.7 | -1.0 | 2.52 | 3 | 3 |
| 598 | 292611936 | DNA topoisomerase 3-alpha | TOP3A | 1.7 | 1.4 | -0.9 | 0.2 | 0.5 | 1.90 | 2 | 2 |
| 599 | 165972373 | TNF receptor-associated protein 1 | TRAP1 | 0.5 | 1.3 | -0.1 | -0.1 | 0.0 | 11.54 | 3 | 4 |
| 600 | 326665627 | e3 ubiquitin/ISG15 ligase TRIM25-like | TRIM25 | 1.6 | 1.6 | -0.2 | 0.3 | -0.2 | 8.01 | 3 | 4 |
| 601 | 326672583 | tripartite motif-containing protein 39-like | TRIM39 | -0.5 | 2.7 | 0.9 | 1.0 | -0.1 | 8.37 | 2 | 8 |
| 602 | 292624553 | e3 ubiquitin-protein ligase TRIM9-like | TRIM67 | -0.3 | -0.3 | 0.1 | 0.7 | 0.5 | 13.41 | 5 | 9 |
| 603 | 326674684 | triple functional domain protein, partial | TRIO | -0.3 | 2.7 | 0.7 | 0.6 | 1.0 | 2.18 | 2 | 7 |
| 604 | 190194321 | transient receptor potential cation channel subfamily M member 5 | TRPM5 | 1.1 | 1.1 | 0.5 | 0.1 | 0.0 | 2.24 | 2 | 8 |
| 605 | 34330186 | transient receptor potential cation channel, subfamily N, member 1 | trpn1 | 1.0 | 0.5 | 0.8 | 0.7 | 0.6 | 6.75 | 7 | 8 |
| 606 | 112293297 | transient receptor potential cation channel subfamily V member 4 | TRPV4 | 0.8 | 1.6 | 0.1 | 0.2 | -0.4 | 6.06 | 4 | 11 |
| 607 | 56693306 | tetratricopeptide repeat protein 26 | TTC26 | 1.5 | 1.7 | -0.5 | -0.2 | -0.8 | 11.31 | 2 | 4 |
| 608 | 32451688 | Ttk protein kinase | TTK | 1.1 | 0.4 | 0.6 | 0.7 | 0.4 | 6.01 | 5 | 8 |
| 609 | 82658236 | tubulin, beta, 2 | TUBB2B | 0.7 | 1.5 | 0.0 | 0.0 | -0.4 | 11.69 | 3 | 4 |
| 610 | 55962850 | novel protein similar to vertebrate tubby like protein 4 (TULP4) | TULP4 | -0.4 | -0.2 | -1.4 | -1.4 | -1.7 | 5.58 | 4 | 5 |
| 611 | 326672994 | transposon TX1 uncharacterized 149 kDa protein-like | TX1 | 1.7 | 1.4 | 0.4 | 0.5 | 0.0 | 1.86 | 2 | 4 |
| 612 | 326666158 | non-receptor tyrosine-protein kinase TYK2-like | TYK2 | -0.7 | -0.3 | -0.5 | 0.0 | 0.0 | 9.02 | 5 | 9 |
| 613 | 326676072 | uveal autoantigen with coiled-coil domains and ankyrin repeats-like | UACA | 0.0 | 0.1 | 0.9 | 2.0 | 0.2 | 7.69 | 3 | 16 |
| 614 | 47087029 | SUMO-activating enzyme subunit 2 | UBA2 | 1.4 | 1.8 | 0.8 | 1.1 | 1.2 | 14.22 | 4 | 7 |
| 615 | 326675715 | e3 ubiquitin-protein ligase UBR1 | UBR1 | 1.3 | 1.5 | 1.3 | 1.4 | 1.3 | 1.18 | 3 | 15 |
| 616 | 256419025 | E3 ubiquitin-protein ligase UBR5 | UBR5 | 0.1 | 1.5 | -0.7 | 0.5 | 0.2 | 4.72 | 9 | 29 |
| 617 | 189523562 | UDP-glucose:glycoprotein glucosyltransferase 2 | UGGT2 | 0.7 | 0.7 | -0.3 | 1.0 | -0.3 | 5.81 | 5 | 9 |
| 618 | 294610624 | UDP glucuronosyltransferase 5 family, polypeptide A4 precursor | ugt5a4 | 0.8 | 1.2 | 0.2 | 0.2 | 0.0 | 15.62 | 3 | 13 |
| 619 | 153792789 | UHRF1-binding protein 1-like | UHRF1BP1L | 2.3 | -0.2 | -0.5 | -0.2 | 0.1 | 2.82 | 2 | 31 |
| 620 | 326674207 | serine/threonine-protein kinase ULK2-like | Ulk2 | 0.9 | 1.5 | 0.3 | 0.1 | -0.4 | 2.82 | 3 | 6 |
| 621 | 320461533 | regulator of nonsense transcripts 3A | UPF3A | 0.4 | -0.5 | 1.4 | 2.0 | 1.4 | 13.81 | 4 | 5 |
| 622 | 115529389 | ubiquinol-cytochrome c reductase complex chaperone | UQCC1 | 1.5 | 0.5 | -0.1 | 0.3 | 0.4 | 5.59 | 2 | 19 |
| 623 | 326664297 | ubiquitin carboxyl-terminal hydrolase 34 | USP34 | 1.4 | -0.2 | 0.7 | 1.3 | 0.7 | 3.03 | 6 | 21 |
| 624 | 187607866 | guanine nucleotide exchange factor VAV3 | VAV3 | 4.3 | 4.8 | 1.4 | 1.4 | -0.2 | 6.69 | 5 | 6 |
| 625 | 124481685 | Zgc:158875 protein | VCAM1 | -0.1 | 0.4 | -2.6 | -1.6 | -1.5 | 11.63 | 6 | 11 |
| 626 | 326669419 | vacuolar protein sorting-associated protein 13C | VPS13C | 1.8 | 2.0 | -0.1 | 0.3 | -0.3 | 7.63 | 13 | 16 |
| 627 | 229892338 | V-set and immunoglobulin domain-containing protein 10 precursor | VSIG10 | 1.1 | 1.3 | 3.9 | 2.7 | 0.9 | 13.42 | 3 | 8 |
| 628 | 156713467 | vitellogenin 7 precursor | vtg7 | 0.5 | -0.2 | -0.2 | 0.2 | 0.4 | 7.81 | 5 | 5 |
| 629 | 190358461 | von Willebrand factor A domain-containing protein 8 | VWA8 | 0.7 | -4.0 | -0.5 | -0.1 | -1.2 | 4.75 | 6 | 6 |
| 630 | 319738599 | WD repeat-containing protein 47-like | WDR47 | 0.7 | 0.2 | 0.1 | 0.5 | 0.4 | 12.97 | 5 | 6 |
| 631 | 292621536 | hypothetical protein LOC100333062 | WDR81 | 1.8 | 2.3 | -0.2 | -1.3 | 0.3 | 6.15 | 5 | 6 |
| 632 | 94574343 | Wee1 protein | WEE1 | 1.4 | 1.8 | -0.9 | -0.4 | -0.9 | 5.15 | 3 | 5 |
| 633 | 160333111 | protein WWC3 | WWC3 | 1.7 | 1.1 | -1.0 | -0.5 | -0.4 | 5.49 | 8 | 9 |
| 634 | 326675414 | XK-related protein 6-like | Xkr6 | -1.7 | -0.4 | -0.4 | 0.1 | 0.2 | 19.90 | 5 | 12 |
| 635 | 326663749 | DNA repair protein complementing XP-G cells homolog | Xpg | 3.0 | 2.8 | -0.1 | 0.3 | -0.1 | 4.01 | 3 | 9 |
| 636 | 139949035 | yrdC domain-containing protein, mitochondrial | YRDC | 0.7 | 1.1 | -0.1 | 0.4 | 0.4 | 17.87 | 3 | 3 |
| 637 | 130507929 | zinc finger BED domain-containing protein 4 | ZBED4 | -1.1 | -0.7 | -0.1 | -0.1 | 0.0 | 3.92 | 4 | 6 |
| 638 | 326673516 | SCAN domain-containing protein 3-like | ZBED9 | -0.2 | 0.6 | 0.4 | 0.9 | 1.1 | 4.50 | 2 | 14 |
| 639 | 89158467 | KAISO-like zinc finger protein | zbtb4 | -0.1 | 2.7 | -0.1 | 0.1 | 0.7 | 3.07 | 2 | 12 |
| 640 | 319738618 | zinc finger CCCH domain-containing protein 13 | ZC3H13 | 1.6 | -0.4 | -0.7 | 0.0 | 0.0 | 9.38 | 18 | 32 |
| 641 | 55153647 | Zc3hdc1l protein, partial | zc3hdc1l | -0.7 | 3.0 | -0.6 | -0.2 | 0.1 | 5.36 | 4 | 18 |
| 642 | 61806554 | palmitoyltransferase ZDHHC2 | ZDHHC2 | 1.3 | 1.1 | -0.4 | -0.4 | -0.4 | 18.28 | 2 | 2 |
| 643 | 326669693 | zinc finger homeobox protein 3-like | ZFHX3 | 0.8 | 1.9 | -0.5 | 0.4 | 0.1 | 2.72 | 9 | 11 |
| 644 | 168480106 | zinc finger RNA-binding protein [Mus musculus] | ZFR | 1.1 | 0.8 | 0.4 | 0.5 | 0.4 | 5.87 | 4 | 4 |
| 645 | 292617947 | zinc finger FYVE domain-containing protein 16-like | ZFYVE16 | 0.8 | 1.6 | -0.7 | -0.5 | -0.8 | 6.95 | 3 | 3 |
| 646 | 62955799 | uncharacterized protein LOC550612 | zgc:113229 | 5.7 | 6.3 | -0.1 | 0.5 | -1.0 | 6.15 | 4 | 9 |
| 647 | 66910302 | Zgc:136472 protein | zgc:136472 | -0.3 | -0.4 | -0.8 | 0.1 | -0.2 | 16.84 | 3 | 3 |
| 648 | 90093334 | transitional endoplasmic reticulum ATPase-like | zgc:136908 | 0.4 | 1.9 | -0.6 | -0.2 | -0.3 | 5.59 | 3 | 18 |
| 649 | 118150536 | uncharacterized protein LOC777712 | zgc:152816 | 1.2 | 1.2 | -0.6 | 0.2 | -0.2 | 2.19 | 2 | 9 |
| 650 | 139948785 | mediator of RNA polymerase II transcription subunit 13-like | zgc:153454 | -1.8 | 4.1 | 1.7 | 1.9 | 0.4 | 2.33 | 5 | 6 |
| 651 | 161611869 | Zgc:171422 protein | zgc:171422 | 0.2 | 0.1 | 1.3 | 0.7 | -0.5 | 7.06 | 3 | 4 |
| 652 | 167555059 | uncharacterized protein LOC564165 | zgc:172323 | 0.1 | 1.0 | -0.6 | -0.8 | -0.8 | 9.45 | 4 | 8 |
| 653 | 55925514 | uncharacterized protein LOC492329 | zgc:92107 | 0.0 | 0.5 | -1.3 | -0.5 | 0.2 | 9.93 | 6 | 39 |
| 654 | 49227592 | zic family member 6 | zic6 | 0.5 | 0.3 | 0.0 | 0.3 | 1.0 | 6.67 | 2 | 23 |
| 655 | 326667339 | zinc finger protein 271-like | ZNF271 | 1.3 | 1.5 | 0.5 | 1.1 | 0.3 | 11.46 | 4 | 7 |
| 656 | 52218888 | zinc finger protein 384 like | ZNF384 | 0.7 | -0.1 | -1.2 | -0.8 | -1.2 | 10.08 | 2 | 3 |
| 657 | 326678440 | zinc finger protein 624 | ZNF624 | -0.3 | -0.1 | -0.4 | -0.1 | -0.1 | 4.45 | 3 | 4 |
| 658 | 326677520 | zinc finger protein 658 | ZNF658 | 1.8 | 1.5 | 0.6 | 0.7 | 0.6 | 8.19 | 3 | 7 |
| 659 | 181327693 | zinc finger protein 692 | ZNF692 | 1.0 | -0.1 | -0.3 | 0.5 | -0.1 | 3.58 | 2 | 5 |
| 660 | 149773447 | uncharacterized protein LOC449909 | ZNF729 | -0.8 | -1.1 | 1.0 | 1.0 | 0.7 | 6.71 | 2 | 23 |
| 661 | 117606303 | zinc finger HIT domain-containing protein 2 | ZNHIT2 | -0.9 | 3.4 | 1.2 | 0.9 | 1.0 | 6.15 | 4 | 5 |
