## Supplementary Table No. 3 for "Understanding the complexity of Epimorphic Regeneration in zebrafish: A Transcriptomic and Proteomic approach"

Supplementary Table 1: List of Genes differentially expressed based on RTPCR analysis.

| S. No | Gene | Gene Family | 12hpa | 24hpa | 2dpa | 3dpa | 7dpa |
| --- | --- | --- | --- | --- | --- | --- | --- |
| 1 | papss2a | NGS validation | -4.8 | -1.9 | -2.3 | -1.9 | -3.1 |
| 2 | adam8a |  | 0.7 | 5.2 | 5.8 | 7.6 | 3.2 |
| 3 | adm2a |  | -3.4 | 1.5 | 1.3 | -0.6 | -2.7 |
| 4 | alox12 |  | -1.3 | 1.2 | 1.1 | 1.2 | 0.2 |
| 5 | bhlha9 |  | 0 | 2 | 1.9 | 1.5 | -2 |
| 6 | bhlhe41 |  | -3 | -0.6 | -1 | -1.7 | -1.7 |
| 7 | bco2a |  | -2.7 | -0.7 | 0.1 | -0.4 | -1.3 |
| 8 | bmb |  | -2.3 | 2.8 | 2.8 | 4.1 | 1 |
| 9 | cav2 |  | -2 | 1 | 1.4 | 0.8 | -0.1 |
| 10 | caprin1b |  | -2.2 | 0.2 | 0.3 | 0.8 | -2.6 |
| 11 | cfd |  | -2.8 | -0.3 | -1 | 0.5 | -1 |
| 12 | cycsb |  | 1.5 | 3.2 | 3.3 | 4.2 | 1.8 |
| 13 | dbpb |  | -3.2 | 0.4 | 0.1 | -2.5 | -1.7 |
| 14 | defbl1 |  | -7.4 | 1.6 | 1.2 | -1.8 | -5.5 |
| 15 | deptor |  | -4 | -1.3 | -0.3 | -1.8 | -2.9 |
| 16 | daam1b |  | 1.5 | 4.2 | 4 | 5.2 | 2.3 |
| 17 | dnmt3 |  | -4.6 | -2.4 | 0.4 | 4.7 | 0.4 |
| 18 | dut |  | -2.5 | 3.1 | 2.9 | 3.3 | -0.4 |
| 19 | ednrb1a |  | -5.2 | 0.9 | 1.2 | -1.6 | -2.9 |
| 20 | eif4ebp3 |  | -3.8 | -3.8 | -2.9 | -1.9 | -3.2 |
| 21 | fam102ba |  | -1.8 | -0.6 | -0.7 | 0.1 | -0.8 |
| 22 | fgf20a |  | -1.7 | 1.1 | 1.3 | 1.5 | -3.1 |
| 23 | fn1b |  | 3.9 | 7.9 | 8 | 9.3 | 5.3 |
| 24 | fn1b |  | 2.3 | 5.6 | 5.1 | 6.9 | 3.9 |
| 25 | fhl2b |  | -2.8 | -0.3 | -0.2 | -1.5 | -0.7 |
| 26 | gins2 |  | 0.9 | 4.3 | 4.4 | 4.2 | 1.7 |
| 27 | gsdf |  | -4.7 | 0.6 | 0.2 | -0.7 | -2.1 |
| 28 | gtpbp4 |  | -0.4 | 1.1 | 1.4 | 1.7 | 0.4 |
| 29 | has1 |  | -3.8 | 1.9 | 1.3 | 0.3 | -3 |
| 30 | hsd17b7 |  | -0.3 | 2.1 | 2.2 | 2.1 | 1.7 |
| 31 | inhbaa |  | -0.3 | 2.5 | 2.6 | 2.2 | 0.3 |
| 32 | il1b |  | -0.3 | 2.4 | 1.4 | 2.3 | 0.4 |
| 33 | il2rb |  | -5.8 | 0.6 | 0.3 | -1.9 | -3.9 |
| 34 | il7r |  | -3.8 | 0.8 | 1.4 | 1.8 | 1.4 |
| 35 | idh2 |  | -0.3 | 2.4 | 2.4 | 3.2 | 1.2 |
| 36 | krt18 |  | 4.3 | 5.9 | 6.6 | 7.4 | 2.6 |
| 37 | lepb |  | 4.5 | 3.9 | 6.2 | 8.3 | 7.7 |
| 38 | mmp13a |  | 4.2 | 1.6 | 2.4 | 6.7 | 5.2 |
| 39 | mmp9 |  | -3.4 | 1.6 | 1 | -0.2 | -1.7 |
| 40 | mdka |  | 0.6 | 4 | 4 | 6 | 3.4 |
| 41 | nos1 |  | -1.8 | -1 | -0.2 | -0.8 | -1.2 |
| 42 | orc3 |  | 1.8 | 5.3 | 4.7 | 5.3 | 2.9 |
| 43 | orc6 |  | 0.4 | 4.5 | 4.5 | 5 | 2.8 |
| 44 | pax7b |  | -4.4 | -1.1 | -1.9 | -2.8 | -3.8 |
| 45 | pim1 |  | -3.5 | -1.8 | -0.3 | -0.1 | -2.6 |
| 46 | pabpc4 |  | -0.9 | 1.3 | 1.2 | 1.7 | -0.6 |
| 47 | kcnj1a |  | -4.4 | -2.8 | -2 | -1.6 | -4.3 |
| 48 | ptgdsb |  | -3.1 | -2.5 | -2.1 | -2.7 | -1.9 |
| 49 | pmepa1 |  | 1 | 1.7 | 1.9 | 0.9 | 0.3 |
| 50 | psmb7 |  | 1.7 | 3.5 | 3.5 | 4.6 | 2.7 |
| 51 | ppp1r3cb |  | -4.9 | -1 | -1.3 | -2.2 | -3.6 |
| 52 | rasl11b |  | -2.1 | 1.3 | 1.3 | 0.3 | -1.2 |
| 53 | rgs5a |  | -2.3 | 0.7 | 0.4 | -0.3 | -0.9 |
| 54 | rgs4 |  | -4.6 | 0.7 | 0.5 | -1.9 | -2.6 |
| 55 | rxfp2b |  | -0.6 | 0.8 | 1.3 | 1.1 | 0.4 |
| 56 | LOC100150965 |  | -3.3 | -0.4 | -0.1 | -1.6 | -2 |
| 57 | rhobtb2a |  | -2.1 | 0.7 | 0.9 | 0.3 | -1 |
| 58 | scpp8 |  | -2.6 | 1.9 | 1 | -0.1 | -2.2 |
| 59 | si:ch211-122l24 |  | -2.3 | 2.1 | 1.5 | 0.9 | -1.9 |
| 60 | si:ch211-126j24 |  | 4.7 | 3.4 | 4.7 | 4.9 | 1.2 |
| 61 | si:busm1-160c18.3 |  | -5 | -3 | -1.2 | -0.2 | -3.8 |
| 62 | si:dkey-51e6 |  | -3.1 | -1.4 | -1.4 | -0.6 | -2.3 |
| 63 | si:dkeyp-113d7 |  | -2.6 | -0.1 | -1.2 | -1.2 | -1.5 |
| 64 | smtnl |  | -2.8 | 1.1 | 1.4 | -0.8 | -1.1 |
| 65 | slc3a2b |  | -1.5 | 1.3 | 0.9 | 1.4 | -0.8 |
| 66 | slc6a6b |  | -0.5 | 1 | 1 | 0.8 | 0.2 |
| 67 | slc6a6b |  | -3.3 | 1.3 | 1 | -1.3 | -2 |
| 68 | sat1a |  | 0.2 | 1.5 | 3 | 3.5 | 2.2 |
| 69 | sox11a |  | -4.5 | 1.2 | 1.3 | 0.2 | -2.2 |
| 70 | scdb |  | -5.5 | -1.4 | 0.8 | -1.5 | -3.6 |
| 71 | sult1st6 |  | -1.5 | -1.1 | 0.8 | -0.4 | -0.2 |
| 72 | socs3b |  | -1.7 | 1.9 | 2.5 | 2.8 | -0.1 |
| 73 | syt4 |  | -5.4 | -2.1 | -0.7 | -2.9 | -3.8 |
| 74 | txn |  | 1 | 4.2 | 4.4 | 5.5 | 2.1 |
| 75 | txndc12 |  | 1.4 | 4 | 4.5 | 5.8 | 3.2 |
| 76 | timp2b |  | -5.2 | -0.7 | 0.6 | -1.8 | -4 |
| 77 | tcea3 |  | -3 | 2.1 | 2 | 2.4 | 1 |
| 78 | tpm4a |  | -1.3 | 1.6 | 1.7 | 1.2 | -0.7 |
| 79 | tuba1b |  | -1.9 | 2.2 | 2.3 | 2.5 | -0.2 |
| 80 | zgc:103599 |  | 2.6 | 4.5 | 4.5 | 5.9 | 3.6 |
| 81 | zgc:123339 |  | -4.1 | 0.9 | 0.2 | -2.9 | -2.6 |
| 82 | zgc:136474 |  | -2.7 | 0.5 | 0.7 | 0.5 | -0.4 |
| 83 | zgc:136902 |  | -2.4 | 1.9 | 2.7 | 1.8 | -1.4 |
| 84 | zgc:136930 |  | -1.4 | -0.1 | 0.3 | 0.5 | 0.5 |
| 85 | zgc:153031 |  | -0.1 | -0.6 | -0.3 | 0.5 | 0 |
| 86 | zgc:153154 |  | -2.9 | -1.4 | -1.8 | -2.1 | -2 |
| 87 | zgc:153629 |  | 2.1 | 6.3 | 6.2 | 6.8 | 2.5 |
| 88 | zgc:153920 |  | -5.3 | 1.1 | 1.4 | -0.9 | -3.3 |
| 89 | zgc:154093 |  | -1 | 0.8 | 0.8 | 0.2 | -1.5 |
| 90 | zgc:162356 |  | -1.4 | 0 | 1 | 2.1 | -0.2 |
| 91 | zgc:163079 |  | -3.9 | -1.1 | -1.5 | -2.3 | -3.4 |
| 92 | zgc:174917 |  | -4.8 | 2.9 | 2.5 | 1.4 | -2.5 |
| 93 | zgc:194409 |  | -2.2 | 2.1 | 2.6 | 1.8 | 0.3 |
| 94 | zgc:194993 |  | -1.6 | 0.3 | 0.2 | 0.2 | 0.4 |
| 95 | zgc:73226 |  | -4 | 1.3 | 1.1 | -1.2 | -3.2 |
| 96 | zgc:77358 |  | -3.5 | 1.1 | 1.4 | 2.6 | -0.9 |
| 97 | zbtb16a |  | -4.5 | -0.2 | 0.1 | -1.9 | -2.8 |
| 98 | si:ch1073-220m6.1 |  | -1.5 | -2.4 | -0.1 | -1.4 | -1.5 |
| 99 | H2 Ab1 |  | -2 | 1.9 | 1.4 | -0.7 | -1.5 |
| 100 | Btla |  | -5.3 | -2.1 | -1.5 | -4.3 | -3.9 |
| 101 | MUC22 |  | -1.5 | -0.4 | 1.1 | -2.4 | -2.9 |
| 102 | zbtb16a |  | -3 | 0.1 | 0.1 | -2.2 | -2.3 |
| 103 | zmy4 |  | -2.6 | 0.5 | 0 | -0.5 | -2.8 |
| 104 | cypCar_00031856 |  | -2.9 | -0.2 | 0 | -1.3 | -1.6 |
| 105 | capn2 |  | -4 | 0.2 | 0.6 | -1.9 | -3.1 |
| 106 | CSPG4 |  | -3.2 | 0.9 | 0.2 | -2.9 | -3.1 |
| 107 | [efemp1](https://www.google.co.in/url?sa=t&rct=j&q=&esrc=s&source=web&cd=1&cad=rja&uact=8&ved=0ahUKEwjm1KTL5t7YAhUMT7wKHQGyDq0QFggmMAA&url=http%3A%2F%2Fwww.uniprot.org%2Funiprot%2FQ5PR37&usg=AOvVaw14PUO3ZkXJgVNbWO0GNf2v) |  | -2.5 | -0.7 | -0.4 | -2.5 | -2.3 |
| 108 | CH211-197O22 |  | -3.4 | -0.4 | 0 | -3.3 | -4.2 |
| 109 | DKEY-246P9 |  | -6.2 | -0.7 | -2.2 | -4.9 | -4 |
| 110 | SLC7A1 |  | -1.9 | -0.3 | -0.4 | -2.8 | -1.5 |
| 111 | DKEY-288C23 |  | -10 | -2.2 | -3.1 | -5.9 | -7.7 |
| 112 | [ft2](https://www.ncbi.nlm.nih.gov/gene/111265018) |  | -2 | -0.2 | -0.2 | -2.5 | -3.2 |
| 113 | [clec19a](https://www.ncbi.nlm.nih.gov/gene/100491258) |  | -2.7 | 0.1 | 0.2 | -1.9 | -2.7 |
| 114 | USP2 |  | -2.3 | -0.1 | -0.7 | -0.8 | -2.1 |
| 115 | si:ch211-252f13.5 |  | -2.7 | 0.7 | 0.6 | -0.4 | 0.2 |
| 116 | DKEY-246P9 |  | -3.3 | 0.6 | 0.7 | -0.5 | -1.3 |
| 117 | [PCDH1](https://www.ncbi.nlm.nih.gov/gene/416194) |  | -3.2 | -0.2 | -0.2 | -1.8 | -2.1 |
| 118 | CH211-76L23 |  | -2.1 | 1.1 | 0.7 | -1.3 | -2 |
| 119 | hsd17b12a |  | -1.9 | 0.4 | -0.1 | -0.5 | -1 |
| 120 | DAN4 |  | -3.1 | 0.7 | 0.7 | -0.6 | -1.3 |
| 121 | LOC107729934 |  | -1.7 | 0.7 | 0.9 | -0.6 | -1.2 |
| 122 | COL9A3 |  | -4.8 | 0.7 | -0.3 | -1.9 | -3.5 |
| 123 | DDB_G0282133 |  | -3.4 | 0.1 | 0.3 | -0.8 | -1.2 |
| 124 | PRSS3 |  | -1.8 | 0.5 | -0.3 | -1.3 | -1.9 |
| 125 | SLK |  | -3.2 | 1.6 | -0.3 | -0.4 | -2 |
| 126 | LOC110438252 |  | -5.5 | 1.8 | 0.4 | -2.8 | -4.1 |
| 127 | N/A |  | -0.9 | -0.7 | -0.1 | -0.5 | -1.7 |
| 128 | wu:fc46h12 |  | -0.6 | 1 | 1.9 | -0.5 | 1.2 |
| 129 | si:dkey-151j17.4 |  | -1.8 | 0.2 | 0.5 | -1.5 | -1.7 |
| 130 | synm |  | -4.1 | -0.3 | -0.4 | -3.2 | -3.3 |
| 131 | traf7l |  | -4.2 | 1 | 0.2 | -2.4 | -2.2 |
| 132 | ADGRF3 |  | -1.3 | -0.4 | 0.5 | -0.3 | 0.1 |
| 133 | sarg |  | -2.7 | 0.7 | -0.2 | -1 | -1.8 |
| 134 | LOC100330916 |  | -2.7 | -1.6 | -1.6 | -2.2 | -1.8 |
| 135 | si:ch211-156b7.4 |  | -3.3 | 1.5 | 1.4 | 0.2 | -1.9 |
| 136 | plxn a2 |  | -4 | 0.8 | -0.4 | -0.6 | -2.3 |
| 137 | CYGB1 |  | -1.4 | 2 | 3.1 | 2.7 | 0.5 |
| 138 | GEN |  | -3.4 | 0.8 | 0.2 | -1 | -2.3 |
| 139 | csp2 |  | -2.7 | 1.8 | 2.5 | 2.4 | 0 |
| 140 | si:ch211-153b23.3 |  | -3.7 | 0.8 | 1.4 | -1 | -1.1 |
| 141 | IL2RGA | Interleukin genes | -3.5 | -4.6 | -4.8 | -2.9 | -4 |
| 142 | NFIL3 |  | -2.9 | -3 | -3.3 | -2.7 | -3.4 |
| 143 | IL4 |  | -1.6 | -1.9 | -3.4 | -1.6 | -2.4 |
| 144 | IL4R.2 |  | -4.3 | -5.8 | -4.5 | -2.8 | -4.5 |
| 145 | IL - 6 (IL6ST) |  | -2.3 | -3.8 | -3.2 | -2.5 | -2.9 |
| 146 | IL - 7 (IL6) |  | -1 | -1.2 | 0.4 | -2.7 | -1.3 |
| 147 | IL - 8 (IL7) |  | -4.6 | -4.7 | -5.8 | -4.7 | -4.4 |
| 148 | IL - 9 (IL10) |  | -3.4 | -4.1 | -2.5 | -3.8 | -4 |
| 149 | IL - 10 (IL12A) |  | -3 | -3.1 | -2.3 | -2.5 | -4.2 |
| 150 | IL - 11 (IL12BA) |  | -1.6 | -0.8 | -2.2 | -1.9 | -2 |
| 151 | IL - 12 (IL12RB2) |  | -4.7 | -3.5 | -4.7 | -3.1 | -2.9 |
| 152 | IL - 13 (IL13RA1) |  | -2.6 | -4.4 | -5.2 | -3.8 | -3.3 |
| 153 | IL - 14 (IL13RA2) |  | -2.2 | -3.8 | -5.5 | -4.5 | -3.2 |
| 154 | IL - 15 (IL15) |  | -5.8 | -5.2 | -6.2 | -4 | -2.6 |
| 155 | IL - 16 (IL16) |  | -5.6 | -4.7 | -4 | -3.2 | -2.8 |
| 156 | IL - 17 (IL17A/F1) |  | -1.1 | -2.6 | -3.8 | -2.7 | -0.9 |
| 157 | IL - 18 (IL17A/F3) |  | -2 | -3.5 | -3.7 | -2.8 | 0.7 |
| 158 | IL - 19 (IL17C) |  | 0.4 | 0.6 | 0.1 | -0.2 | 2.8 |
| 159 | IL - 20 (IL17D) |  | -2.5 | -2.9 | -3 | -3 | -2.8 |
| 160 | IL - 21 (IL20RA) |  | -3.3 | -4.5 | -5.3 | -4.3 | -4 |
| 161 | IL - 22 (IL22) |  | -1.7 | -2.8 | -3.1 | -2.9 | -3.3 |
| 162 | IL - 23 (IL34) |  | -2.6 | -4.1 | -4.3 | -3.4 | -3 |
| 163 | IL - 24 (OSMR) |  | 1 | 1 | 1.6 | 1.5 | 4.2 |
| 164 | IL - 25 (LIFRA) |  | -4.1 | -4.9 | -5.9 | -4.5 | -4.6 |
| 165 | IL - 26 (CNTFR) |  | -3.9 | -4.4 | -3.1 | -2 | -4.1 |
| 166 | IL - 27 (CRFB16) |  | -5 | -5.2 | -6.1 | -6.1 | -4.3 |
| 167 | IL - 28 (IL1β) |  | -1.1 | -1.5 | -4 | -1.3 | -2.2 |
| 168 | IL - 29 (IL1RAP) |  | -3 | -4.1 | -4.9 | -4.1 | -3.5 |
| 169 | SLC17A9B | Solute Career gene | -1.02 | 2.18 | 2.38 | 2.02 | 2.11 |
| 170 | SLC1A4 |  | 0.55 | 5.19 | 5.99 | 5.54 | 5.77 |
| 171 | SLC23A1 |  | 0.17 | 3.52 | 3.4 | 1.96 | 2.19 |
| 172 | SLC24A5 |  | -2.34 | -1.18 | -1.56 | -0.73 | 0.33 |
| 173 | SLC25A11 |  | -1.56 | 2.23 | 3.02 | 2.57 | 3.09 |
| 174 | SLC25A1A |  | -0.73 | 3.09 | 3.57 | 2.41 | 3.29 |
| 175 | SLC25A25B |  | -1.04 | 1.21 | 3.01 | 1.93 | 2.49 |
| 176 | SLC25A32B |  | -2.11 | 1.76 | 1.85 | 0.65 | 1.54 |
| 177 | SLC30A1A |  | -1.51 | 1.71 | 2.03 | 1.06 | 1.59 |
| 178 | SLC34A2B |  | -1.22 | 1.5 | 1.95 | 0.68 | 1.16 |
| 179 | SLC35E3 |  | -1.44 | 1.72 | 2.72 | 1.69 | 1.85 |
| 180 | SLCO2B1 |  | -2.62 | 1.11 | 1.71 | 1.17 | 1.52 |
| 181 | SLC7A7 |  | -5.15 | -3.16 | -4.38 | -4.29 | -3.87 |
| 182 | SLC38A4 |  | -1.77 | 1.57 | 2.27 | 1.71 | 1.38 |
| 183 | SLC38A5B |  | -4.62 | -2.55 | -2.68 | -3.09 | -1.71 |
| 184 | SLC3A2B |  | -1.51 | 1.45 | 1.29 | 0.8 | 0.91 |
| 185 | PRMT1 | Protein arginine methyltransferases | 0.3 | 0.8 | 1 | 1.3 | 1.5 |
| 186 | PRMT2 |  | -1 | 0.8 | 0.6 | -0.8 | 0.9 |
| 187 | PRMT3 |  | 0.3 | 0 | -0.6 | -0.7 | 0 |
| 188 | PRMT4 |  | -0.7 | -0.2 | -0.1 | -0.7 | 0.3 |
| 189 | PRMT5 |  | 0.7 | 1.1 | 1 | 1.1 | 1.5 |
| 190 | PRMT6 |  | -1.1 | -0.2 | 0.6 | -1.9 | -0.6 |
| 191 | PRMT7 |  | -2 | -0.7 | -1 | -1.3 | -0.5 |
| 192 | PRMT8 |  | 0.4 | 0.9 | 2.2 | 1.1 | 2.2 |
| 193 | PMRT9 |  | -1.9 | -0.2 | 1.8 | -0.3 | 0.2 |
| 194 | HOXA1A | Homeobox genes | -0.2 | 6.9 | 6.7 | 5.3 | 4.8 |
| 195 | HOXA2B |  | 0.4 | 7.1 | 7.2 | 5.5 | 4.4 |
| 196 | HOXA5A |  | 0.2 | 6.6 | 6.5 | 5.5 | 4.7 |
| 197 | HOXA9A |  | -0.3 | 6.1 | 5.9 | 4.9 | 4 |
| 198 | HOXA10B |  | -0.3 | 6.8 | 6.2 | 4.9 | 4.6 |
| 199 | HOXA11A |  | -0.3 | 6.4 | 6.2 | 4.1 | 4.3 |
| 200 | HOXA13B |  | -0.6 | 5 | 4.8 | 4.1 | 2.1 |
| 201 | HOXB2A |  | 1.3 | 8.2 | 7.4 | 6.6 | 5.5 |
| 202 | HOXB9A |  | 0.3 | 6.6 | 6.5 | 5.2 | 4.7 |
| 203 | HOXC5A |  | 0.4 | 5.3 | 5.9 | 5.2 | 4.1 |
| 204 | HOXC11A |  | -0.1 | 5.9 | 5.3 | 5.6 | 4.4 |
| 205 | HOXC13B |  | 0.8 | 7.1 | 6.9 | 5.4 | 5.5 |
| 206 | HOXD10A |  | 0.5 | 7.3 | 5.9 | 5.7 | 5.2 |
| 207 | Dbh | Neurotransmitter genes | -1.3 | 1.1 | 0.6 | -0.2 | -0.5 |
| 208 | Ddc |  | 1.1 | 3.9 | 1.4 | 2.2 | 3.6 |
| 209 | qdprb1 |  | -0.6 | 1.5 | 1.6 | 1.1 | 2.5 |
| 210 | slc18a2 |  | -1 | 2 | 3 | 1.6 | 3.5 |
| 211 | Chata |  | -0.2 | 3.6 | 3.7 | 2.2 | 4.4 |
| 212 | chrna7 |  | 1.2 | 1.9 | 2.2 | 0.9 | 1.6 |
| 213 | chrna4b |  | 0 | 2.7 | 4.1 | 2.7 | 3.7 |
| 214 | chrm3a |  | -0.4 | 2.7 | 3.2 | 1.4 | 3.9 |
| 215 | Ache |  | -0.5 | 0.5 | 0.4 | 0.9 | 0.9 |
| 216 | cht1 |  | -0.5 | 2.7 | 3.4 | 2.3 | 3.8 |
| 217 | slc6a1b |  | 0.4 | 2.6 | 3.4 | 1.9 | 2.1 |
| 218 | tph1a |  | -0.6 | 1.1 | 1.1 | 1 | 3.2 |
| 219 | tph1b |  | 0.6 | 3.3 | 3.9 | 3.6 | 4.8 |
| 220 | slc6a4b |  | 0.1 | 2.5 | 2.8 | 1.5 | 0 |
| 221 | adra2c |  | -1 | 1.4 | 3.1 | 0.7 | 3.5 |
| 222 | htr1ab |  | 0.4 | 3.5 | 5.6 | 4.8 | 7.5 |
| 223 | slc17a8 |  | 0.9 | 3.3 | 1.5 | 3.3 | 5.4 |
| 224 | Mao |  | -0.9 | 2.2 | 3.3 | 1.8 | 3 |
| 225 | slc32a1 |  | 0.4 | 2.6 | 4.7 | 2.7 | 4.3 |
| 226 | hdc |  | 0 | 2.6 | 3.5 | 2.6 | 3.9 |
| 227 | hrh3 |  | -0.3 | 2.7 | 4.2 | 3.7 | 3.9 |
